# Dissecting rice nonhost resistance to the wheat blast fungus

**DOI:** 10.64898/2026.11.14.766612

**Authors:** Motoki Shimizu, Yukie Hiraka, Soichiro Asuke, Hiroaki Adachi, Sayaka Yuki, Eiko Kanzaki, Kazue Ito, Akiko Hirabuchi, Naomi Miyaji, Takuma Kajino, Koki Minoji, Mina Ohtsu, Toshiyuki Sakai, Mark J Banfield, Akira Abe, Yukio Tosa, Sophien Kamoun, Ryohei Terauchi

**Author notes:** Corresponding author &.

## Abstract

The pandemic wheat blast fungus *Magnaporthe oryzae* destroys wheat but cannot infect rice, a form of resistance so uniform that it does not segregate in genetic crosses and cannot be mapped. We broke this nonhost resistance into single-gene phenotypes by expressing individual effectors from the wheat-infecting pathotype, one at a time, in a rice-infecting isolate. Screening 35 effectors across 20 rice cultivars (700 combinations) identified six that trigger resistance. One, eTO5, is recognized by *Pi-eTO5*, a genetically linked quartet of NLR immune receptors: SQ1, SQ2, and SQ4 are required for resistance, whereas SQ3 prevents constitutive defense activation dependent on the other three. eTO5 is conserved across the pandemic B71 lineage spanning South America, Asia and Africa. Effector transfer into a compatible pathogen should make nonhost resistance genetically tractable in other host–pathogen systems.

**One-Sentence Summary:** One-at-a-time effector transfer turns a nonhost barrier that cannot be genetically mapped into single-gene resistance phenotypes, yielding known and novel rice immune receptors.

## Main text

Plants detect invading pathogens through immune receptors, and many resistance (*R*) genes deployed in crops encode intracellular receptors of the nucleotide-binding leucine-rich repeat (NLR) class (1). Most cloned *R* genes have been identified from crop germplasm or its wild relatives, where resistance segregates and can therefore be genetically mapped. Nonhost species form a different kind of *R* gene reservoir. Their NLR receptors have not been under selection from coevolution with the pathogen in question, and resistance transferred from a nonhost species to a crop may therefore provide a distinct and potentially durable source of resistance (2). However, these genes are difficult to access: nonhost resistance is often uniform across a species, does not segregate in crosses, and cannot be genetically mapped. In addition, which pathogen effectors with an avirulence activity (AVR) are detected by nonhost *R* genes remains poorly known.

The blast fungus *Magnaporthe* (Syn. *Pyricularia*) *oryzae* causes disease on dozens of wild and cultivated grasses and is differentiated into host-associated genetic lineages, with occasional gene flow between them (3, 4). Wheat blast was first reported in Paraná, Brazil, in 1985 and subsequently spread across South America. In 2016 and 2018, the disease emerged in Bangladesh and Zambia, respectively, following separate introductions of the pandemic B71 clonal lineage from South America (5–9). Only a limited number of wheat resistance genes effective against wheat blast have been described (10, 11). Adaptation of *M. oryzae* to wheat has also involved loss of AVR effectors: functional loss of *PWT3*, for example, enabled escape from *Rwt3*-mediated resistance (12). Rice, in contrast, is uniformly highly resistant to the wheat-infecting pathotype.

We sought a way to genetically dissect rice nonhost resistance to the wheat-infecting pathotype of *M. oryzae*. We reasoned that the extensive NLR diversity of rice germplasm could include receptors capable of detecting wheat blast effectors. Because resistance to the intact pathogen is likely to involve multiple recognition events, we screened rice with effector genes rather than with the pathogen itself. Individual candidate effectors from the wheat-infecting pathotype were expressed, one at a time, in a rice-infecting isolate that is otherwise fully virulent on rice. In this way, each transformant isolates one component of the nonhost barrier as a genetically tractable resistance phenotype. The screen recovered effector recognition mediated by *Pii* and *Pia*, resistance loci previously characterized against the rice-infecting pathotype, showing that the barrier to the wheat-infecting pathotype rests in part on the resistance repertoire rice already maintains against its own pathogen. It also identified *eTO5*, an effector without a detectable orthologue in the rice-infecting pathotype, as recognized by a subset of cultivars. We then used *eTO5* to genetically isolate its cognate resistance locus, *Pi-eTO5*, and found that *eTO5* is conserved across the pandemic B71 lineage of the wheat blast fungus.

### Host specificity between rice and wheat blast enables systematic discovery of AVR effectors

To examine host specificity, we inoculated leaves of three rice cultivars, Hitomebore, Sasanishiki and Nipponbare, with the rice-infecting *M. oryzae* pathotype *Oryza* (MO) isolate Ao92-06-2 and the wheat-infecting pathotype *Triticum* (MT) isolate Br48. The MO isolate Ao92-06-2 produced expanding lesions on all three rice cultivars, whereas the MT isolate Br48 failed to infect any of them. These results confirmed the contrasting host specificity of the two pathotypes on rice (**Fig**. **1A****; Fig. S1**).

**Fig. 1.**
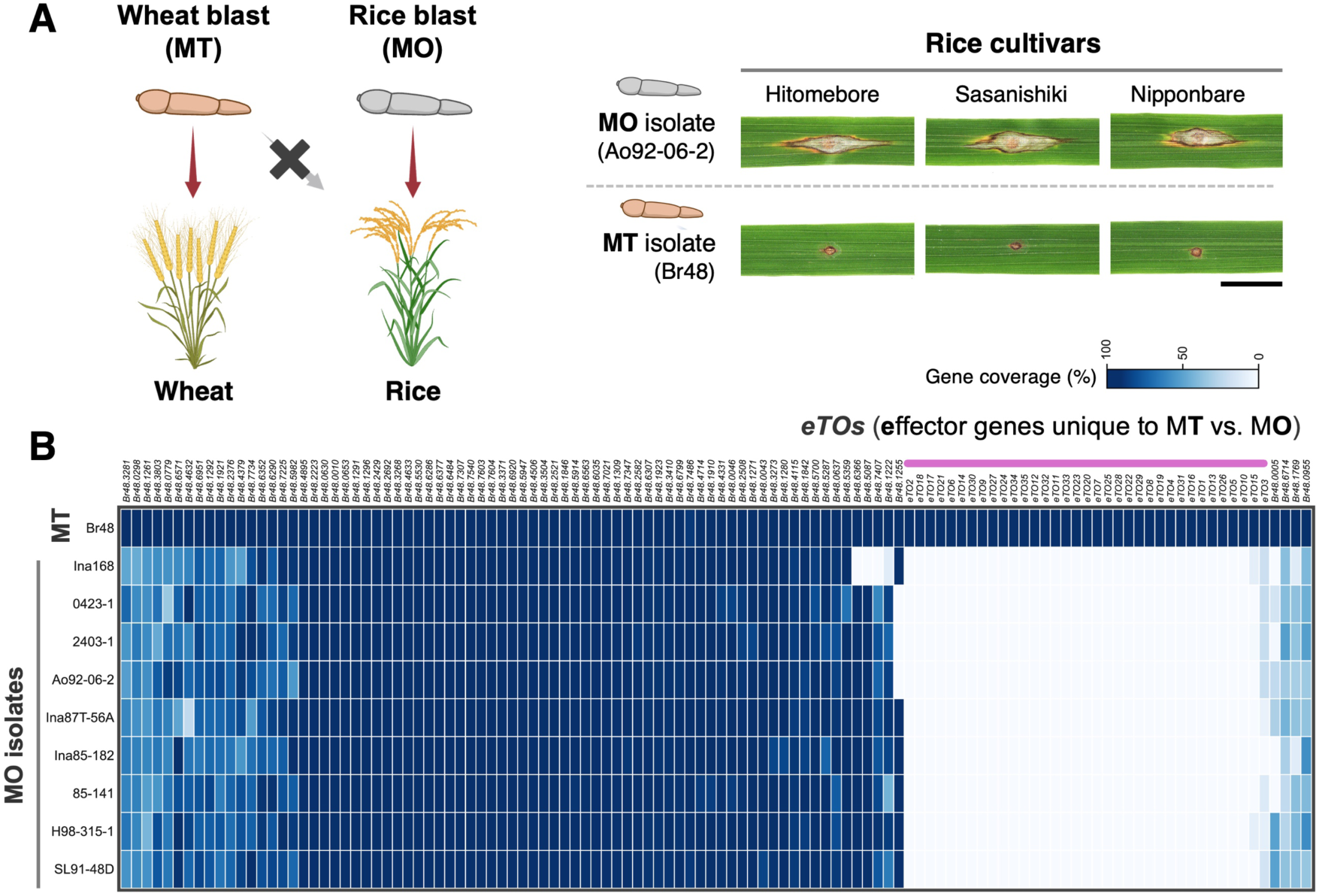
Host specificity between rice- and wheat-infecting *M. oryzae* pathogens enables systematic identification of MT-specific candidate effectors. (A) Host-specificity of *Magnaporthe oryzae* pathotype *Oryza* (MO) and pathotype *Triticum* (MT). The rice-infecting MO isolate Ao92-06-2 produced expanding lesions on three rice cultivars (Hitomebore, Sasanishiki, and Nipponbare), whereas the wheat-infecting MT isolate Br48 failed to produce expanding lesions on any of them. Representative images from punch-inoculation assays at 7 days post-inoculation (dpi) are shown. Scale bar, 1cm. **(B)** Comparative genomic analysis of candidate cytoplasmic effector genes in MT and MO isolates. The heatmap shows the gene coverage for 114 candidate effector genes identified in the MT isolate Br48 across Br48 and nine MO isolates. Candidates were selected based on four criteria: a predicted signal peptide, expression of TPM > 100 at 24 hours post-inoculation, classification as a cytoplasmic or cytoplasmic/apoplastic effector by EffectorP, and a predictied protein length of <150 amino acids. Thirty-five genes present in Br48 but with ≤ 30% gene coverage in each of the nine MO isolates were operationally defined as absent from MO and designated *eTOs,* for effector genes unique to MT versus MO. Gene order along the horizontal axis and isolate order along the vertical axis were determined by hierarchical clustering using Euclidean distance and average linkage.

We reasoned that this host specificity could reflect the collective recognition of MT-specific effectors as avirulence (AVR) factors by rice immune receptors and could therefore be exploited as a discovery platform to systematically identify AVR genes. We first sought to define the repertoire of candidate effector genes associated with the wheat blast pathogen.

We generated a *de novo* genome assembly of Br48 (**Data S1**) and performed RNA sequencing during barley infection to identify genes expressed *in planta*. We identified 114 candidate cytoplasmic effector genes that met four criteria: they encoded predicted secreted proteins, were highly expressed at 24 hours post inoculation (TPM>100), were classified as cytoplasmic or cytoplasmic/apoplastic effectors by EffectorP (13), and encoded proteins shorter than 150 amino acids **(Fig. S2**).

We then assessed the presence of these genes in nine MO isolates (**Data S2**). Thirty-five of the 114 candidates were present in Br48 but showed minimal sequence coverage (≤30%) in all nine tested MO isolates and were therefore operationally defined as absent from MO (**Fig. 1B; Fig. S3**). We designated these MT-specific genes *eTOs*, for effectors unique to MT versus MO (**Data S3**).

### Functional screen of MT-specific effectors reveals multiple AVR activities in rice

To functionally identify MT-specific effector genes (*eTO*s) that trigger resistance in rice, we developed a systematic functional screening pipeline (**Fig. S4**). We generated a library of transgenic *M. oryzae* Ao92-06-2 strains, each expressing one of the 35 *eTO* genes under the control of the *pex22* (*AVR-Pia*) promoter, which drives effector expression during host infection (14).

Each transformant was inoculated onto 20 rice cultivars that were susceptible to the parental Ao92-06-2 isolate (**Data S4**). Resistance specifically induced by an *eTO*-expressing transformant was interpreted as AVR activity recognized by the corresponding rice cultivar.

Screening all 35 transformants across the 20 rice cultivars generated 700 host-pathogen combinations. Six *eTOs* (*eTO1*, *eTO2*, *eTO5*, *eTO16*, *eTO24*, and *eTO31*) triggered resistance in at least one cultivar (**Fig. 2; Fig. S5; Fig. S6**). Notably, *eTO2* triggered resistance in all 20 rice cultivars tested, showing that it was broadly recognized across the panel and identifying it as a candidate determinant of the inability of MT to infect rice.

**Fig. 2.**
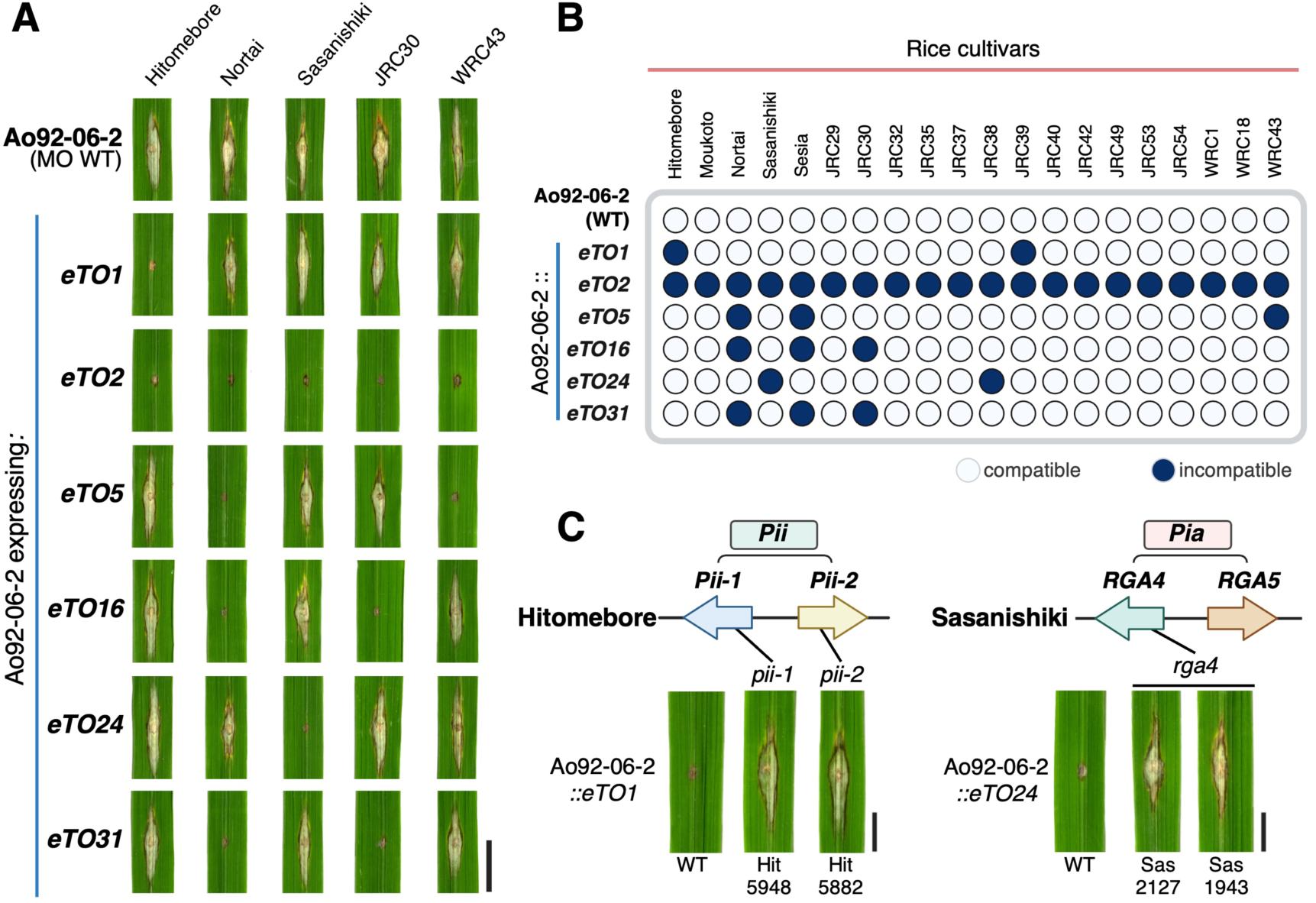
Functional screen of MT-specific effector genes identifies multiple AVR activities in rice. **(A)** Representative punch-inoculation phenotypes of five rice cultivars inoculated with the parental MO isolate Ao92-06-2 or Ao92-06-2 transformants expressing individual MT-specific *eTO* genes under the control of the *pex22* promoter from *AVR-Pia*. Leaves were photographed at 10 days post-inoculation (dpi). Six *eTOs* (*eTO1, eTO2, eTO5, eTO16, eTO24*, and *eTO31*) triggered resistance in at least one rice cultivar. Quantitative lesion area measurements are shown in **Fig. S6**. Scale bar, 1 cm. **(B)** Compatibility matrix showing the responses of 20 rice cultivars to the parental Ao92-06-2 isolate and the six resistance-inducing *eTO*-expressing transformants. Filled blue circles indicate incompatible interactions (resistance), whereas open circles indicate compatible interactions (susceptibility). *eTO2* triggered resistance in all 20 cultivars tested, whereas the other five *eTOs* showed cultivar-specific recognition patterns. **(C)** Genetic validation of *eTO1-* and *eTO24*-triggered resistance. Hitomebore was resistant to Ao92-06-2 expressing *eTO1*, whereas two independent *pii* mutant lines, Hit5948 and Hit5882, were susceptible, demonstrating that *eTO1*-triggered resistance requires *Pii*. Similarly, Sasanishiki was resistant to Ao92-06-2 expressing *eTO24*, whereas two independent *rga4* mutant lines, Sas2127 and Sas1943, were susceptible, demonstrating that *eTO24*-triggered resistance requires *Pia*. Scale bars, 0.5 cm.

The screen independently recovered two previously characterized AVR–R gene relationships without prior knowledge of their identities. eTO1 and eTO24 showed amino acid sequence similarity to the previously characterized MO AVR effectors AVR-Pii and AVR1-CO39, respectively (**Fig. S7**), which are recognized by the rice resistance loci *Pii* (15) and *Pia* (16). To test whether *eTO1*-triggered resistance requires *Pii*, we inoculated Hitomebore and two independent *pii* mutant lines, Hit5948 and Hit5882, with Ao92-06-2 expressing *eTO1.* Hitomebore was resistant, whereas both *pii* mutants were susceptible (**Fig. 2C; Fig. S8A**), demonstrating that *eTO1*-triggered resistance requires *Pii*. Similarly, Sasanishiki was resistant to Ao92-06-2 expressing *eTO24,* whereas two independent *rga4* mutant lines, Sas2127 and Sas1943, were susceptible (**Fig. 2C; Fig. S8B**), demonstrating that *eTO24*-triggered resistance requires *Pia.* These genetic results establish that the screen recovers genetically defined AVR–R relationships. They also show that two of the six recognition events are mediated by *R* genes that rice maintains against the rice-infecting pathotype, indicating that the barrier to MT is assembled in part from the standing anti-MO resistance repertoire rather than from machinery dedicated to nonhost recognition. Because *Pii* and *Pia* segregate among cultivars, however, neither can by itself account for resistance that is uniform across the species. We therefore focused on *eTO5*, whose recognition is not attributable to a known rice blast resistance gene, to identify its cognate locus.

### *Pi-eTO5* comprises a genetically linked NLR quartet

Using the pipeline shown in **Fig. S4B**, we sought to identify the rice resistance locus responsible for *eTO5*-triggered resistance, hereafter designated *Pi-eTO5* (**Fig. 3**). We crossed the three resistant rice cultivars, Nortai, WRC43 and Sesia with the susceptible cultivar Hitomebore. Recombinant inbred line (RIL) populations derived from Nortai × Hitomebore and Sesia × Hitomebore and an F_2_ population derived from WRC43 × Hitomebore were subjected to QTL-seq (17). Using the Nortai genome as the reference (**Data S5**), the three analyses identified an overlapping interval on chromosome 11 associated with *eTO5*-triggered resistance: 24.7–29.6 Mb in Nortai, 24.4–28.4 Mb in WRC43, and 19.9–27.8 Mb in Sesia, with a common overlap of 24.7–27.8 Mb (**Fig. 3A, Fig. S9**). Subsequent fine mapping using recombinant progeny derived from Nortai × Hitomebore delimited *Pi-eTO5* to an approximately 0.5-Mb interval containing four adjacent complete NLR genes, *SQ1*, *SQ2*, *SQ3* and *SQ4* (**Fig. 3B; Fig. S10; Data S6**). High gene coverage of all four NLR genes was observed in Nortai, Sesia, and WRC43, the three cultivars that exhibited *eTO5*-triggered resistance, whereas the *SQ1*–*SQ4* haplotype was absent from the susceptible cultivar Hitomebore (**Fig. S10C**).

**Fig. 3.**
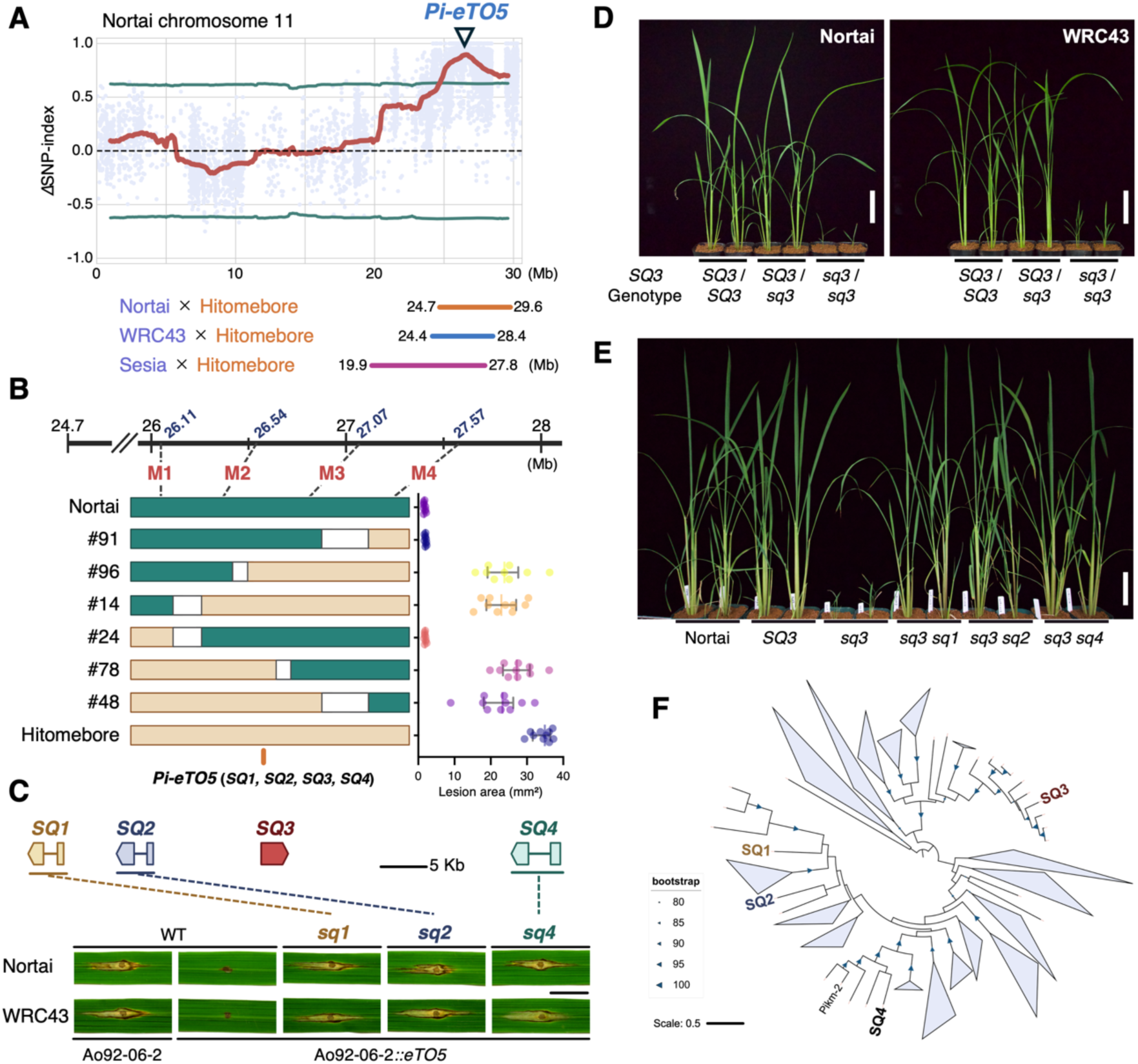
A genetically linked NLR quartet underlies *Pi-eTO5*-mediated resistance. **(A)** QTL-seq analyses of the Nortai × Hitomebore, WRC43 × Hitomebore, and Sesia × Hitomebore populations, using the Nortai genome as the reference, identified an overlapping interval between 24.7 and 27.8 Mb on chromosome 11 that was associated with *eTO5*-triggered resistance. The plot shows the ΔSNP-index for the Nortai x Hitomebore population. The red line indicates the ΔSNP-index smoothed using a sliding-window approach, and the green lines indicate the 99% confidence limits. The mapping intervals obtained from the three populations are shown below. **(B)** Fine mapping of *Pi-eTO5* using recombinant progeny derived from the Nortai × Hitomebore cross. Recombinants #91 and #24 delimited *Pi-eTO5* to an approximately 0.53-Mb interval between M2 and M3. This interval contains four adjacent NLR genes, *SQ1*, *SQ2*, *SQ3*, and *SQ4*. **(C)** Requirement of *SQ1*, *SQ2*, and *SQ4* for *eTO5*-triggered resistance. Representative punch-inoculation phenotypes of wild-type and CRISPR-Cas9 knockout lines in the Nortai and WRC43 backgrounds are shown at 10 days post-inoculation. Disruption of *SQ1*, *SQ2*, or *SQ4* abolished resistance to Ao92-06-2 expressing *eTO5*. Images shown in (C) are representative images from the corresponding experiments shown in Fig. S11. Scale bar, 1 cm. **(D)** Growth phenotypes of segregating T_2_ progeny derived from T_1_ plants heterozygous for an *SQ3* mutation in the Nortai and WRC43 backgrounds. Homozygous *sq3/sq3* plants displayed severe dwarfism, whereas wild-type (*SQ3/SQ3*) and heterozygous (*SQ3/sq3*) siblings showed normal growth. Scale bars, 10 cm. **(E)** Genetic suppression of the *sq3* dwarf phenotype. Additional disruption of *SQ1*, *SQ2*, or *SQ4* suppressed the dwarf phenotype in the *sq3* background, showing that the *sq3* dwarf phenotype requires functional *SQ1*, *SQ2*, and *SQ4*. Scale bar, 10 cm. **(F)** Maximum-likelihood phylogeny based on the NB-ARC domains of rice NLR proteins, including SQ1–SQ4. SQ1, SQ2, SQ3, and SQ4 fall into distinct phylogenetic clades, arguing against their origin through a recent local duplication event.

Target gene knockout analyses revealed that disruption of *SQ1*, *SQ2* and *SQ4* abolished resistance to Ao92-06-2 expressing *eTO5* in both the Nortai and WRC43 backgrounds (**Fig. 3C; Fig. S11**), demonstrating that all three genes are required for *eTO5*-triggered resistance. By contrast, homozygous disruption of *SQ3* caused severe dwarfism in both the Nortai and WRC43 backgrounds, whereas wild-type and *SQ3/sq3* heterozygous siblings displayed normal growth (**Fig. 3D; Fig. S12**). Because *sq3* homozygotes are dwarfed, *eTO5*-triggered resistance could not be scored in this background. Additional disruption of *SQ1, SQ2*, or *SQ4* suppressed the dwarf phenotype of *sq3* plants (**Fig. 3E; Fig. S13**), demonstrating that the dwarf phenotype caused by loss of *SQ3* depends on functional *SQ1*, *SQ2*, and *SQ4*. RNA-seq analysis of *sq3/sq3* seedlings before visible growth defects revealed broad transcriptional reprogramming, including increased expression of defense-associated genes and enrichment of stress-response pathways (**Fig. S14; Data S7–S8**). Together with the genetic suppression of *sq3* dwarfism by disruption of *SQ1*, *SQ2*, or *SQ4*, these results support a role for *SQ3* in restraining immune activity.

Phylogenetic analysis of the NB-ARC domain placed SQ1, SQ2, SQ3, and SQ4 in distinct clades, arguing against their origin through a recent local duplication event (**Fig. 3F; Data S9**). Genes corresponding to *SQ1*–*SQ4* were also identified across the japonica and indica groups of cultivated rice and in *O. rufipogon* and other wild *Oryza* species, indicating that putative complete SQ1–SQ4 haplotypes occur in a subset of cultivated and wild *Oryza* accessions (**Fig. S15; Data S10–S13**).

Together, these results identify *Pi-eTO5* as a genetically linked NLR quartet with specialized functions. *SQ1*, *SQ2*, and *SQ4* are required for *eTO5*-triggered resistance, whereas *SQ3* is required to prevent constitutive defense activation that depends on *SQ1, SQ2,* and *SQ4,* thereby avoiding associated growth defects.

### Evolutionary conservation and molecular associations of eTO5

eTO5 is broadly conserved across the B71 pandemic clonal lineage, with increased copy number in several geographically dispersed derivatives (**Fig. 4A**; **Data S14–S15**). eTO5 and eTO1, both members of the ZiF effector family (18) (**Fig. S16A**), show a similar copy-number increase in isolates from the geographically expanded derivatives of this lineage (**Fig. 4A; Fig. S16B**). This parallel copy-number increase suggests that copy-number expansion can occur in multiple structurally related effectors rather than being unique to eTO5. In contrast, eTO16 and eTO31 are consistently absent from Bangladeshi isolates within the pandemic lineage (**Fig. 4A**), indicating that both copy-number expansion and gene loss contribute to effector repertoire variation within the pandemic lineage.

**Figure 4.**
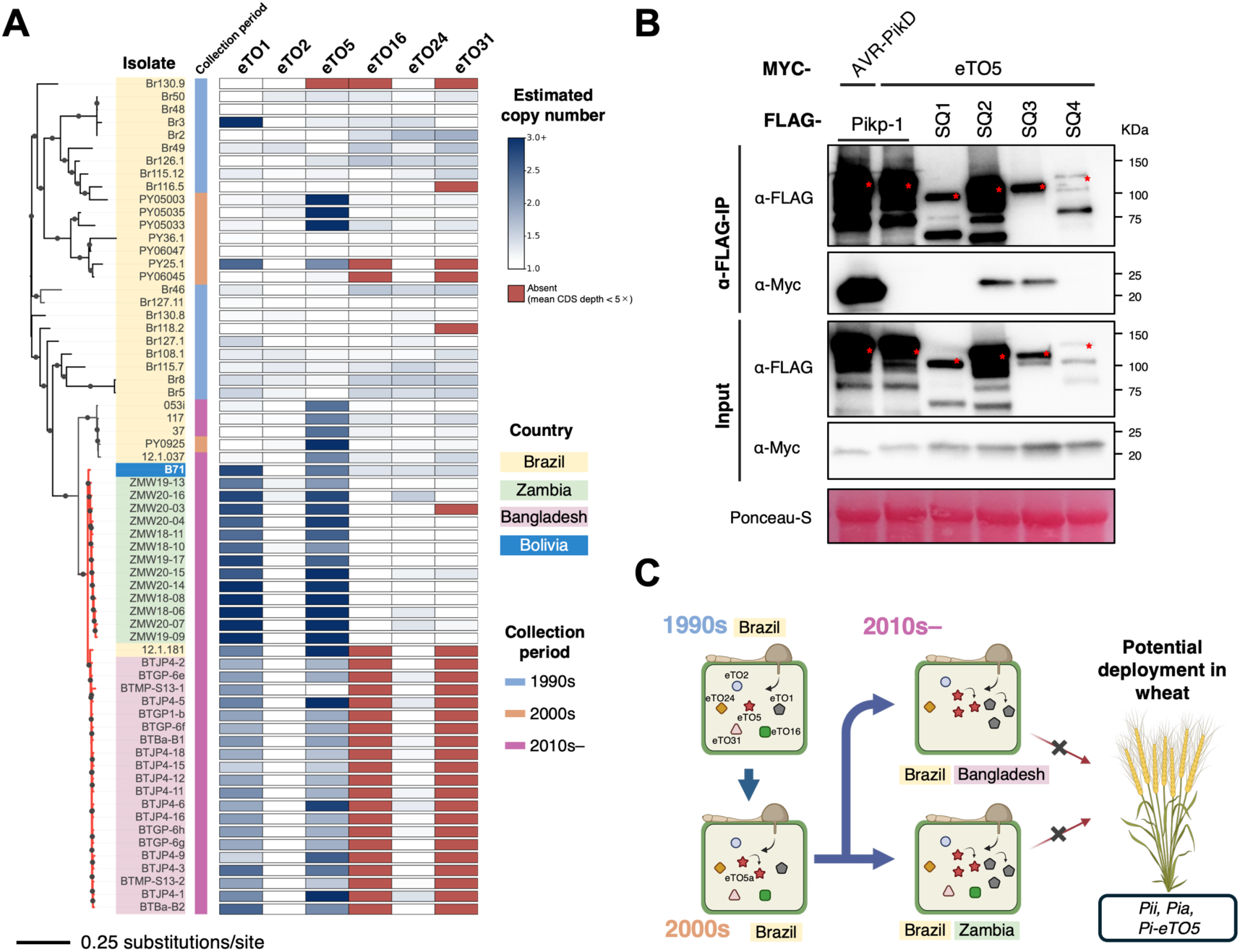
Phylogenetic distribution of *eTO* copy-number variation and functional association of eTO5 with Pi-eTO5 NLRs. **(A)** Maximum-likelihood phylogenetic tree of 65 *M. oryzae* pathotype *Triticum* (MT) isolates based on genome-wide SNPs, midpoint-rooted and annotated with copy-number (CN) estimates for six eTO loci (*eTO1*, *eTO2*, *eTO5*, *eTO16*, *eTO24*, and *eTO31*). Geographic origin (Brazil, Zambia, Bangladesh, and Bolivia) and collection period are indicated alongside the tree. Heatmap colors represent estimated copy number: red indicates loci classified as absent (mean CDS depth < 5×), white indicates loci within the isolate-specific single-copy range, and the blue gradient indicates increasing CN estimates above the isolate-specific empirical BUSCO P95 threshold. CN estimates and BUSCO normalization statistics are provided in **Data S15** and S**16**, respectively. The scale bar indicates 0.25 substitutions per site. **(B)** Co-immunoprecipitation assays examining association of eTO5 with Pi-eTO5-linked NLR proteins. 4×Myc-tagged eTO5 was co-expressed with 3×FLAG-tagged SQ1, SQ2, SQ3, or SQ4 in Nicotiana benthamiana. FLAG-tagged proteins were immunoprecipitated using anti-FLAG agarose, and input and immunoprecipitated fractions were analyzed by immunoblotting with anti-FLAG and anti-Myc antibodies. Pikp-1–3×FLAG and AVR-PikD–4×Myc were included as controls. eTO5 co-immunoprecipitated with SQ2 and SQ3 but was not detected with SQ1 or SQ4 under these conditions. Asterisks indicate nonspecific bands. **(C)** Schematic model illustrating variation in the *eTO* repertoire during geographic expansion of the MT lineage and the potential use of rice resistance genes recognizing MT effectors for deployment in wheat. *eTO5a* denotes a diverged *eTO5* copy present in later isolates (**Fig. S16C**).

To characterize the functional features of the *Pi-eTO5*-linked NLRs, we analyzed the protein sequences of SQ1, SQ2, SQ3, and SQ4. SQ4 uniquely contains a MADA motif (19) (**Fig. S17**), a conserved N-terminal motif associated with NLR-mediated cell-death activity. We next examined whether eTO5 associates with the Pi-eTO5 NLR proteins using co-immunoprecipitation (co-IP) assays following transient expression of the proteins in *Nicotiana benthamiana*. eTO5 co-immunoprecipitated with SQ2 and SQ3, but not with SQ1 or SQ4 (**Fig. 4B; Fig. S18**). This selective association pattern, together with the distinct distribution of NLR-associated motifs among SQ1–SQ4, is consistent with functional specialization among the four *Pi-eTO5*-linked NLRs.

## Discussion

*Magnaporthe oryzae* exhibits strong host specificity to different grass species. Here we identified six wheat blast effector genes that trigger resistance in rice. Notably, *eTO2* is the only effector recognized by all rice cultivars tested, suggesting that recognition of *eTO2* may contribute broadly to the host-specific barrier against MT in rice. The absence of a susceptible genotype among the cultivars tested prevented genetic mapping of the corresponding resistance determinant. This broad recognition could reflect one or more widely distributed rice immune receptors, although its genetic basis remains unresolved. Together, these results reveal overlapping layers of effector recognition within rice nonhost resistance to MT. Although recognition of individual effectors such as eTO5 segregates among rice genotypes, the pathogen appears to remain unable to infect because additional recognition specificities mask this segregation. Thus, effector-by-effector screening exposes genetic variation in resistance that is hidden when nonhost resistance is examined only at the whole-pathogen level. Uniformity at the species level is therefore emergent from overlapping specificities rather than a property of any single receptor.

Unlike *eTO2*, wheat blast effectors including *eTO1, eTO5, eTO16, eTO24* and *eTO31* were recognized by a subset of rice cultivars. Of these, *eTO1-* and *eTO24-*triggered resistance were genetically dependent on *Pii* and *Pia*, respectively. Focusing on *eTO5*, we isolated the cognate resistance locus using our genetic pipeline. These results support the idea that recognition of multiple MT effectors contributes substantially to the nonhost resistance of rice. The contrast with wheat is informative. Under selection from wheat, MT lost *PWT3* and escaped *Rwt3* (12). Rice imposes no reciprocal selection, and MT accordingly retains effectors such as *eTO1* and *eTO24* that rice R genes still detect.

*Pi-eTO5* comprises four tightly linked NLR genes with genetically distinct contributions to immunity and growth regulation. Disruption of *SQ1*, *SQ2*, or *SQ4* abolished *eTO5*-triggered resistance, whereas loss of *SQ3* caused severe dwarfism and constitutive defense-associated transcriptional activation. The dwarf phenotype of *sq3/sq3* plants was suppressed by additional disruption of *SQ1*, *SQ2*, or *SQ4*, indicating that *SQ3* is required to restrain immune activity dependent on the other three NLRs. Consistent with functional specialization within the quartet, SQ4 uniquely contains a MADA motif, whereas eTO5 co-immunoprecipitated with SQ2 and SQ3. These observations support a model in which the four linked NLRs perform distinct but coordinated functions, although the molecular sequence of effector recognition and immune activation remains to be established.

*eTO5* and *eTO1* both show copy-number increases within the B71 clonal lineage (**Fig. 4A; Data S14–S16**), with elevated copy-number estimates observed in geographically dispersed derivatives of the lineage. In contrast, *eTO16* and *eTO31* are absent from the Bangladeshi isolates examined, illustrating that both copy-number expansion and gene loss contribute to variation in the effector repertoire of the pandemic lineage. We summarize these patterns in a model in which the effector repertoire varies during geographic spread of the lineage, while several *eTOs* recognized by rice resistance genes remain present in currently circulating isolates (**Fig. 4C**). The retention of *eTO5* across the sampled B71 pandemic isolates provides a rationale for testing *Pi-eTO5* as a source of wheat blast resistance. Whether all four NLRs are required in a heterologous background, and whether a four-gene cassette can be deployed in wheat, remain to be determined.

More broadly, this functional screening approach provides a strategy for genetically accessing nonhost resistance that would otherwise be difficult to map. By transferring individual effectors into a related pathogen that can infect the nonhost species, complex resistance to the intact pathogen can be decomposed into genetically tractable recognition phenotypes. Applied across diverse host–pathogen systems, this strategy could facilitate systematic discovery of AVR– R relationships and uncover resistance genes with potential for deployment in crops. Thus, effector-driven screening converts genetically inaccessible nonhost resistance into tractable phenotypes for resistance-gene discovery.

## Supporting information

DataS1

DataS2

DataS3

DataS4

DataS5

DataS6

DataS7

DataS8

DataS9

DataS10

DataS11

DataS12

DataS13

DataS14

DataS15

DataS16

DataS17

## Acknowledgements

This study was supported by JSPS KAKENHI grants 23K20042 and 24H00010 (to R.T.) and 25K01992 (to M.S.). Some computations were performed on the supercomputer at the National Institute of Genetics, Research Organization of Information and Systems. We thank K. Oikawa, S. Nagata, S. Takei, and T. Fujiwara for technical assistance. We thank the National BioResource Project at the National Institute of Genetics, Japan, for providing seeds of the wild Oryza accessions. We also thank the Genebank of the National Agriculture and Food Research Organization, Japan, for providing seeds of accessions from the Japanese Rice Core Collection and the World Rice Core Collection (WRC1, WRC18, and WRC43).Created in BioRender. SHIMIZU, M. (2026) https://BioRender.com/3j4650t

## Materials and Methods

### Fungal isolates and culture conditions

The *M. oryzae* isolates Ao92-06-2 and Br48 were maintained on potato sucrose agar (PSA). For experiments, isolates were transferred to oatmeal agar and cultured at 25°C for 7 days. Aerial mycelia were then removed by scraping to induce conidiation, and the cultures were incubated under near-ultraviolet light for 2 days.

### Rice inoculation and lesion quantification

Rice leaf-blade punch inoculation assays were performed using the indicated *M. oryzae* strains. Conidial suspension adjusted to 3×10⁵ conidia mL⁻¹ was applied to punched sites on leaves of rice plants approximately 1 month after sowing. Prior to inoculation, wild-type plants were grown in a naturally lit greenhouse under ambient temperature conditions, whereas transgenic plants were grown in a naturally lit P1 containment greenhouse maintained at approximately 28°C. Inoculated plants were maintained in a dew chamber at 27°C in darkness for 24 h and then transferred to a growth chamber under a 16-h light/8-h dark photoperiod at 27°C. Lesion areas were quantified using ImageJ software (1).

### Genome assembly of Br48

Publicly available Illumina paired-end reads of the *M. oryzae* pathotype *Triticum* isolate Br48 (DDBJ Sequence Read Archive accession DRX054196; 2) were downloaded and quality-filtered using FaQCs v2.10 (3). Filtered reads were assembled *de novo* using MaSuRCA (4) with default parameters. Assembly completeness was assessed using BUSCO v5.5.0 in genome mode with the sordariomycetes_odb10 Data set (5). The assembly contained 97.1% complete BUSCOs (96.8% single-copy and 0.3% duplicated), 0.9% fragmented BUSCOs, and 2.0% missing BUSCOs among 3,817 ortholog groups. Of these, 3,696 BUSCO genes classified as complete and single-copy were subsequently used for depth normalization in the effector copy-number analysis (**Fig. 4A**). Assembly metrics are summarized in **Data S1**.

### Identification of MT effector candidates

Seeds of the barley cultivar Nigrate were sown in soil, and 1-week-old seedlings were inoculated with *M. oryzae* pathotype *Triticum* isolate Br48. Leaf tissues were collected at 24 hpi from multiple plants and bulked, and total RNA was extracted using the SV Total RNA Isolation System (Promega). RNA-seq libraries were prepared using the NEBNext Ultra II Directional RNA Library Prep Kit for Illumina (New England Biolabs) and sequenced to generate paired-end reads (DDBJ Sequence Read Archive accession DRR594671). RNA-seq reads were quality-filtered using FaQCs v2.10 (3) and aligned to the assembled Br48 genome using HISAT2 v2.2.1 (6) with the -- dta option. Genome-guided transcript assembly and transcript abundance estimation were performed using StringTie v2.2.1 (7), and transcript sequences were extracted using gffread v0.12.7 (8). Transcripts shorter than 200 bp were excluded, resulting in 7,774 transcript sequences. Open reading frames were predicted using TransDecoder v5.5.0 (9), and exact duplicate protein sequences were removed using CD-HIT v4.8.1 (10) with a sequence identity threshold of 1.0. Candidate cytoplasmic effector genes were selected using the pipeline shown in **Fig. S2** using the following criteria: (i) the encoded protein contained a signal peptide predicted using SignalP v4.1 (11); (ii) transcript abundance exceeded 100 TPM at 24 hpi; (iii) the protein was classified as a cytoplasmic or dual apoplastic/cytoplasmic effector by EffectorP v3.0 (12); and (iv) the predicted protein was shorter than 150 amino acids. In total, 114 candidate cytoplasmic effector genes were identified.

### Selection of *eTO* genes

To identify MT-specific effector candidates, publicly available whole-genome sequencing reads from Br48 and nine MO isolates (**Data S2**) were quality-filtered using FaQCs v2.10 (3) and aligned to the Br48 genome using Bowtie2 v2.3.5.1 (13). Alignments were sorted and indexed using SAMtools v1.19.2 (14). For each of the 114 candidate effector genes, sequence coverage was calculated using mosdepth v0.3.6 (15) as the proportion of nucleotide positions supported by a read depth of at least 3× (**Fig. S3**). Genes were operationally defined as absent from MO when sequence coverage was ≤30% in all nine MO isolates. This analysis identified 35 MT-specific candidate effector genes, designated *eTO1* to *eTO35* (effector unique to MT versus MO; **Data S 3**).

### Generation of *eTO*-expressing MO transformants

The coding sequences of *eTO1* to *eTO35*, except *eTO29*, were amplified by PCR from cDNA prepared from barley leaves infected with the MT isolate Br48 using the primers listed in **Data S17**. Because *eTO29* could not be amplified from cDNA, a synthetic coding sequence corresponding to the predicted Br48 *eTO29* sequence was obtained from Azenta Life Sciences. The amplified or synthesized coding sequences were cloned downstream of the *pex22* (*AVR-Pia*) promoter in the pCB1531-*pex22p* vector described previously (16), either by digestion with BamHI and XbaI followed by ligation or by In-Fusion cloning (Takara Bio) into the BamHI/XbaI -linearized vector. The method used for each construct is indicated in **Data S17**. The resulting constructs were individually introduced into the MO isolate Ao92-06-2 by polyethylene glycol– mediated transformation of fungal protoplasts as described previously (17). Transformants were selected on bialaphos-containing medium, and the presence of each construct was confirmed by PCR. Two independent transformants for each *eTO* gene were used in the primary screen.

### Effector screening across rice cultivars

The resulting Ao92-06-2 transformants expressing individual *eTO* genes were inoculated onto 20 rice cultivars susceptible to the parental isolate Ao92-06-2, yielding 700 cultivar–*eTO* combinations (**Fig. S4**; **Data S4**). Cultivar–*eTO* combinations scored as resistant in the primary screen were retested in a secondary screen to confirm the phenotype. An interaction was scored as resistant (R) when no lesion expansion beyond the inoculation site was observed and as susceptible (S) when any lesion expansion was detected.

### Genetic validation of *eTO1*- and *eTO24*-triggered resistance

To test whether *eTO1*-triggered resistance requires *Pii*, Hitomebore and two independent *pii* mutant lines, Hit5948 (18) and Hit5882 (19), were inoculated with Ao92-06-2 expressing *eTO1*. Similarly, to test whether *eTO24*-triggered resistance requires *Pia*, Sasanishiki and two independent *rga4* mutant lines, Sas2127 and Sas1943 (20), were inoculated with Ao92-06-2 expressing *eTO24*. Inoculations were performed using the punch inoculation procedure described above.

### Rice genome sequencing and *de novo* assembly

High-molecular-weight genomic DNA was extracted from leaves of the rice cultivars Nortai, WRC43, and Sesia using the NucleoBond HMW DNA kit (Macherey-Nagel) according to the manufacturer’s instructions. Libraries were prepared using the Ligation Sequencing Kit V14 (SQK-LSK114; Oxford Nanopore Technologies) and sequenced on a PromethION 2 Solo using R10.4.1 flow cells. Raw signal data were base-called using Dorado v0.9.6 with the dna_r10.4.1_e8.2_400bps_sup@v5.0.0 model and a minimum Q-score of 12. Reads from each cultivar were assembled *de novo* using hifiasm v0.25.0-r726 in ONT mode (21). Assemblies were generated with duplication purging disabled (-l 0), an estimated genome size of 400 Mb, and the - -rl-cut and --sc-cut parameters set to 10,000 and 12, respectively. Primary contig graphs were converted to FASTA format using gfatools. Contigs were ordered and oriented into pseudochromosomes using RagTag v2.1.0 (22), with IRGSP-1.0 (Nipponbare) as the reference genome. Assembly completeness was assessed using BUSCO v5.5.0 (5) with the poales_odb10 lineage data set. Assembly statistics are provided in **Data S5**.

### QTL-seq analysis

Three mapping populations were analyzed: 50 Nortai × Hitomebore recombinant inbred lines (RILs), 83 WRC43 × Hitomebore F₂ individuals, and 38 Sesia × Hitomebore RILs. Resistance to Ao92-06-2::*pex22p*:*eTO5* was evaluated by punch inoculation, and lesion areas were measured using ImageJ at 7–9 days post-inoculation. Individuals or lines with the smallest and largest lesion areas were selected for the resistant bulk (RB) and susceptible bulk (SB), respectively. Each bulk contained 20 individuals or lines for the Nortai × Hitomebore and WRC43 × Hitomebore populations and 15 RILs for the Sesia × Hitomebore population. Genomic DNA was extracted using the NucleoSpin Plant kit (Macherey-Nagel), pooled according to phenotype, and used to prepare libraries with the Collibri ES DNA Library Prep Kit for Illumina (Thermo Fisher Scientific). Paired-end sequencing was performed on an Illumina HiSeq X platform. Reads were quality-filtered using fastp (23) and aligned to the Nortai reference genome using Bowtie2 (13). BAM files were analyzed using the QTL-seq pipeline (18, 24). SNP-index and ΔSNP-index values were calculated for each population, and genomic regions exceeding the 99% confidence limits were considered candidate QTL regions.

### Fine mapping of *Pi-eTO5*

CAPS markers were designed from sequence polymorphisms between Nortai and Hitomebore using V-primer (25). PCR products were digested with XbaI or BamHI to distinguish the parental alleles. Marker sequences, primer pairs, and restriction enzymes are provided in **Data S17**. From a population of 250 Nortai × Hitomebore RILs, lines carrying recombination events between markers M1 and M4 were selected and genotyped with additional markers within this interval. Recombinant lines were inoculated with Ao92-06-2 and Ao92-06-2::*pex22p*:*eTO5*. Recombinant haplotypes and *eTO5*-triggered resistance phenotypes were used to delimit the *Pi-eTO5* locus.

### Transcriptome-guided gene annotation and identification of candidate NLR genes

Total RNA was extracted from leaves of the rice cultivar Nortai using the SV Total RNA Isolation System (Promega). Strand-specific RNA-seq libraries were prepared using the NEBNext Ultra II Directional RNA Library Prep Kit for Illumina (New England Biolabs) and sequenced on an Illumina HiSeq X platform to generate paired-end reads. Reads were adapter-trimmed and quality-filtered using fastp (23), aligned to the Nortai genome assembly using HISAT2 (6), and subjected to genome-guided transcript assembly using StringTie (7). Transcript sequences were extracted using gffread (8), and open reading frames were predicted using TransDecoder (9). Predicted proteins were analyzed using NLRtracker (26). Complete CNLs containing coiled-coil (CC), nucleotide-binding NB-ARC, and leucine-rich repeat (LRR) domains were retained as candidate NLRs. whereas degenerate NLRs lacking an intact NB-ARC domain were excluded. Four complete CNLs were identified within the *Pi-eTO5* candidate interval: *Nortai.21965.1* (*SQ1*), *Nortai.21966.1* (*SQ2*), *Nortai.21967.1* (*SQ3*), and *Nortai.21968.1* (*SQ4*). Nortai.21962.5, a degenerate NLR lacking an intact NB-ARC domain, was excluded. Genes supported by leaf RNA-seq and annotated within this interval are listed in **Data S6**. The presence or absence of the *SQ1*– *SQ4* haplotype was concordant with the eTO5-response phenotype: lines carrying *SQ1*–*SQ4* were resistant, whereas lines lacking the haplotype were susceptible (**Fig. S10B**).

### Gene coverage analysis of *SQ1*–*SQ4* across rice cultivars

To examine the distribution of *SQ1*, *SQ2*, *SQ3*, and *SQ4* across the 20 rice cultivars used in the functional screen, whole-genome sequencing reads were analyzed. Publicly available reads for the Japanese Rice Core Collection (JRC) and World Rice Core Collection (WRC) accessions were obtained from Tanaka et al. (27, 28), whereas reads for Hitomebore, Moukoto, Nortai, Sasanishiki, Sesia, and WRC43 were generated in this study. Reads were adapter-trimmed and quality-filtered using fastp (23), aligned to the Nortai reference genome using Bowtie2 v2.3.5.1 (13), and processed using SAMtools v1.19.2 (14). For each cultivar, gene coverage of *SQ1, SQ2, SQ3,* and *SQ4* was calculated using mosdepth v0.3.6 (15) as the percentage of nucleotide positions within each annotated gene region supported by a read depth of at least 3×. Coverage values were visualized as a heatmap. Cultivar names, sequence accession numbers, and data sources are provided in **Data S4**. Responses to Ao92-06-2::*pex22p*:*eTO5* were determined by the functional screening described above and are shown below the heatmap in **Fig. S10C**.

### Generation of rice knockout mutants using the CRISPR-Cas9 system

Rice knockout mutants of *SQ1, SQ2, SQ3*, and *SQ4* were generated using the CRISPR-Cas9 system described previously (29). Guide RNA target sequences were designed using CRISPRdirect (30). Complementary oligonucleotides corresponding to each target sequence were annealed and cloned into the pU6::ccdB::gRNA vector at the BbsI site. The resulting gRNA cassette containing the OsU6 promoter was subsequently transferred into the pZH::gYSA::MMCas9 binary vector using AscI and PacI. Constructs were introduced into *Agrobacterium tumefaciens* strain EHA101 and transformed into the rice cultivars Nortai and WRC43 as described previously (20). Regenerated T_0_ plants were genotyped by PCR and Sanger sequencing of the targeted regions. Independent mutant alleles carrying mutations predicted to disrupt protein function were selected for subsequent analyses. T_1_ plants or segregating T_2_ progeny were used for phenotypic analyses, as indicated in the corresponding figures. For *SQ3*, heterozygous T_1_ plants were self-pollinated, and the resulting T_2_ progeny were genotyped by PCR– restriction fragment length polymorphism analysis using AcuI or EcoNI to distinguish wild-type, heterozygous, and homozygous mutant siblings. Mutations in the other edited genes were determined by Sanger sequencing. Guide RNA target sequences and primers used for genotyping are provided in **Data S17**.

### Generation and genotyping of double mutants

Double-knockout plants were generated by introducing CRISPR-Cas9 constructs targeting *SQ1, SQ2*, or *SQ4* into calli induced from segregating T_2_ seeds obtained from an *SQ3/sq3* T_1_ plant. After regeneration, plants were genotyped at the *SQ3* locus and the additionally targeted *SQ* gene. Among regenerated plants homozygous for the *sq3* mutation, none carried a wild-type or monoallelic mutant genotype (WT/mt1) at the additionally targeted locus; all carried either homozygous (mt1/mt1) or biallelic (mt1/mt2) mutations. These plants were used as double-knockout lines for growth analyses. Guide RNA target sequences and primers used for genotyping are provided in **Data S17**.

### RNA sequencing and differential gene expression analysis of segregating *SQ3/sq3* progeny

Segregating T_2_ progeny derived from a self-pollinated T_1_ plant heterozygous for the *SQ3* mutation were grown on Murashige and Skoog medium under sterile conditions. At 7 days after germination, seedlings were genotyped, and five individual *SQ3/SQ3* and five individual *sq3/sq3* seedlings were selected before visible growth defects appeared. Each seedling was analyzed independently as a biological replicate. Total RNA was extracted from whole seedlings using the SV Total RNA Isolation System (Promega). Strand-specific RNA-seq libraries were prepared using the NEBNext Ultra II Directional RNA Library Prep Kit for Illumina (New England Biolabs) and sequenced on an Illumina NovaSeq platform to generate paired-end reads. Reads were adapter-trimmed and quality-filtered using fastp (23). Because the *SQ1*–*SQ4* locus is absent from the public IRGSP-1.0 Nipponbare reference genome, the corresponding Nortai genomic region was appended to the reference together with gene models for *SQ1*–*SQ4*. Reads were aligned to this composite reference genome using HISAT2 (6), retaining only properly paired, uniquely mapped read pairs (MAPQ ≥ 20). Gene-level read counts were obtained using featureCounts (31). Differential expression was analyzed a quasi-likelihood F-test in edgeR (32), with genotype (*sq3/sq3* versus *SQ3/SQ3*) as the sole factor. Genes with a false-discovery rate (FDR) < 0.05 and |log2-fold change| ≥ 1 were considered differentially expressed (**Data S7**). For principal component analysis, raw gene counts were normalized by variance-stabilized using DESeq2 (33), and PCA was performed on the transformed values. For visualization of individual gene expression, transcript abundance was expressed as transcripts per million (TPM), calculated from gene-level read counts using the exon-union gene length reported by featureCounts, and shown as log2 (TPM + 1).

Functional enrichment analysis of genes significantly upregulated in each genotype was performed using clusterProfiler v4.10.0 (34) with a custom Gene Ontology Biological Process (GO BP) annotation derived from the RAP-DB IRGSP-1.0 representative gene annotation. The background set was restricted to retained after low-count filtering in the edgeR analysis. GO BP terms with Benjamini-Hochberg (BH)-adjusted P value < 0.05 were considered significantly enriched (**Data S8**). Genes upregulated in *sq3/sq3* within significantly enriched defense-related GO terms were visualized as heat maps of VST-normalized, row-scaled expression values. Immune-related genes belonging to NB-LRR, WRKY, pathogenesis-related, and receptor-like kinase families were identified from RAP-DB annotation using InterPro domain and description keywords and visualized in the same manner.

### Phylogenetic analysis of rice NLR proteins

Predicted protein sequences ≥800 amino acids in length from the *Oryza sativa* ssp. *japonica* cv. Nipponbare reference annotation were obtained from the Rice Annotation Project Database (RAP-DB) (35). NLR proteins were identified using NLRtracker (26). Degenerate NLRs lacking an intact NB-ARC domain were excluded, and genes sharing a duplicate gene identifier were consolidated into single entries. The predicted protein sequences of SQ1, SQ2, SQ3, and SQ4, together with the paired NLRs Pii-1 (AB820896) and Pii-2 (LC190730) (18, 19), Pikm-1 (AB462324) and Pikm-2 (AB462325) (36), Pit-1 (AB379816) and Pit-2 (AB379819) (37), and RGA4 (AB604622) and RGA5 (AB604627) (20), were added to the Nipponbare NLR data set. NB-ARC domain sequences were extracted from each NLR protein and aligned using MUSCLE v5.2 (38). A maximum-likelihood phylogenetic tree was inferred using RAxML-NG v1.2.2 under the JTT+G4+F amino acid substitution model with 1,000 nonparametric bootstrap replicates (39, 40). The resulting tree was visualized and annotated using iTOL v6 (41). Genes and NB-ARC domain sequences used in this analysis are listed in **Data S9**.

### Conservation analysis of *SQ1*–*SQ4* across *Oryza* genomes

To assess the conservation of *SQ1, SQ2, SQ3*, and *SQ4* across cultivated and wild rice, we first analyzed Nanopore-based genome assemblies of *O. sativa japonica* and *indica* and the wild progenitor *O. rufipogon* (42–44). The predicted Nortai amino acid sequences of SQ1, SQ2, SQ3, and SQ4 were used as queries against each genome assembly using Exonerate v2.4.0 in protein2genome mode (45), and the highest-scoring alignment was retained for each gene and accession. Relative similarity to Nortai was calculated by normalizing the Exonerate bit score for each accession to that obtained by aligning the corresponding Nortai sequence to the Nortai genome (relative score = Bᵢ,ⱼ / Bᵢ,Nortai) (**Data S10**). Accessions with relative scores ≥ 0.6 for all four genes were conservatively classified as carrying a putative complete SQ1–SQ4 haplotype. Such haplotypes were detected in 20 of 143 *O. sativa* accessions and 15 of 109 *O. rufipogon* accessions, whereas most other accessions showed only low alignment scores, consistent with presence/absence-type variation of the SQ1–SQ4 region. We next extended the analysis to a broader panel of wild *Oryza* species using short-read-based genome assemblies (46). Putative complete SQ1–SQ4 haplotypes were likewise detected in only 12 of 108 accessions, which were largely restricted to *O. glumaepatula* (**Data S10**). Because these assemblies were generated from short reads, however, physical linkage of the four genes could not be confirmed. To examine the locus structure directly, we generated Nanopore *de novo* genome assemblies for two representative wild accessions, W2149 (*O. glumaepatula*) and W1577 (*O. punctata*). Genomic DNA was sequenced using Oxford Nanopore R9.4.1 flow cell chemistry. Reads were base-called using Dorado v0.9.0 with the dna_r9.4.1_e8_sup@v3.6 super-accuracy model and a minimum q-score of 8, error-corrected with Ratatosk (47) using newly generated Illumina short reads, and assembled *de novo* using hifiasm v0.25.0-r726 in ONT mode (**Data S11**). For accessions classified as carrying putative complete SQ1–SQ4 haplotypes, amino acid sequences of SQ1–SQ4 were manually determined from CDS-based alignments and used for haplotype comparison (**Data S12 and S13**). To examine the evolutionary relationships of SQ1, SQ2, SQ3, and SQ4 alleles, amino acid sequences were aligned independently for each gene using MAFFT v7 with the –auto option (48). Maximum-likelihood phylogenetic trees were inferred using IQ-TREE v2 with automatic model selection (-m MFP) (49, 50), with branch support assessed using 1,000 ultrafast bootstrap replicates and 1,000 SH-aLRT replicates (-bb 1000 -alrt 1000).

### Conservation analysis of AVR–active *eTO* genes across MT isolates

Sixty-five *M. oryzae* pathotype *Triticum* (MT) isolates were characterized (**Data S14**). Whole-genome short-read sequencing data were obtained from public databases (2, 51–56). Quality-trimmed paired-end data were mapped to the Br48 reference genome using BWA MEM (57). PCR duplicates were marked using Picard MarkDuplicates, and BAM files were indexed with SAMtools (14).

A set of 3,696 complete single-copy BUSCO genes identified using the sordariomycetes_odb10 data set (5) was used for depth normalization. Mean read depth across BUSCO loci and the coding sequences of six AVR-active *eTO* genes (*eTO1, eTO2, eTO5, eTO16, eTO24*, and *eTO31*) was calculated using mosdepth (15), retaining reads with mapping quality ≥20. For each isolate, normalized copy-number (CN) was calculated as the mean depth of the *eTO* coding sequence divided by the median depth of the 3,696 BUSCO genes. An eTO locus with a mean CDS depth <5× was classified as absent. For loci with mean CDS depth ≥5×, an isolate-specific single-copy threshold was defined as the empirical 95th percentile of normalized BUSCO depths (BUSCO P95 depth/BUSCO median depth). Loci with normalized CN values above this threshold were classified as amplified, whereas those at or below the threshold were classified as single-copy. Because depth was measured against the Br48 reference assembly, CN values reflect the combined depth of each *eTO* locus and of any closely related sequences that are present in an isolate but not represented in the reference. CN estimates are provided in **Data S15**, and BUSCO depth statistics and isolate-specific thresholds are provided in **Data S16**.

SNPs were called against the Br48 reference genome using bcftools (14). Filtering: Variants were retained if they were biallelic SNPs with QUAL ≥30, depth ≥10, and ≤10% missing data. The filtered VCF was converted to PHYLIP format using vcf2phylip (58). Maximum-likelihood phylogenetic trees were inferred using IQ-TREE 2 (59) under the GTR+ASC model (60), with branch support assessed using 1,000 ultrafast bootstrap replicates (61) and SH-aLRT (62). Trees were visualized and annotated using iTOL (41) with isolate metadata and copy-number data from **Data S15**.

### *De novo* assembly of MT isolates and assembly-based detection of eTO5-, eTO5a-, and eTO1-related sequences

To obtain assembly-based evidence for the presence of multiple eTO5- and eTO1-related sequences in representative MT isolates (**Fig. S16B**), publicly available Oxford Nanopore long reads were assembled *de novo* for selected isolates from Bangladesh and Zambia. Nanopore reads for the Bangladeshi isolates were obtained from BioProject PRJEB27137 (MinION; 63), and those for the Zambian isolates from BioProject PRJEB53707 (PromethION; The John Innes Centre, 2022). Matched Illumina paired-end reads used for error correction were obtained from BioProject PRJEB22035 for the Bangladeshi isolates (55) and PRJEB43836 for the Zambian isolates (56).

Illumina reads were adapter-trimmed and quality-filtered using fastp v1.3.6 (23). Nanopore reads were hybrid error-corrected with the matched Illumina reads using Ratatosk v0.9.0 (47) in correct mode. Corrected reads were assembled *de novo* using hifiasm v0.25.0-r726 (21) in ONT mode with duplication purging disabled, and the primary contig graph was converted to FASTA format using gfatools v0.4-r214.

The amino acid sequences of eTO5, eTO5a, and eTO1 were searched against each assembly using tblastn v2.12.0+ (64) with default parameters. Because eTO5 and eTO5a differ at only 7 of 96 amino acid positions and therefore cross-detect each other, only alignments spanning the full query length with 100% amino acid identity were retained, allowing each retained hit to be assigned unambiguously to one of the three query sequences. The positions of retained hits were plotted along the corresponding contigs using a custom Python script based on matplotlib v3.6.3.

### Nicotiana benthamiana growth conditions

Wild-type *Nicotiana benthamiana* plants were grown in a growth chamber at 23-25°C under long-day conditions (16-h light/8-h dark).

### Co-immunoprecipitation assay

To test associations between the Pi-eTO5-linked NLRs and eTO5, NLR (SQ1, SQ2, SQ3, SQ4, and Pikp-1) and effector (eTO5 and AVR-PikD) genes were synthesized with codon optimization for *Nicotiana benthamiana* (Integrated DNA Technologies). The synthesized genes were assembled into binary vector pJK268c (26) using the Golden Gate system with the MoClo Tool Kit (65) and MoClo Plant Parts Kit (66). The promoter module pICH51288 (2×35S; Addgene no. 50269) and terminator module pKH1001, containing synthesized *Arabidopsis thaliana HEAT SHOCK PROTEIN 18.2* (At5g59720) terminator, were used for all constructs. NLRs and effectors were tagged using 3×FLAG (pICSL50007; Addgene no. 50308) and 4×Myc (pICSL30009; Addgene no. 50301), respectively.

Transient gene expression in *N. benthamiana* was performed by agroinfiltration as described previously (67). Briefly, leaves of four-week-old *N. benthamiana* plants were infiltrated with *Agrobacterium tumefaciens* strain GV3101 carrying the indicated binary constructs using needleless syringes. *Agrobacterium* cultures were resuspended in infiltration buffer containing 10 mM MES-NaOH (pH 5.6), 10 mM MgCl_2_, and 150 µM acetosyringone and adjusted to an OD_600_ of 0.5. Cultures carrying the NLR and effector constructs were mixed at a 1:1 ratio before infiltration.

Two infiltrated leaves were harvested 3 days after infiltration, frozen in liquid nitrogen, and ground to a fine powder. Total proteins were extracted with two volumes of ice-cold GTEN-based extraction buffer relative to tissue fresh weight [25 mM Tris-HCl (pH 7.5), 150 mM NaCl, 1 mM EDTA, 10% (v/v) glycerol, 10 mM DTT, 0.3% (v/v) NP-40, 2% (w/v) polyvinylpolypyrrolidone (PVPP), and protease inhibitor cocktail (Merck, P9599)]. Extracts were clarified by centrifugation at 10,000 × *g* for 10 min at 4°C, and an aliquot of each clarified protein extract was used as the input sample. FLAG-tagged NLR proteins were immunoprecipitated by incubating the clarified extracts with 20 µL of anti-FLAG agarose beads (Merck, A2220) for 1 h at 4°C. Beads were washed five times with GTEN buffer supplemented with 0.3% (v/v) NP-40. Bound proteins were eluted by incubation with FLAG peptide at a final concentration of 150 µg mL⁻¹ for 30 min at 4°C. Immunoprecipitated proteins and input samples were separated by SDS-PAGE and transferred to PVDF membranes using a Trans-Blot Turbo Transfer System (Bio-Rad). Membranes were blocked with 3% (w/v) skim milk in TBS-T [50 mM Tris-HCl (pH 7.6), 150 mM NaCl, and 0.05% (v/v) Tween 20] for 1 h at room temperature and subsequently incubated with Horseradish peroxidase (HRP)-conjugated anti-FLAG (Merck, A8592) or anti-Myc (Santa Cruz Biotechnology, SC-40 HRP) antibodies at a 1:5,000 dilution for 1 h at room temperature. After washing with TBS-T, chemiluminescent signals were detected using SuperSignal West Dura Extended Duration Substrate (Thermo Scientific) and a Luminograph II EM imaging system (ATTO). Protein loading and transfer were assessed by Ponceau S staining. The co-immunoprecipitation experiment was performed independently three times with similar results.

**Fig. S1.**
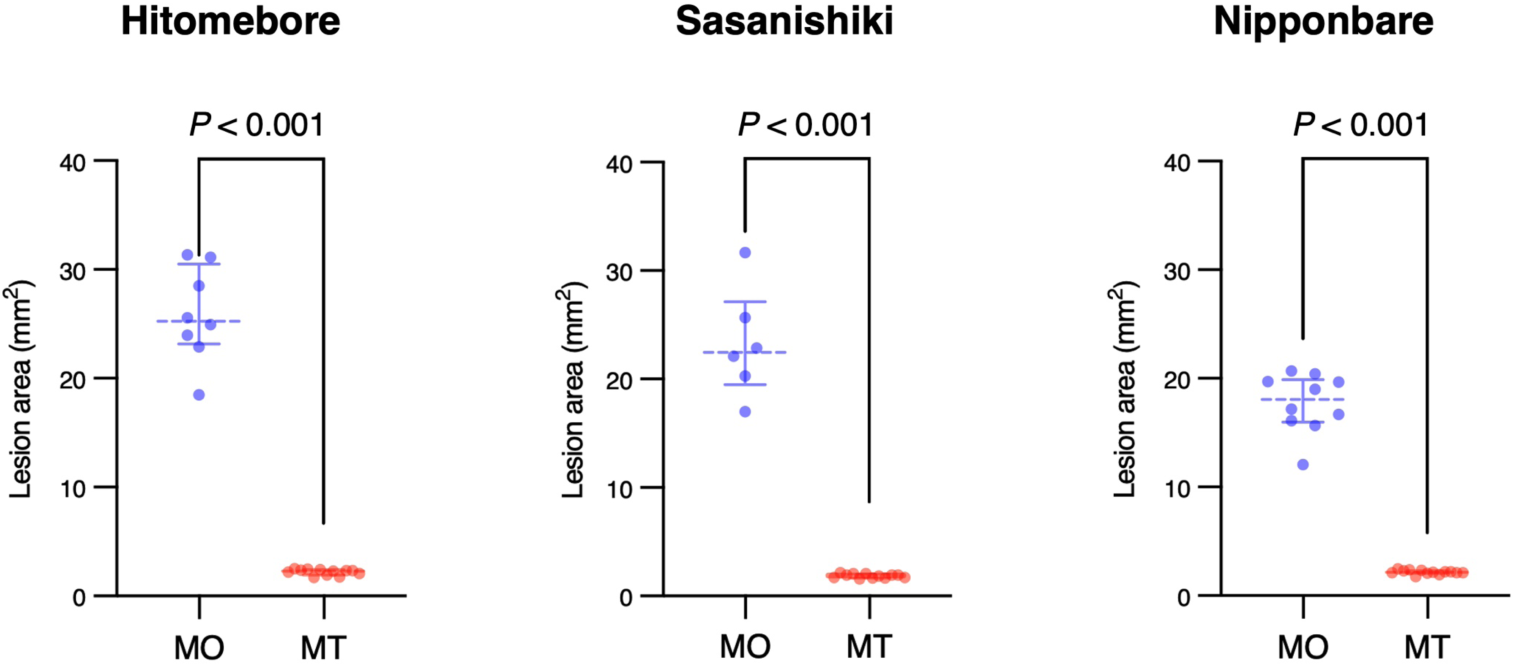
Host specificity of MO and MT isolates on rice. Lesion areas were measured on leaves of three *Oryza sativa* cultivars (Hitomebore, Sasanishiki, and Nipponbare) inoculated with the rice-infecting MO isolate Ao92-06-2 or the wheat-infecting MT isolate Br48. Each dot represents an individual plant. Horizontal lines indicate the median, and error bars indicate the interquartile range. Differences between Ao92-06-2 and Br48 were assessed separately for each cultivar using a two-tailed Mann–Whitney U test. *P* values are indicated above the comparisons.

**Fig. S2.**
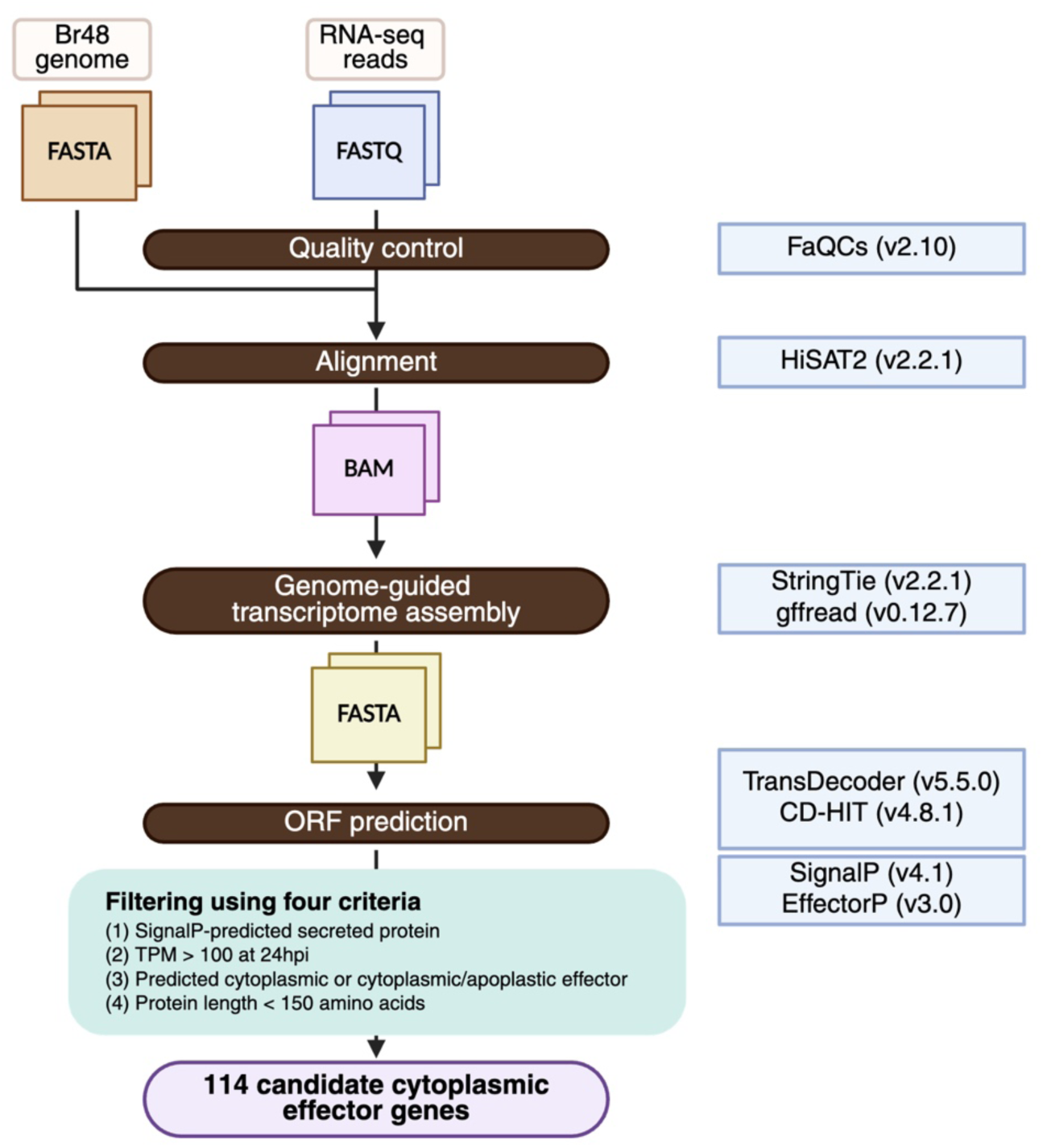
Pipeline for identification of candidate cytoplasmic effector genes from *M. oryzae* pathotype *Triticum* isolate Br48. RNA was extracted from leaves of barley cultivar Nigrate at 24 hours post-inoculation with the MT isolate Br48. RNA-seq reads were processed through the following steps: (1) quality control using FaQCs v2.10; (2) alignment to the Br48 genome using HISAT2 v2.2.1; (3) genome-guided transcriptome assembly using StringTie v2.2.1, followed by transcript sequence extraction using gffread v0.12.7, yielding 7,774 transcripts longer than 200 bp; (4) open reading frame prediction using TransDecoder v5.5.0 and removal of exact duplicate protein sequences using CD-HIT v4.8.1; and (5) selection of candidates encoding SignalP-predicted secreted proteins, showing expression of TPM > 100 at 24 hpi, classified as cytoplasmic or cytoplasmic/apoplastic effectors by EffectorP v3.0, and encoding proteins shorter than 150 amino acids. This pipeline identified 114 candidate cytoplasmic effector genes.

**Fig. S3.**
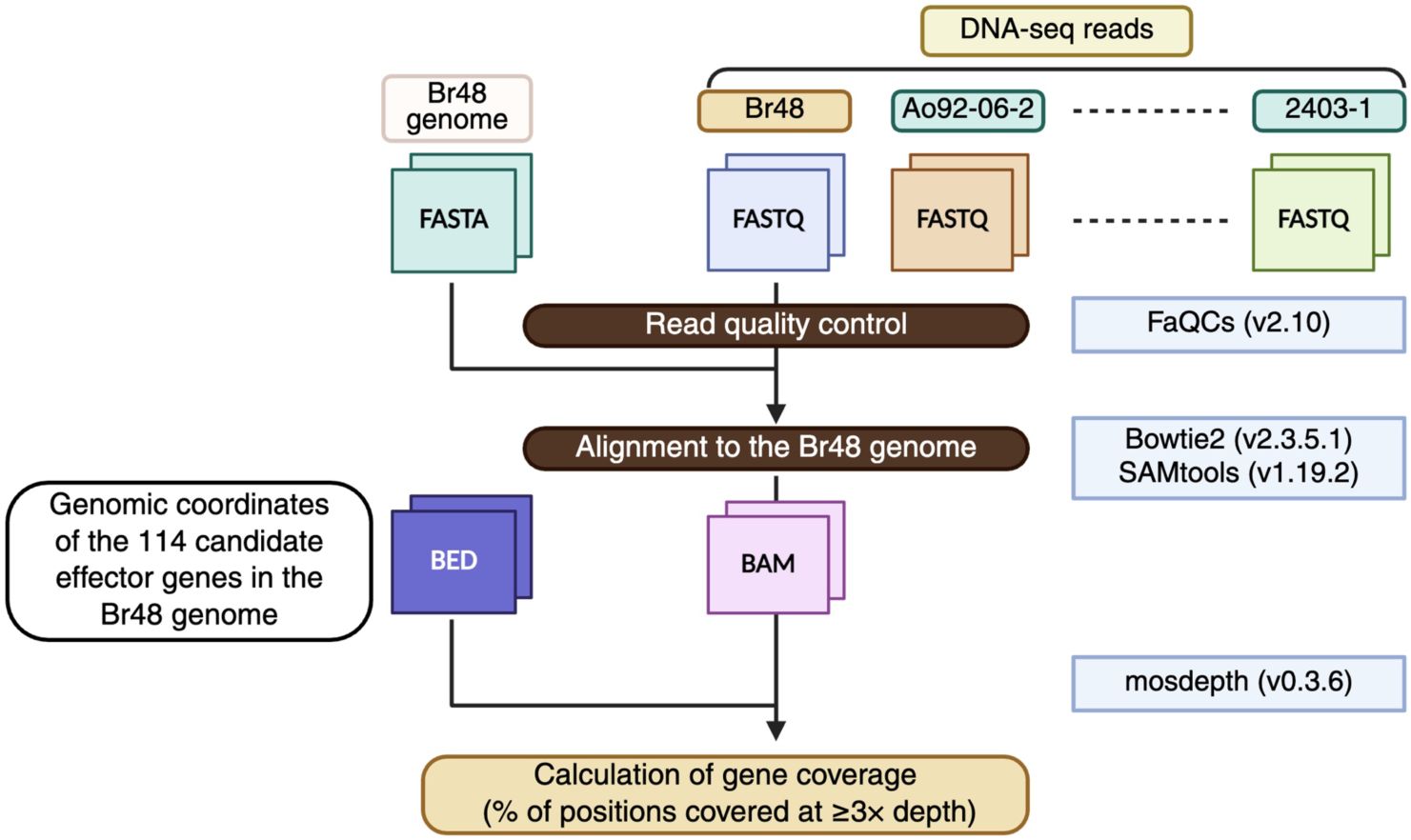
Pipeline for estimating candidate effector gene coverage across *M. oryzae* isolates. Whole-genome sequencing reads from the MT isolate Br48 and nine MO isolates were quality-filtered using FaQCs v2.10 and aligned to the Br48 reference genome using Bowtie2 v2.3.5.1. The resulting alignments were processed into sorted and indexed BAM files. A BED file containing the genomic coordinates of the 114 candidate effector genes in the Br48 genome was used for coverage analysis with mosdepth v0.3.6. For each candidate effector gene, gene coverage was calculated as the percentage of nucleotide positions supported by a read depth of ≥3×.

**Fig. S4.**
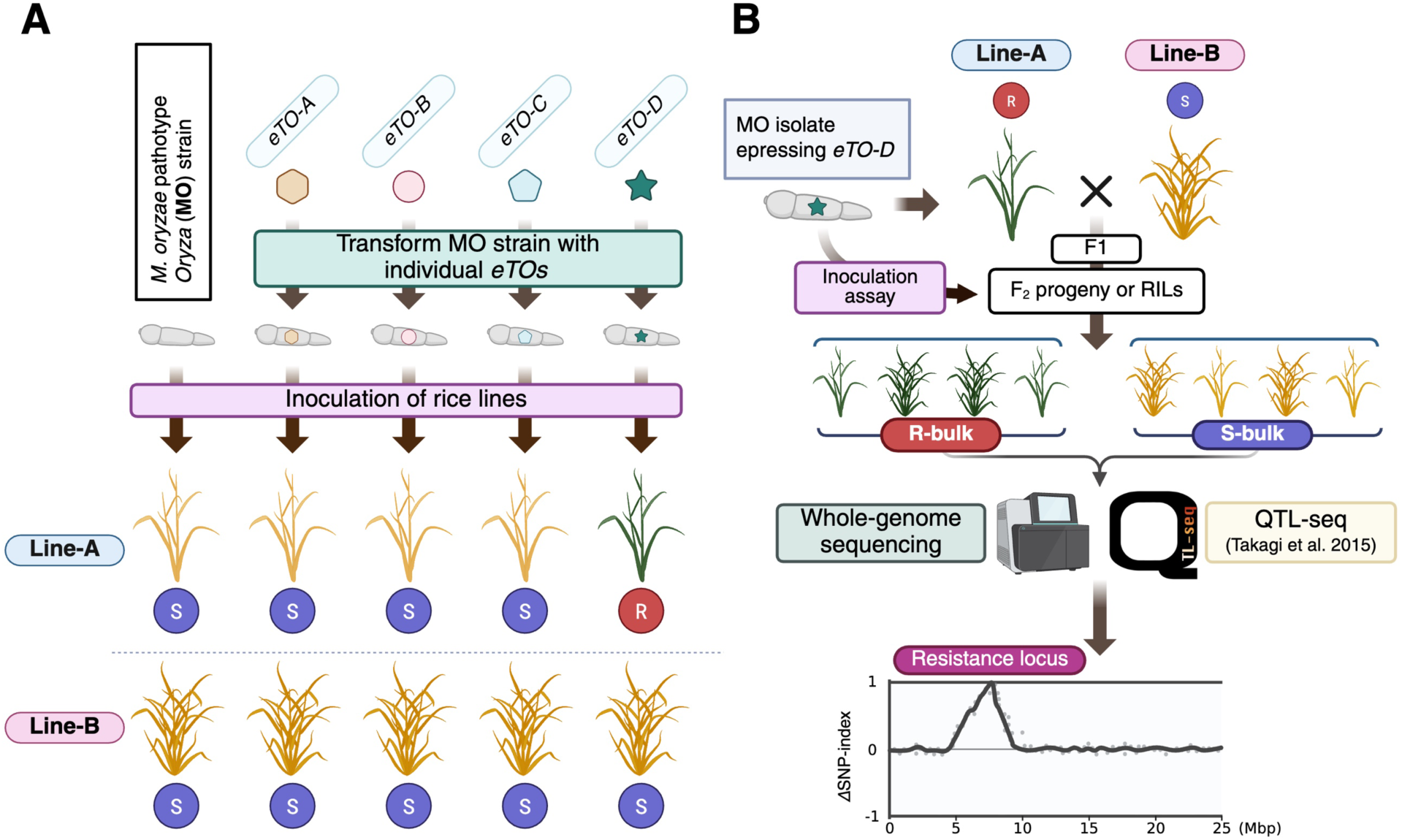
Pipeline for systematic identification of MT-specific AVR genes and their cognate rice resistance loci. **(A)** Functional screen of MT-specific effector genes. Individual *eTO* genes are introduced into the rice-infecting MO isolate Ao92-06-2, and the resulting transformants are inoculated onto multiple rice lines. An *eTO* that specifically triggers resistance in a rice line is identified as a candidate AVR gene. In the example shown, *eTO-D* triggers resistance in Line-A but not in Line-B. R and S indicate resistant and susceptible responses, respectively. **(B)** Mapping of the resistance locus cognate to the identified AVR gene. The resistant Line-A is crossed with the susceptible Line-B, and F_2_ progeny or recombinant inbred lines (RILs) are inoculated with the Ao92-06-2 transformant expressing *eTO-D*. Resistant and susceptible progeny are pooled separately to generate R-bulks and S-bulks, which are subjected to whole-genome sequencing and QTL-seq analysis. Differences in allele frequencies between the two bulks are used to identify the resistance locus responsible for *eTO-D*-triggered resistance.

**Fig. S5.**
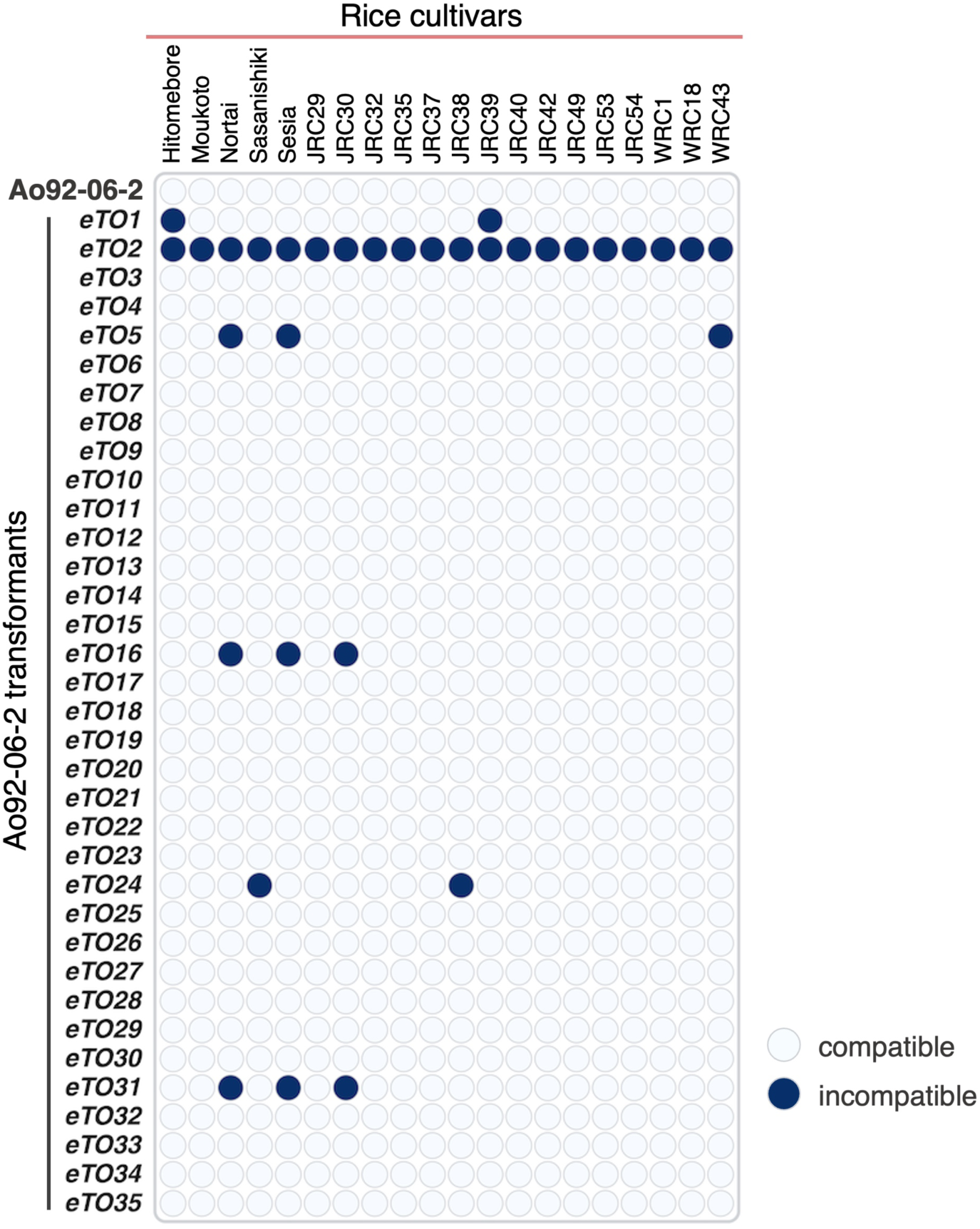
Complete compatibility matrix for functional screening of 35 MT-specific effector genes across 20 rice cultivars. The parental MO isolate Ao92-06-2 and 35 Ao92-06-2 transformants, each expressing an individual *eTO* gene under the control of the *pex22* promoter, were inoculated onto 20 rice cultivars that were susceptible to the parental isolate. Columns indicate rice cultivars (**Data S4**), and rows indicate the parental isolate or transformants expressing individual *eTO* genes. Filled blue circles indicate incompatible interactions (resistance), whereas open circles indicate compatible interactions (susceptibility). The parental Ao92-06-2 isolate was compatible with all cultivars tested. Six *eTOs* (*eTO1, eTO2, eTO5, eTO16, eTO24*, and *eTO31*) triggered resistance in at least one cultivar.

**Fig. S6.**
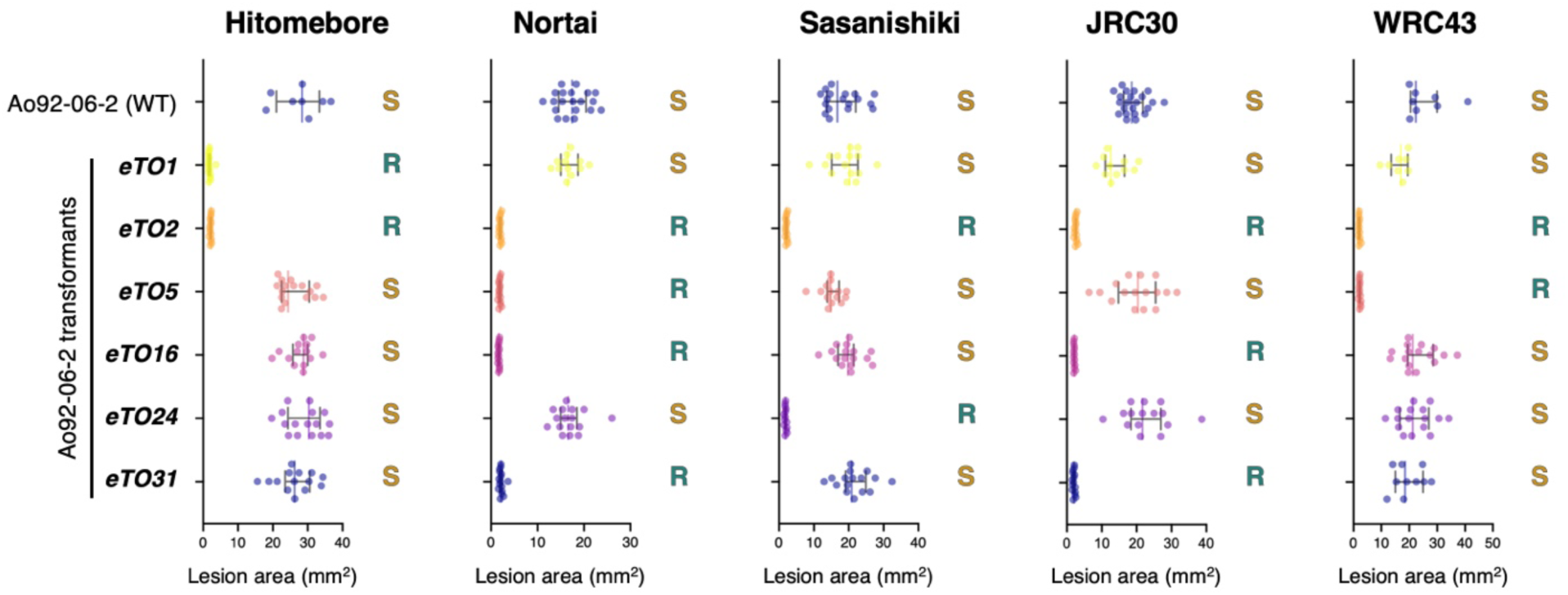
Quantification of lesion areas in rice cultivars inoculated with Ao92-06-2 transformants expressing selected *eTO* genes. Lesion areas were measured in the indicated rice cultivars following inoculation with the parental MO isolate Ao92-06-2 or Ao92-06-2 transformants expressing *eTO1, eTO2, eTO5, eTO16, eTO24,* or *eTO31* under the control of the *pex22* promoter. Representative inoculation phenotypes are shown in **Fig. 2A**. Each dot represents an individual plant. Horizontal lines indicate the median, and error bars indicate the interquartile range. Interactions were classified as resistant (R) when no lesion expansion beyond the inoculation site was observed and as susceptible (S) when lesion expansion occurred. Lesion areas were measured at 10 days post-inoculation.

**Fig. S7.**
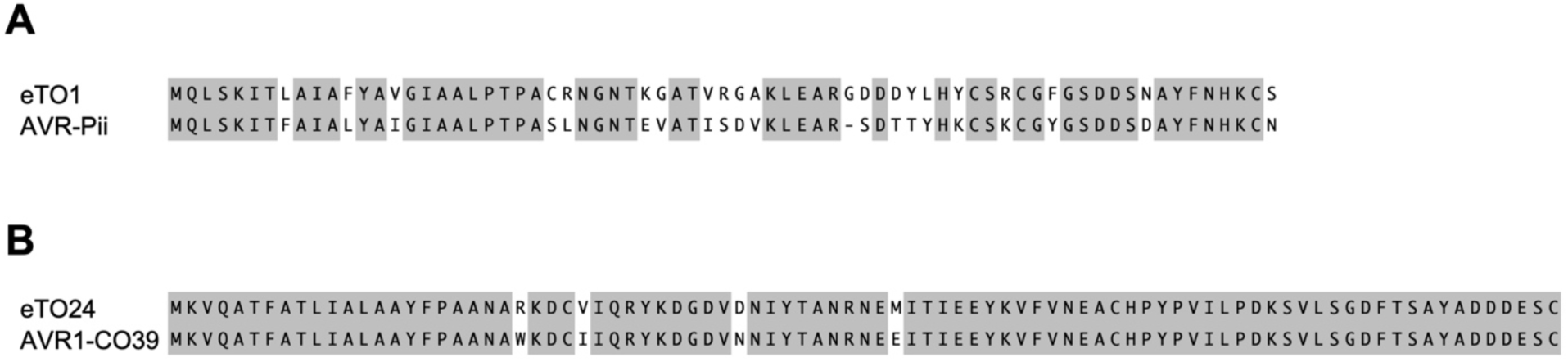
Amino acid sequence alignments eTO1 with AVR-Pii and eTO24 with AVR1-CO39. (A) Pairwise amino acid sequence alignment of eTO1 and AVR-Pii. (B) Pairwise amino acid sequence alignment of eTO24 and AVR1-CO39. Identical amino acid residues are highlighted in gray.

**Fig. S8.**
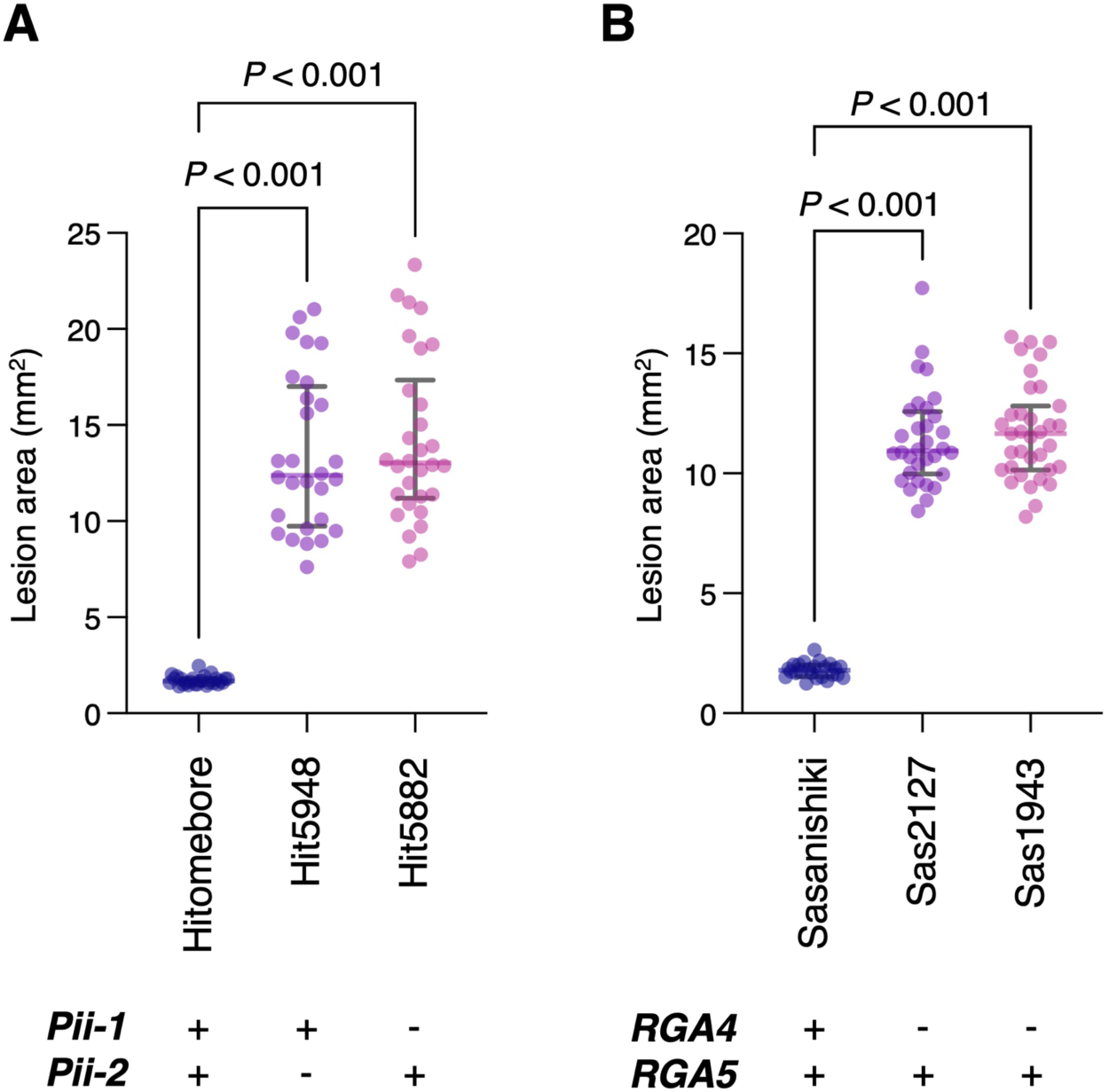
*Pii* and *Pia* are required for *eTO1-* and *eTO24*-triggered resistance, respectively. **(A)** Lesion areas in Hitomebore, which carries the paired NLR genes *Pii-1* and *Pii-2*, and two independent *pii* mutant lines, Hit5948 and Hit5882, following inoculation with Ao92-06-2 expressing *eTO1* under the control of the pex22 promoter. **(B)** Lesion areas in Sasanishiki, which carries the paired NLR genes *RGA4* and *RGA5* at the *Pia* locus, and two independent *rga4* mutant lines, Sas2127 and Sas1943, following inoculation with Ao92-06-2 expressing *eTO24* under the control of the pex22 promoter. Each dot represents an individual plant. Horizontal lines indicate the median, and error bars indicate the interquartile range. Plus and minus symbols indicate the presence or absence of *Pii-1* and *Pii-2* in (A) and *RGA4* and *RGA5* in (B). Statistical significance was assessed using a Kruskal–Wallis test followed by Dunn’s multiple-comparison test. *P* values are indicated above the comparisons.

**Fig. S9.**
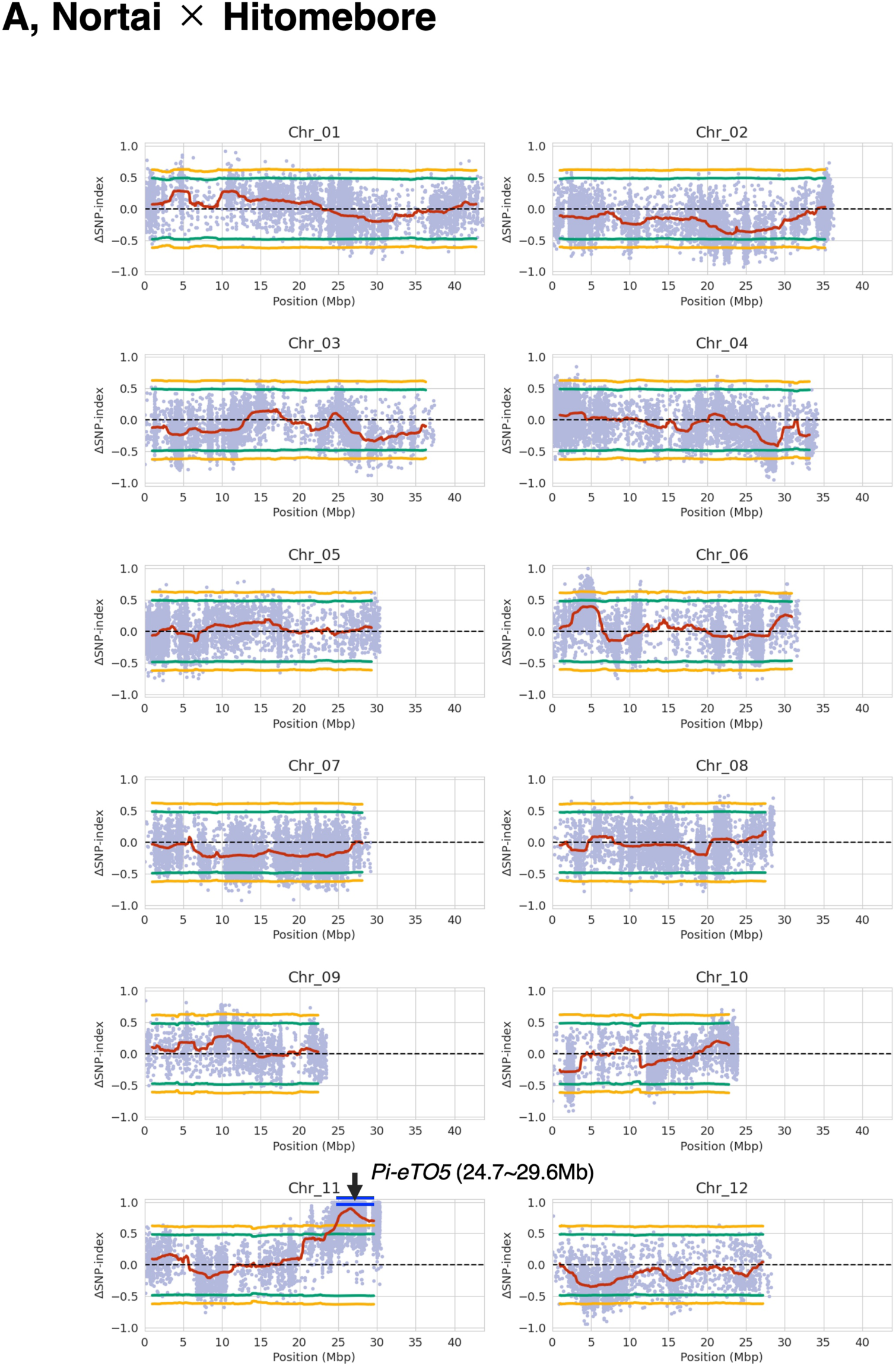
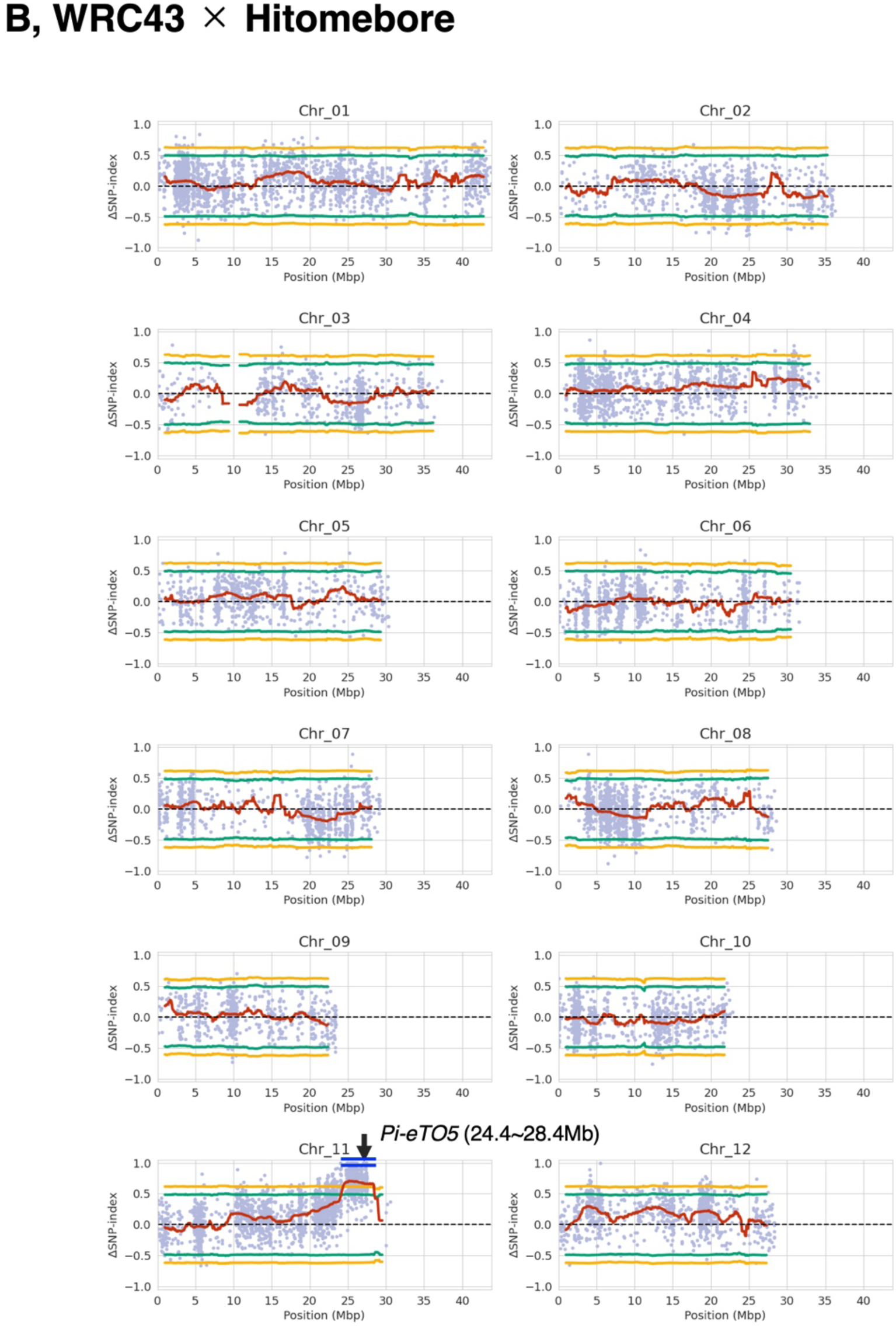
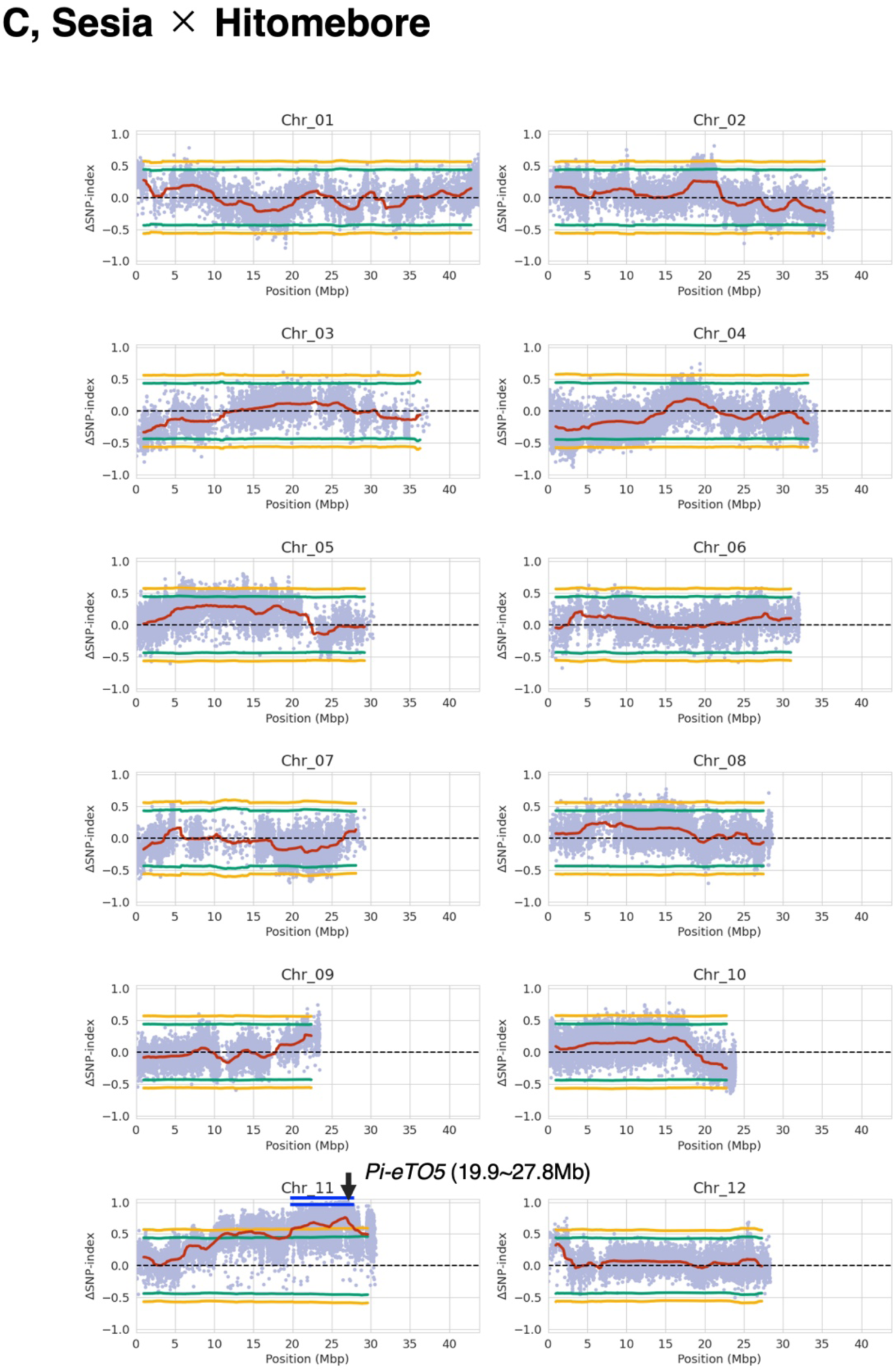
Independent QTL-seq analyses identify an overlapping *Pi-eTO5* interval on chromosome 11. Genome-wide ΔSNP-index profiles obtained from recombinant inbred line (RIL) populations derived from **(A)** Nortai × Hitomebore and **(C)** Sesia × Hitomebore, and from **(B)** an F_2_ population derived from WRC43 × Hitomebore, are shown. QTL-seq analyses were performed using the Nortai genome as the reference. Blue dots indicate ΔSNP-index values for individual SNPs. The red line indicates the sliding-window-smoothed ΔSNP-index. Green and orange lines indicate the 95% and 99% confidence limits, respectively. The three populations identified intervals of 24.7–29.6 Mb, 24.4–28.4 Mb, and 19.9–27.8 Mb on chromosome 11, respectively, defining a common interval of 24.7–27.8 Mb associated with *eTO5-*triggered resistance.

**Fig. S10.**
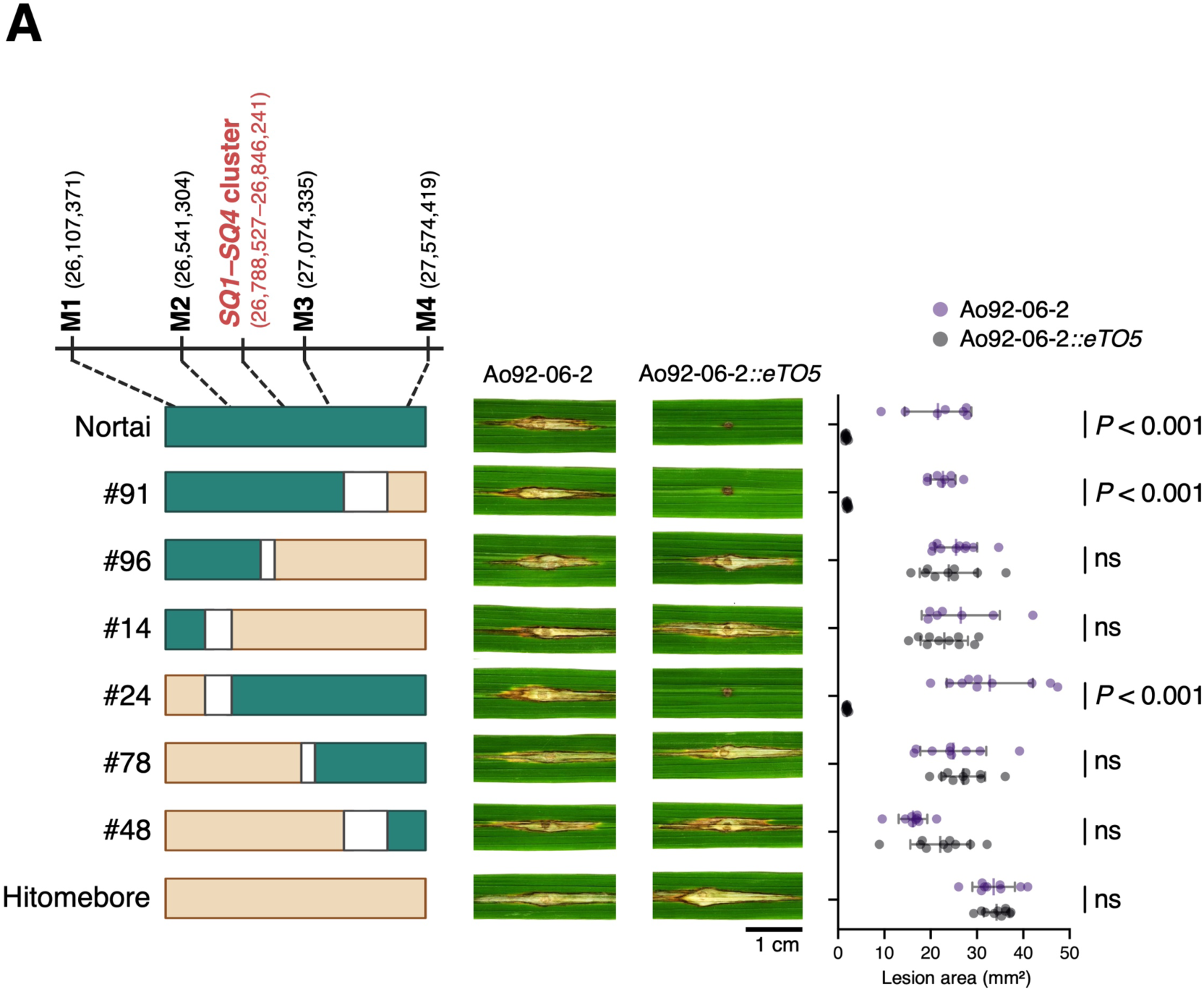
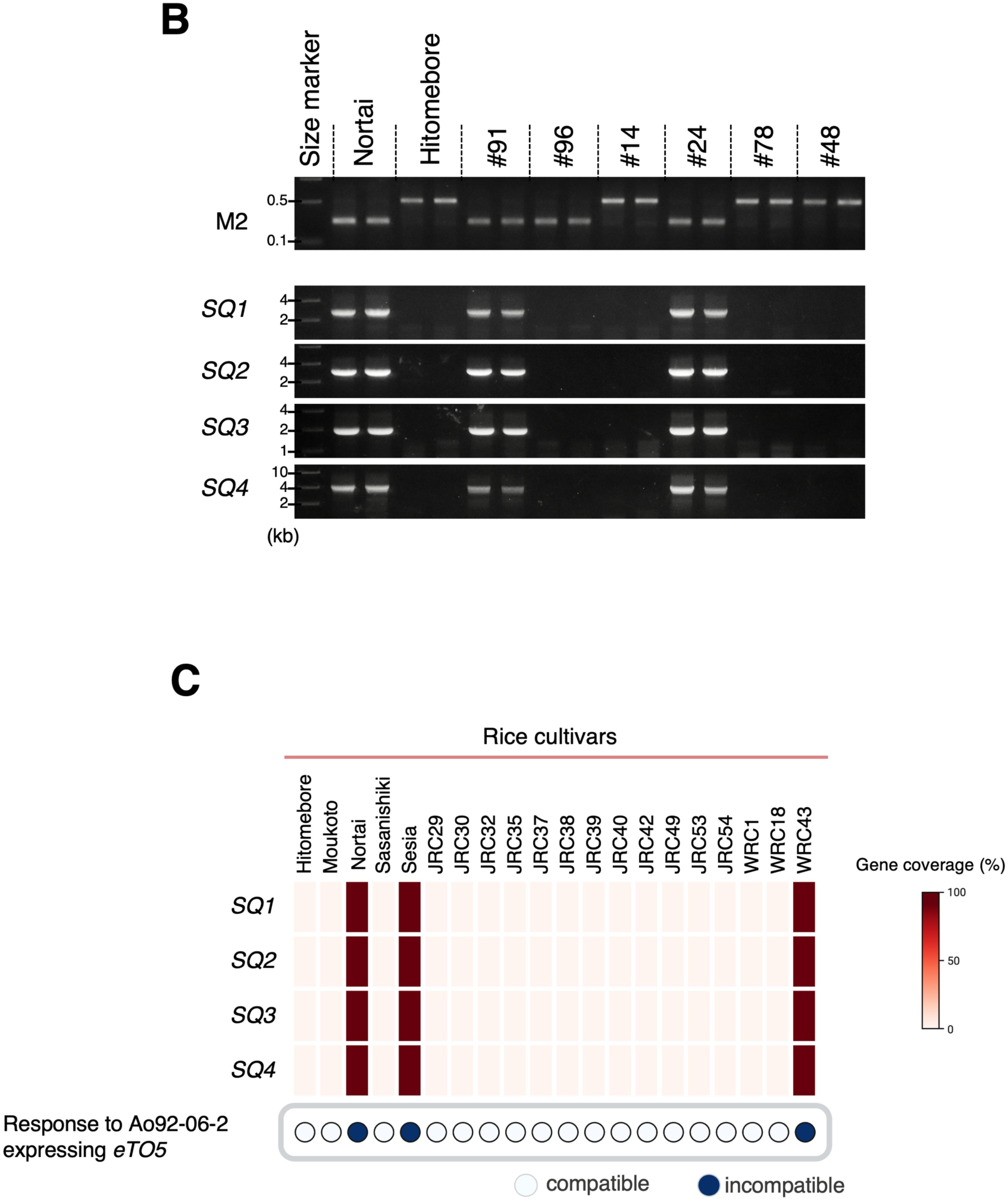
Fine mapping and cultivar distribution of the *Pi-eTO5* NLR quartet. **(A)** Fine mapping of *Pi-eTO5* using recombinant progeny derived from the Nortai×Hitomebore cross. Recombinant haplotypes between markers M1 and M4 are shown together with representative punch-inoculation phenotypes and lesion areas following inoculation with Ao92-06-2 or Ao92-06-2 expressing *eTO5*. Horizontal lines in the right graph indicate the median, and error bars indicate the interquartile range. Differences between Ao92-06-2 and Ao92-06-2 expressing *eTO5* were assessed within each rice line using two-way ANOVA followed by Šídák’s multiple-comparison test. *P* values are indicated; ns, not significant. The interval delimited by recombinants #91 and #24 spans approximately 0.53 Mb and contains the *SQ1*–*SQ4* gene cluster. Scale bar, 1 cm**. (B)** PCR-based genotyping of marker M2 and *SQ1*–*SQ4* in the parental lines and recombinant progeny shown in **(A)**. **(C)** Gene coverage of *SQ1*, *SQ2*, *SQ3*, and *SQ4* across the 20 rice cultivars used for functional screen. Gene coverage was calculated as the percentage of nucleotide positions supported by a read depth of ≥ 3×. The compatibility response to Ao92-06-2 expressing *eTO5* is shown below. Filled blue circles indicate incompatible interactions, whereas open circles indicate compatible interactions. High gene coverage of all four NLR genes was observed in Nortai, Sesia, and WRC43, the three cultivars that exhibited *eTO5*-triggered resistance.

**Fig. S11.**
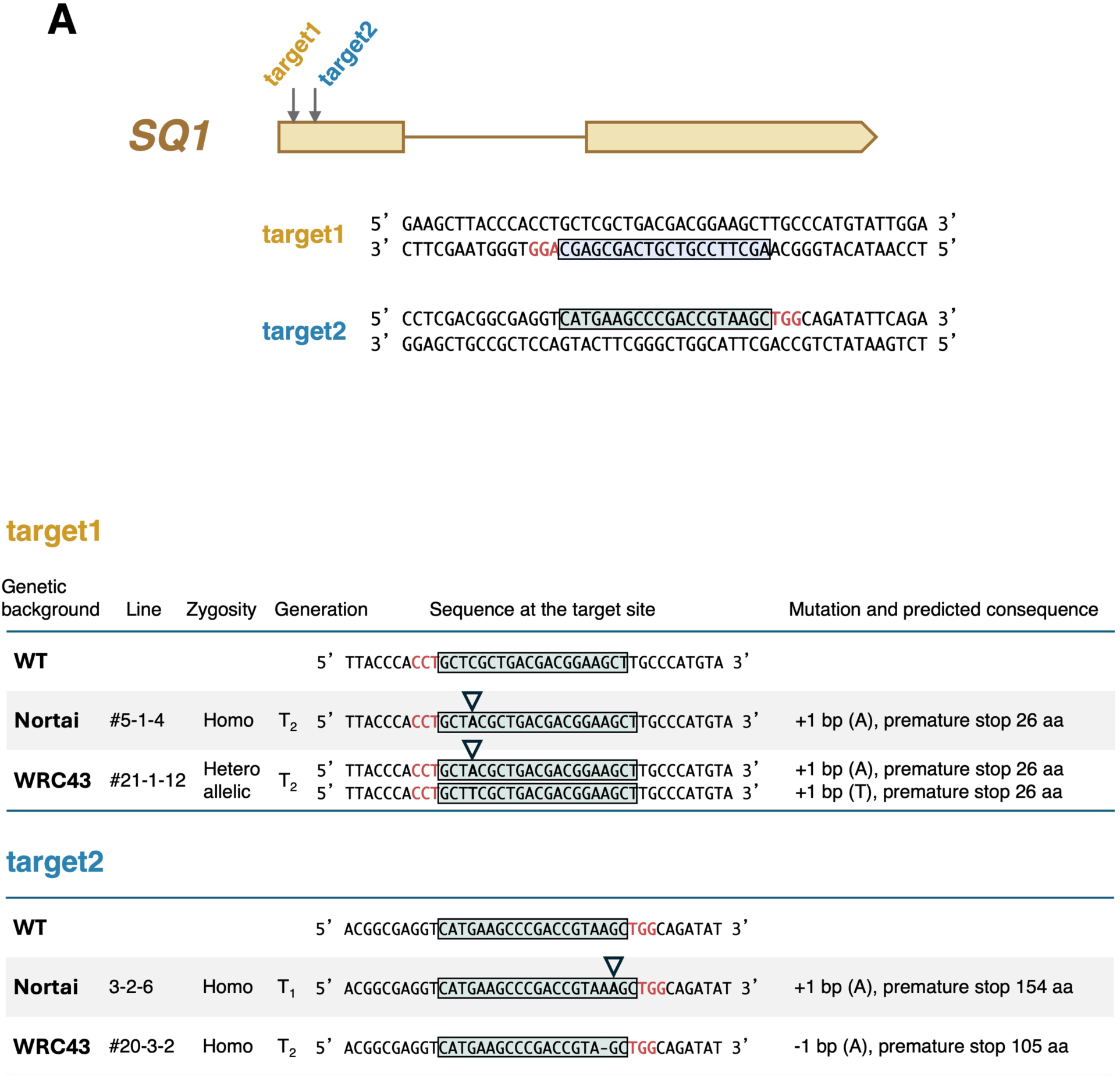
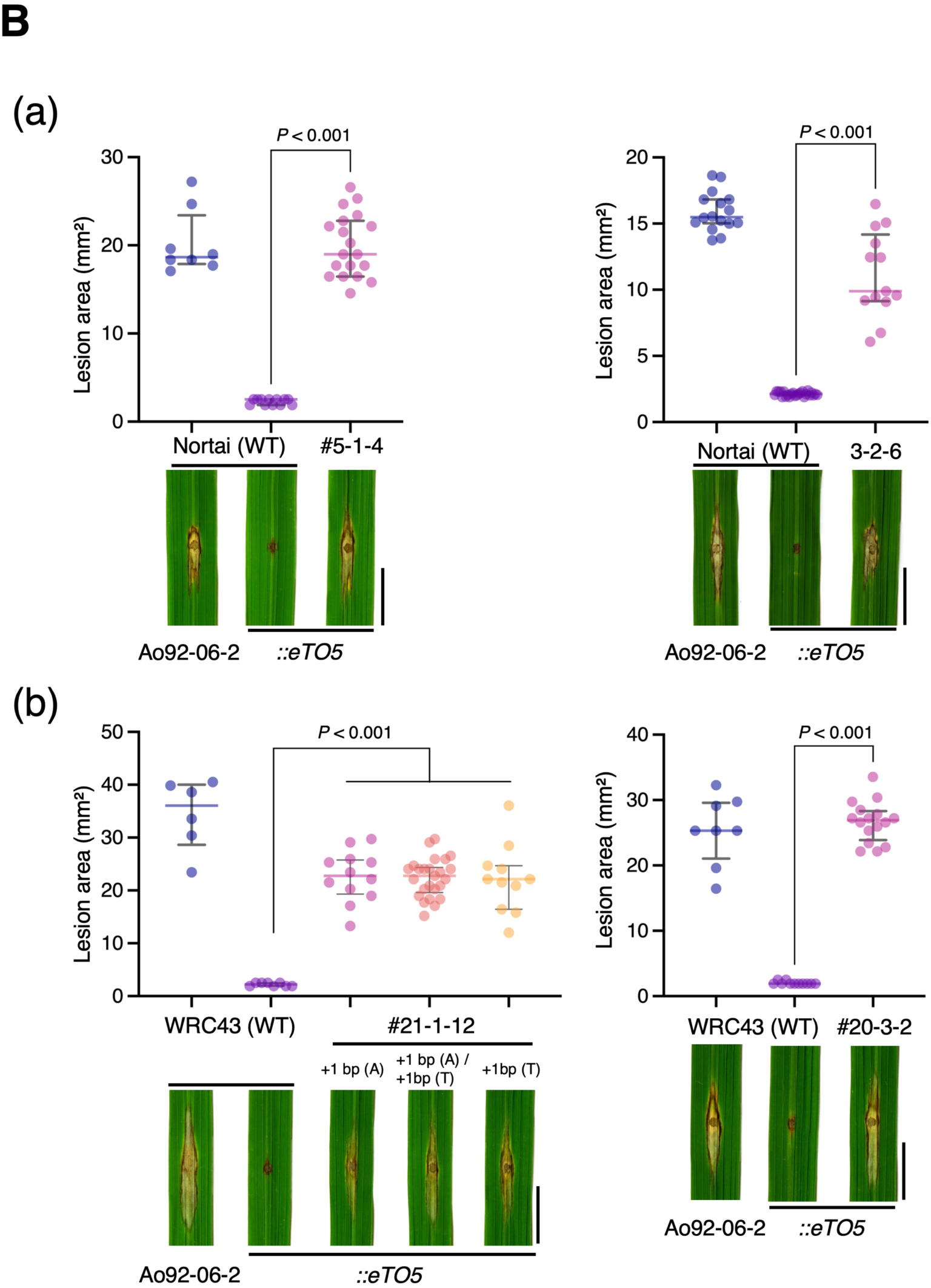
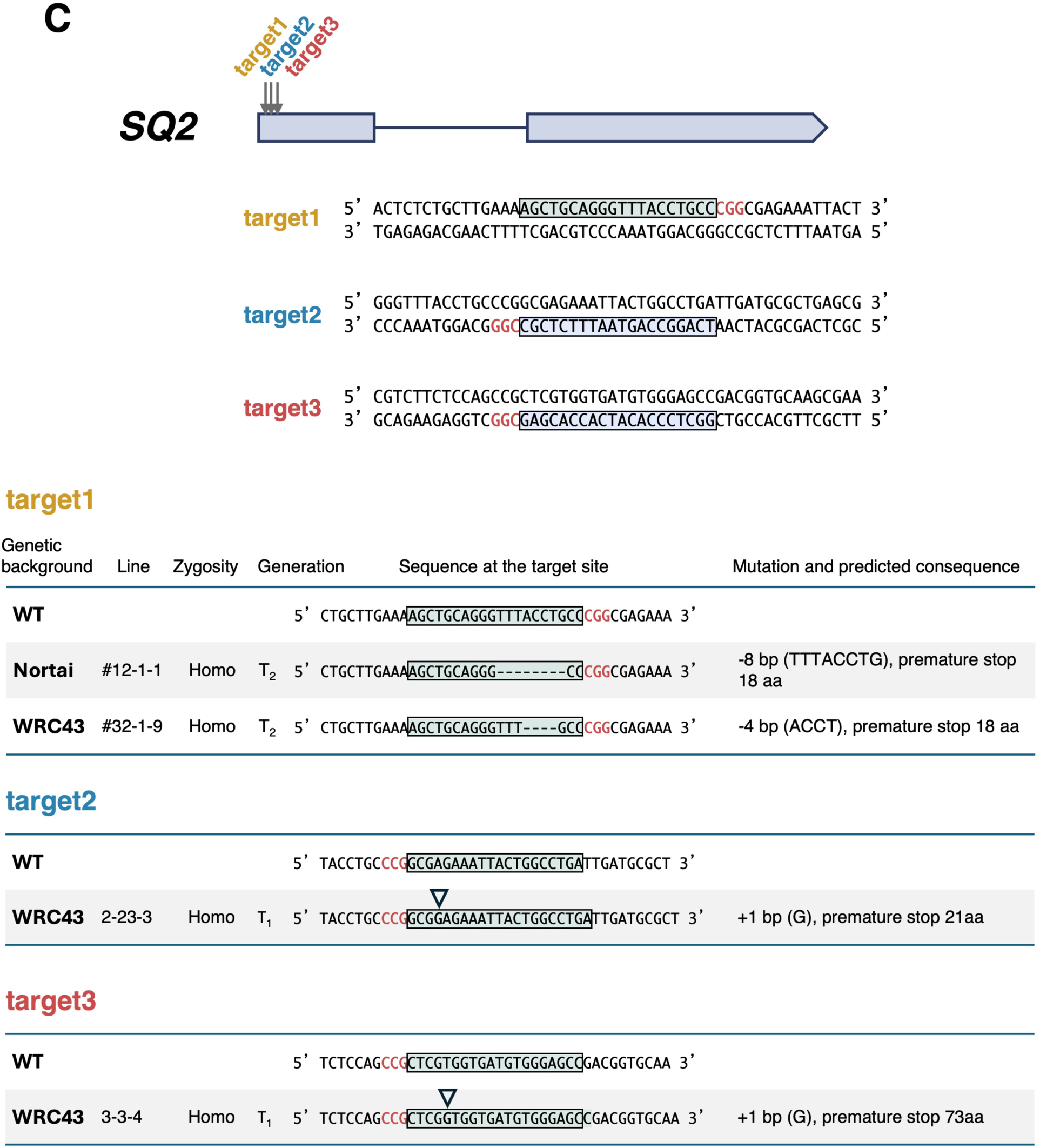
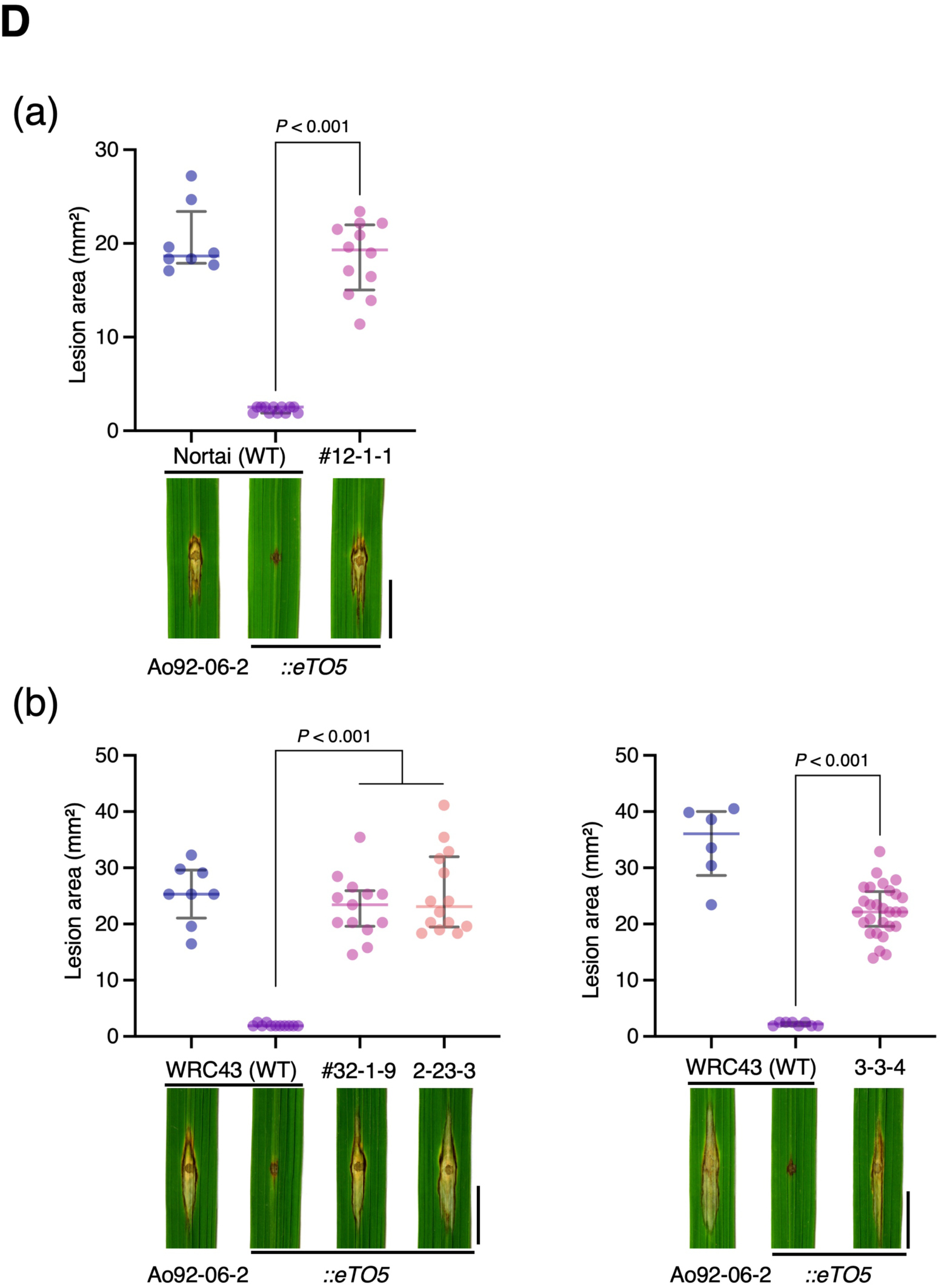
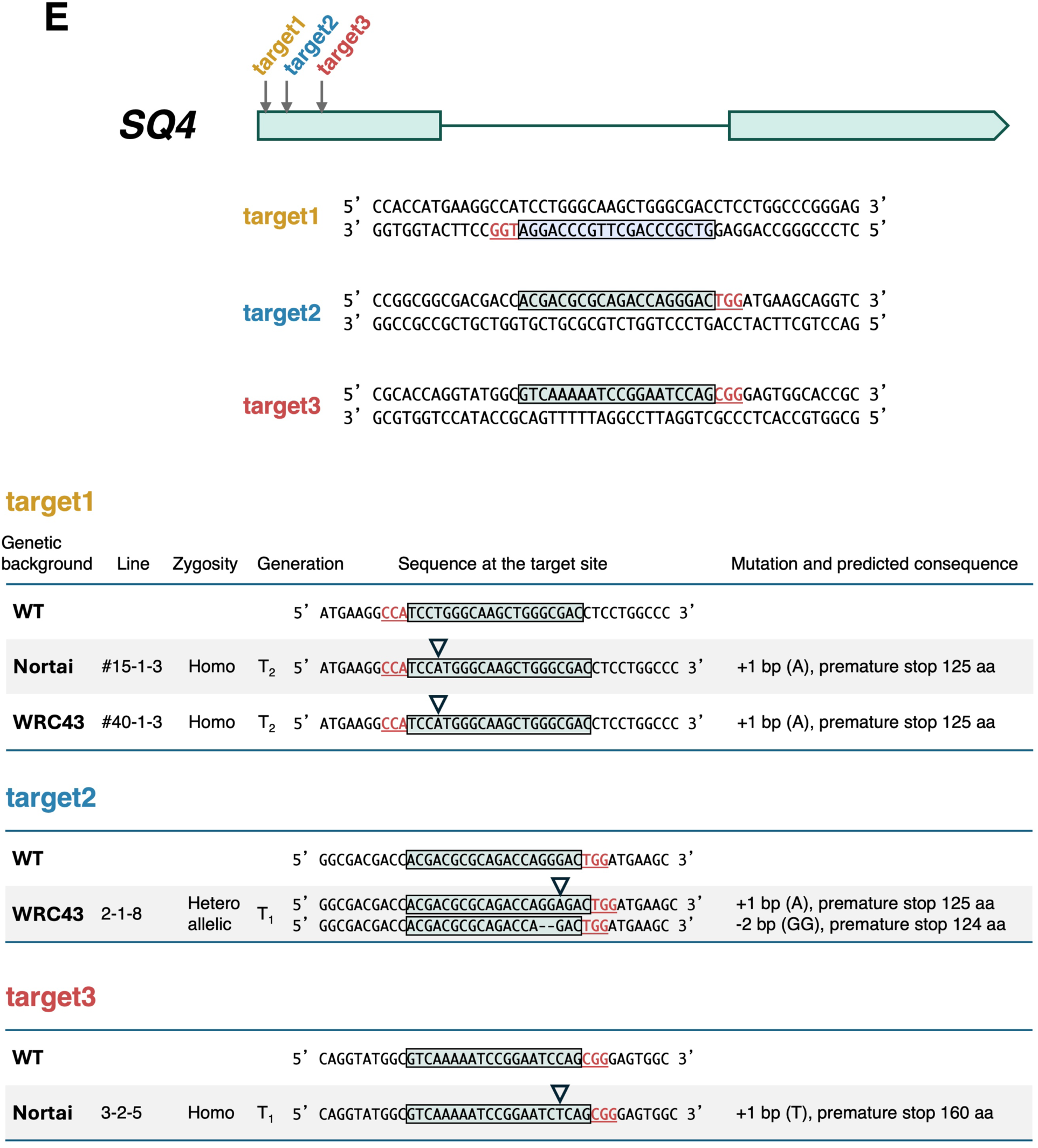
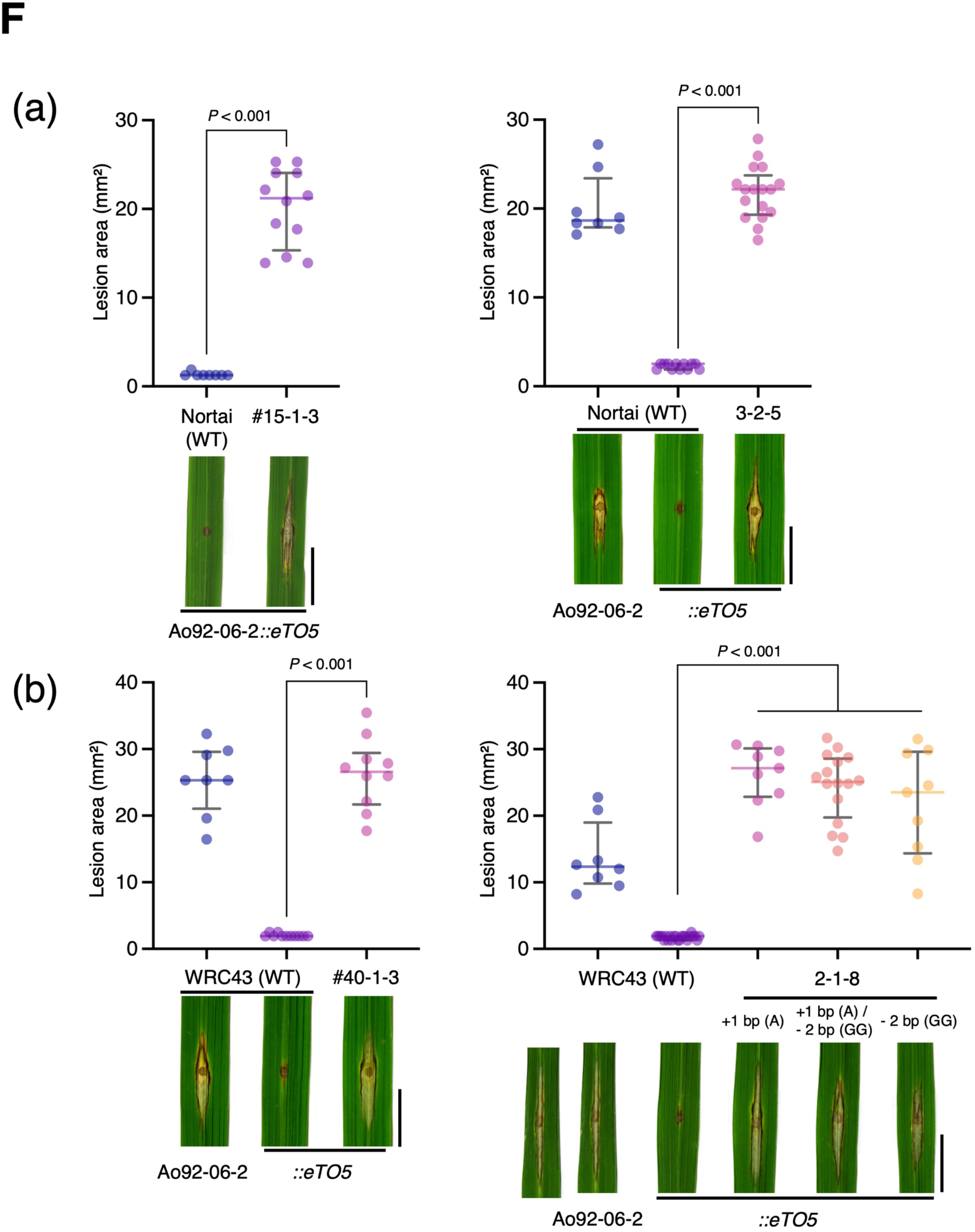
CRISPR-Cas9-mediated disruption of *SQ1*, *SQ2*, and *SQ4* abolishes *eTO5*-triggered resistance. (A, C, **E)** CRISPR-Cas9 target sites and mutations introduced into *SQ1*, *SQ2*, and *SQ4*, respectively. Gene structures and target positions are shown above the aligned wild-type and mutant sequences. Protospacer sequences are boxed, and protospacer-adjacent motifs are shown in red. Insertions are indicated by triangles, and deletions by dashes. The predicted consequences of each mutation, including premature termination and the resulting protein length, are indicated on the right. Homo, homozygous; Heteroallelic, carrying two distinct mutant alleles; T_1_ and T_2_ indicate plant generations. **(B, D, F)** Lesion areas and representative punch-inoculation phenotypes of wild-type and mutant lines for *SQ1*, *SQ2*, and *SQ4*, respectively, in the (a) Nortai and (b) WRC43 backgrounds. Plants were inoculated with the parental Ao92-06-2 isolate or Ao92-06-2 expressing *eTO5*. Each dot represents an individual plant. Horizontal lines indicate the median, and error bars indicate the interquartile range. Statistical significance was assessed using a two-tailed Mann–Whitney U test. P values are indicated above the comparisons. Scale bars, 1 cm.

**Fig. S12.**
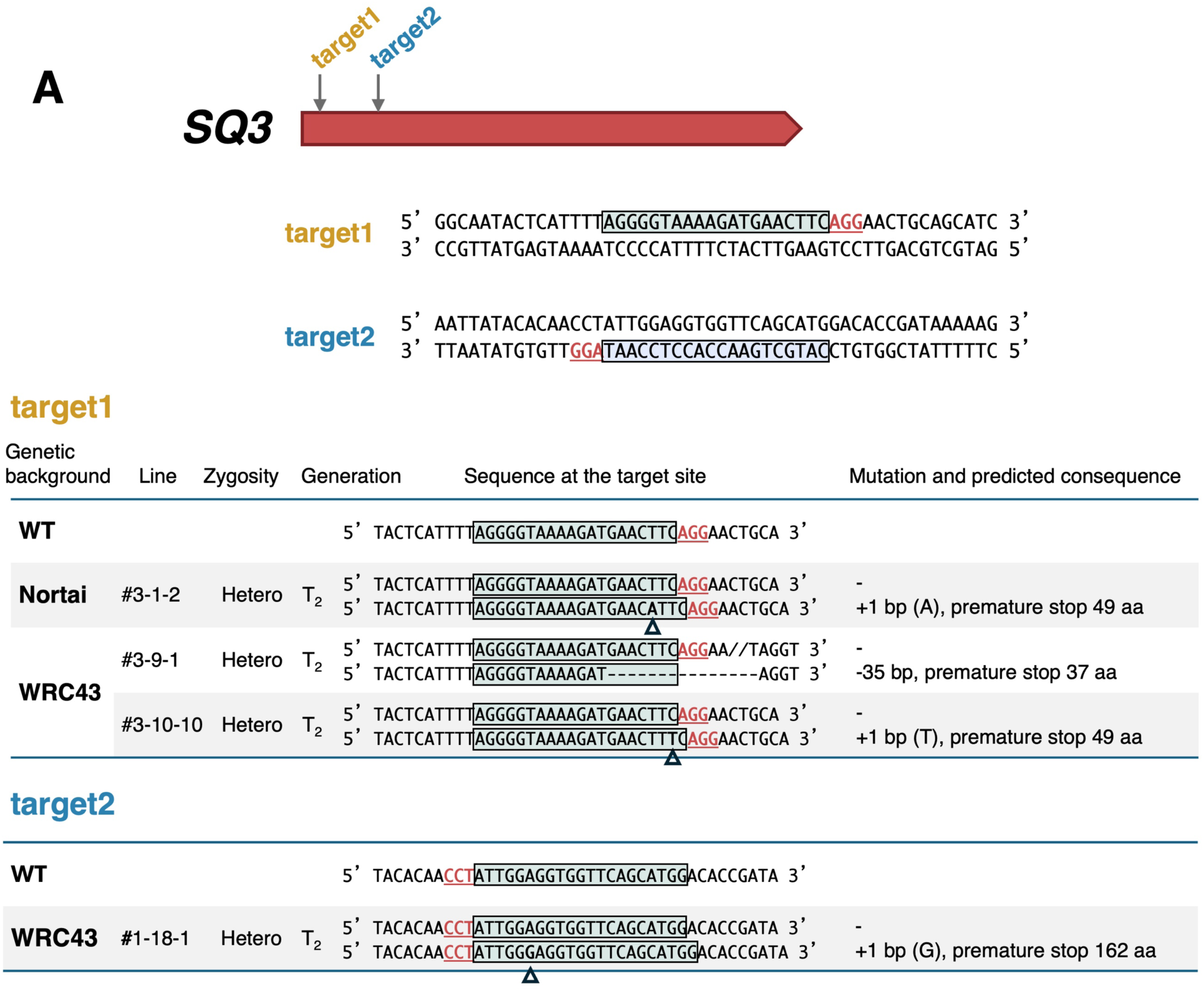
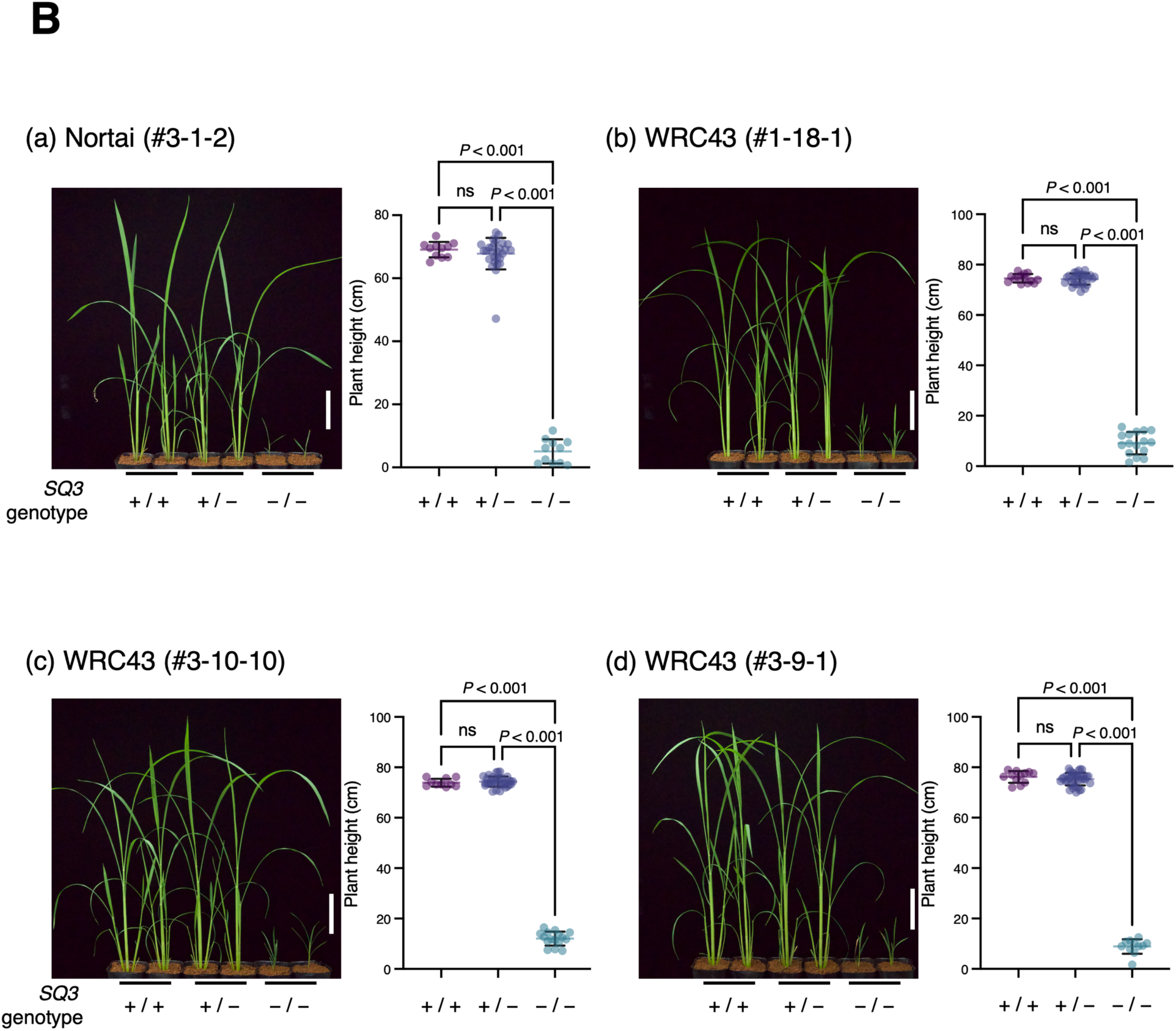
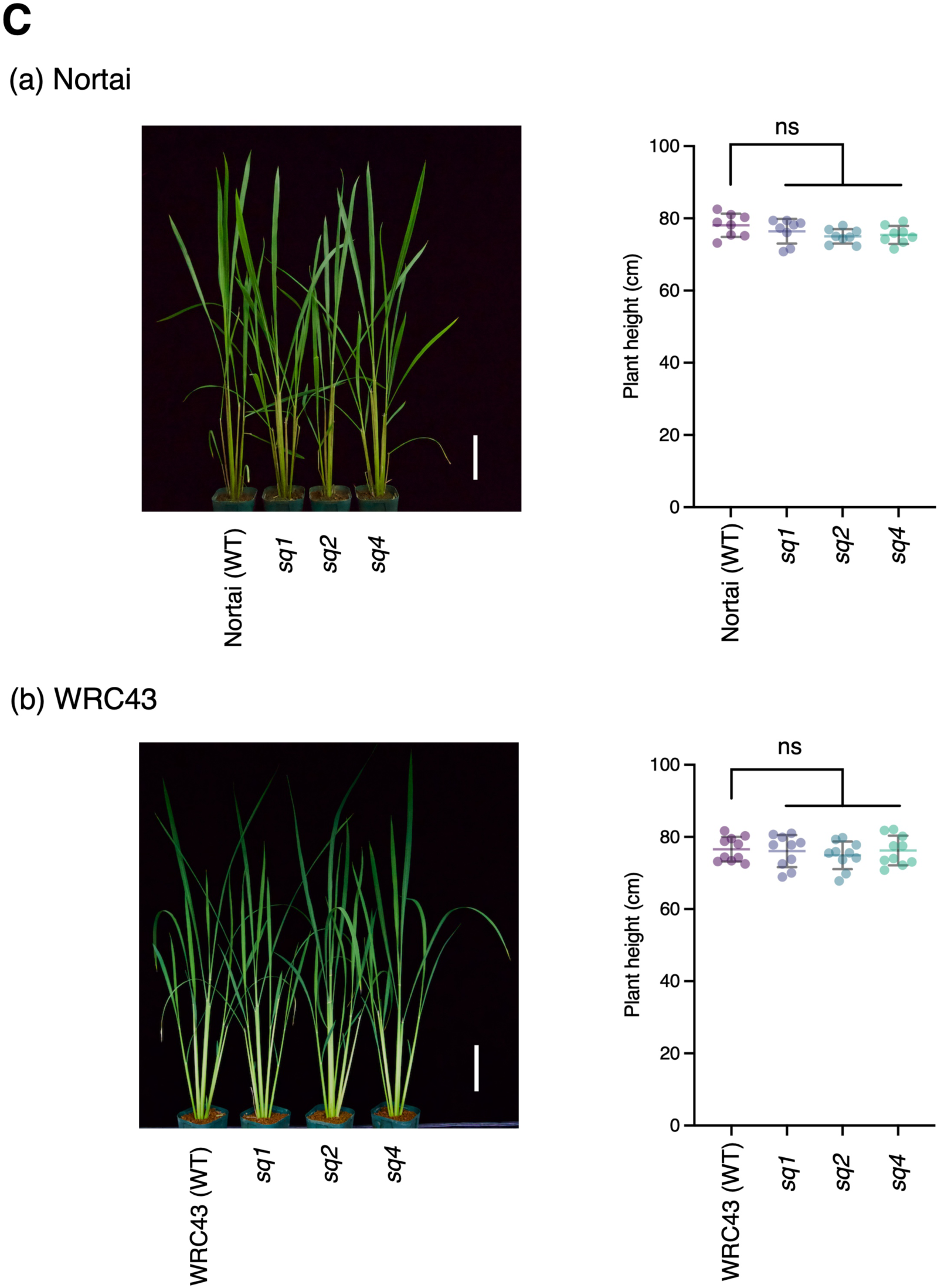
Independent disruption of *SQ3* causes severe dwarfism, whereas disruption of *SQ1*, *SQ2*, or *SQ4* does not affect plant growth. (A) CRISPR-Cas9 target sites and mutations introduced into *SQ3*. The gene structure and target positions are shown above the aligned wild-type and mutant sequences. Protospacer sequences are boxed, and protospacer-adjacent motifs are shown in red. Insertions are indicated by triangles and deletions by dashes. **(B)** Growth phenotypes of segregating T_2_ progeny derived from independent T_1_ plants heterozygous for *SQ3* mutations in the (a) Nortai and (b-d) WRC43 backgrounds. Representative plants and plant-height measurements are shown for *SQ3/SQ3* (+/+), *SQ3/sq3* (+/−), and *sq3/sq3* (−/−) progeny. Homozygous *sq3* plants displayed severe dwarfism, whereas wild-type and heterozygous siblings showed normal growth. Statistical significance was assessed separately for each edited line using one-way ANOVA followed by Tukey’s multiple-comparison test. *P* values are indicated above the comparisons; ns, not significant. Scale bars, 10 cm. **(C)** Representative growth phenotypes and plant-height measurements of *SQ1*, *SQ2*, and *SQ4* knockout lines in the (a) Nortai and (b) WRC43 backgrounds. The Nortai lines were *sq1* (#5-1-4), *sq2* (#12-1-1), and *sq4* (#15-1-3), and the WRC43 lines were *sq1* (#20-3-2), *sq2* (#32-1-9), and *sq4* (#40-1-3). Statistical significance was assessed separately for each genetic background using one-way ANOVA followed by Dunnett’s multiple-comparison test, with each knockout line compared with the corresponding wild type. No significant differences in plant height were detected. ns, not significant. Scale bars, 10 cm.

**Fig. S13.**
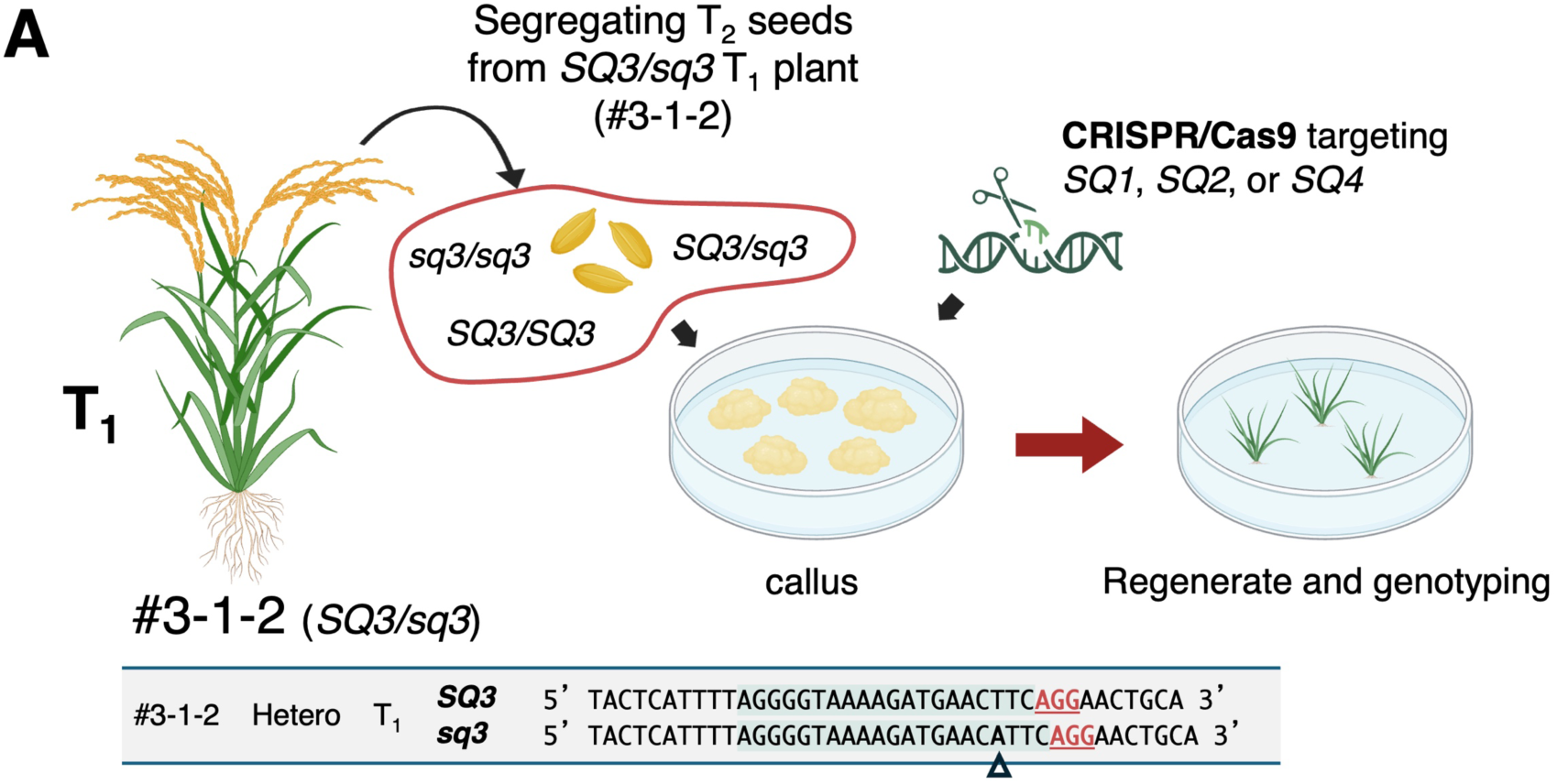
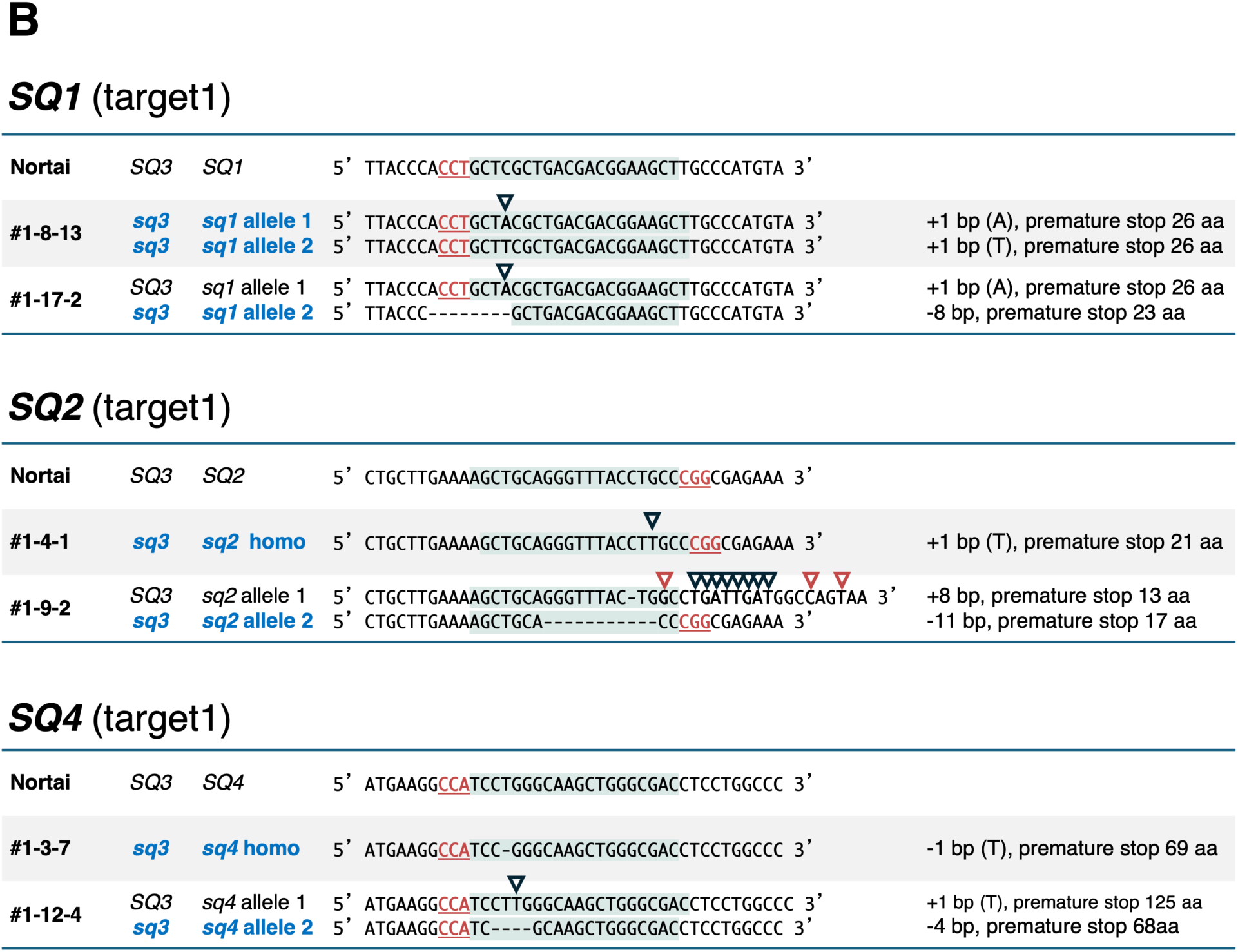
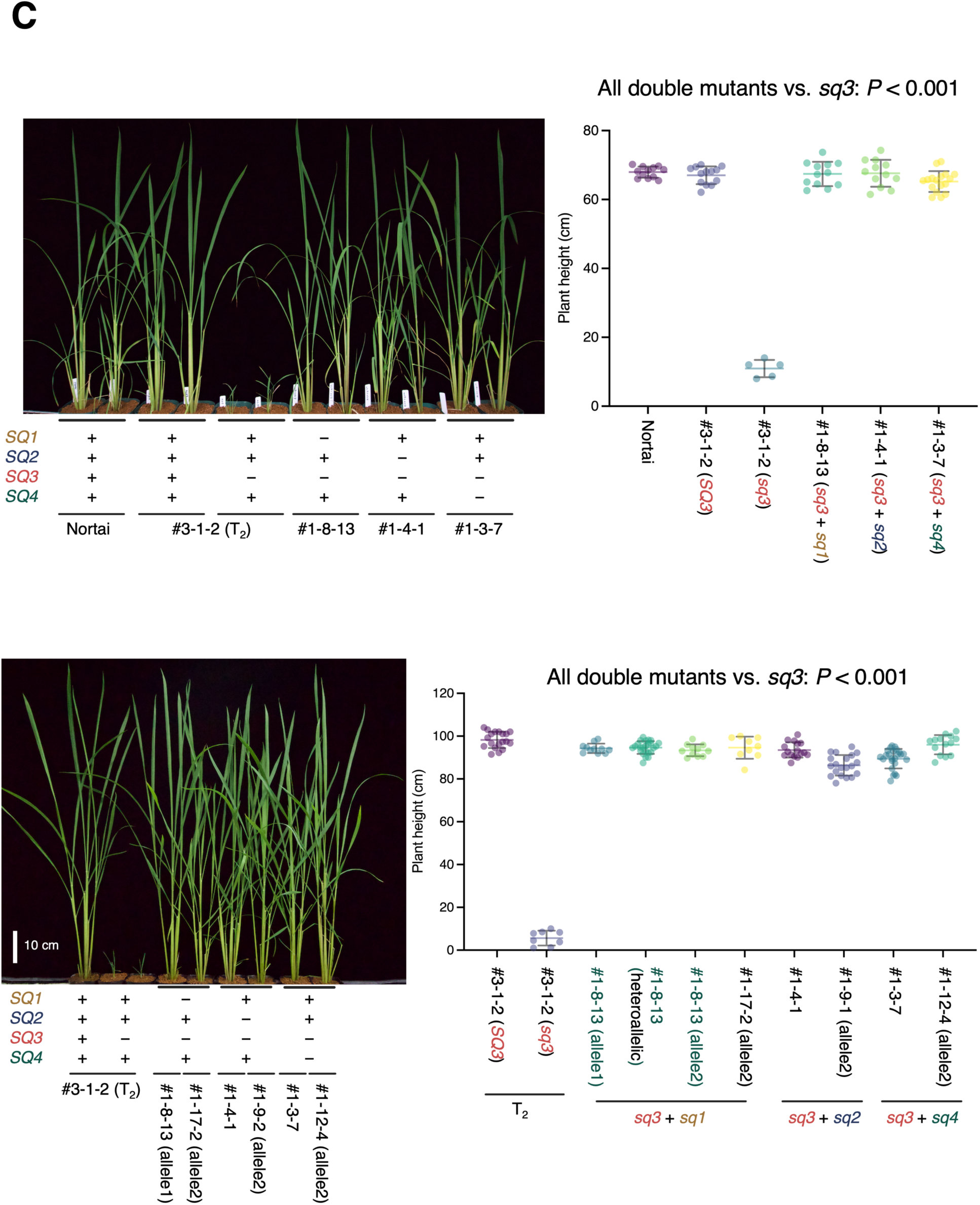
Additional disruption of *SQ1*, *SQ2*, or *SQ4* suppresses the severe dwarf phenotype of *sq3* plants. **(A)** Strategy used to generate double-knockout lines in the *sq3/sq3* background. Segregating T_2_ seeds derived from the T_1_ *SQ3/sq3* line #3-1-2 were used for callus induction. CRISPR-Cas9 constructs targeting *SQ1*, *SQ2*, or *SQ4* were introduced into the calli, followed by plant regeneration and genotyping. The *SQ3* genotype and target-site sequences of line #3-1-2 are shown below. **(B)** Mutations introduced into *SQ1*, *SQ2*, or *SQ4* in the *sq3/sq3* background. Wild-type and edited target-site sequences are shown. Protospacer sequences are boxed, and protospacer-adjacent motifs are shown in red. Insertions are indicated by triangles and deletions by dashes. The predicted consequences of each mutation, including premature termination and the resulting protein length, are indicated on the right. **(C)** Growth phenotypes and plant-height measurements of *sq3/sq3* single mutants and double mutants additionally disrupted for *SQ1*, *SQ2*, or *SQ4*. The upper panels correspond to the initial experiment shown in **Fig. 3E**, whereas the lower panels show an independent biological replicate using additional edited lines. The *sq3/sq3* plants displayed severe dwarfism, whereas additional disruption of *SQ1*, *SQ2*, or *SQ4* restored plant growth in both experiments. Each dot represents an individual plant. Horizontal lines indicate the mean, and error bars indicate the standard deviation. Statistical significance was assessed separately for each experiment using one-way ANOVA followed by Dunnett’s multiple-comparison test, with the *sq3* single mutant as the reference group. *P* values are indicated above the comparisons. Scale bars, 10 cm.

**Fig. S14.**
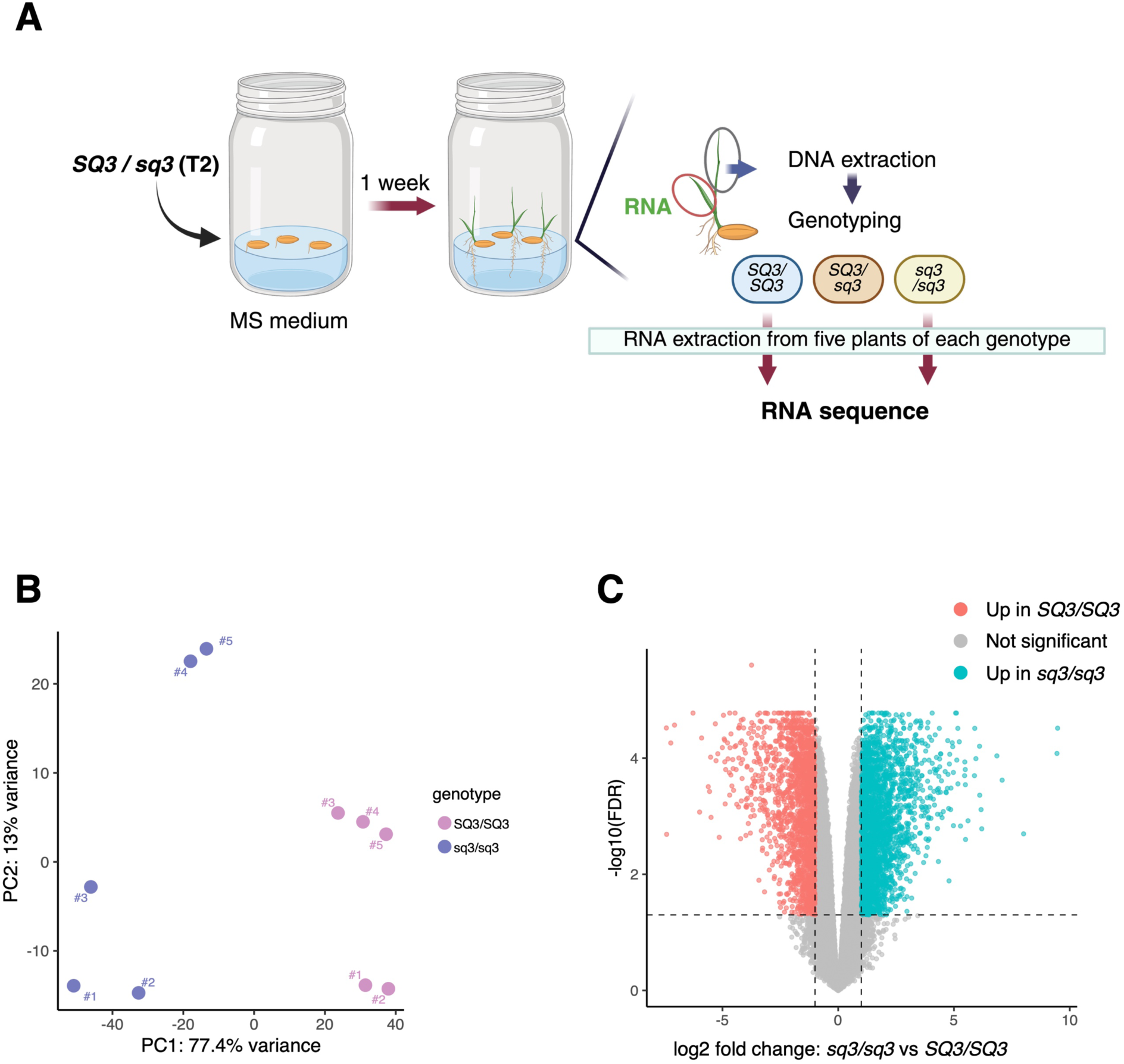
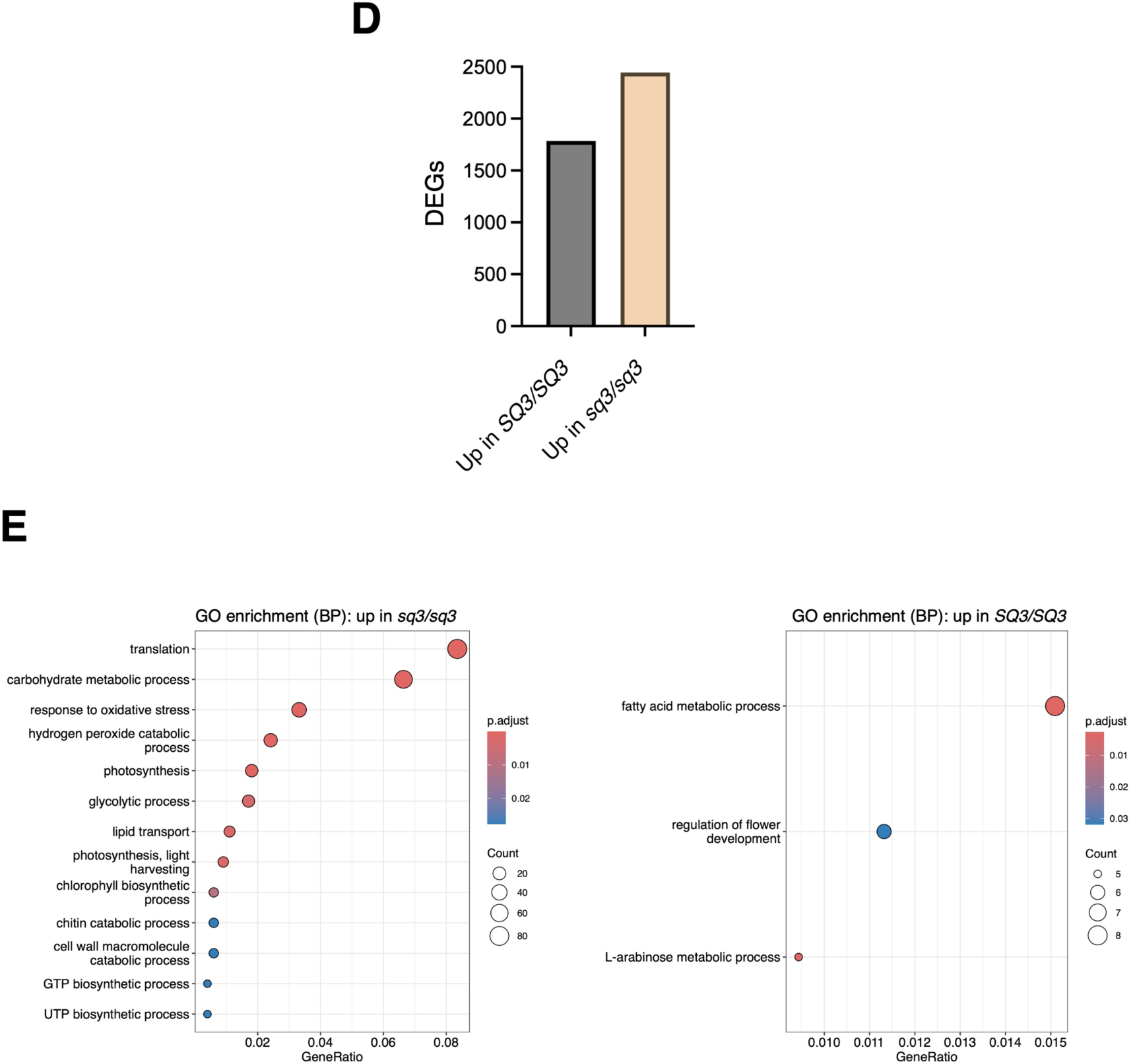
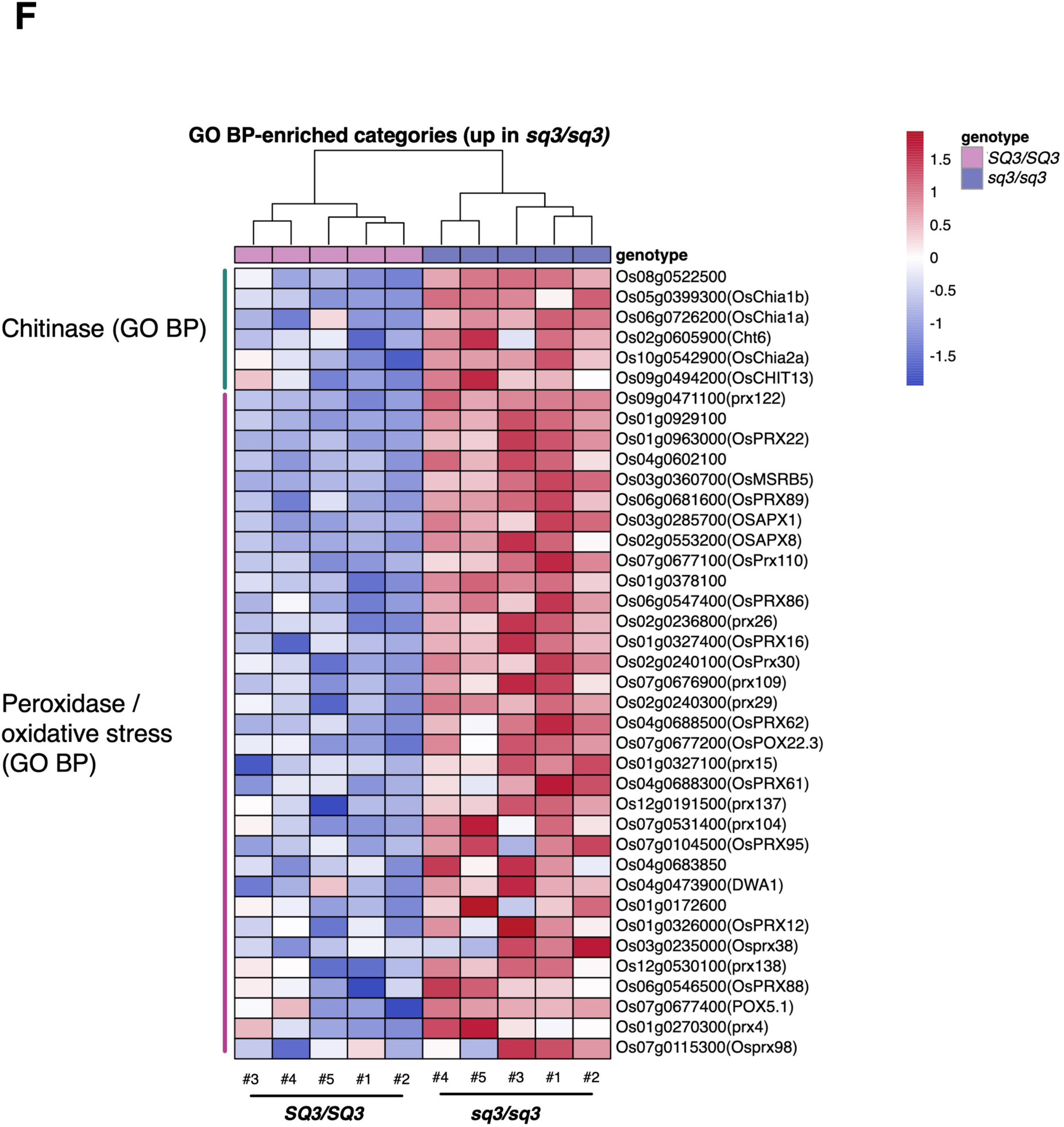
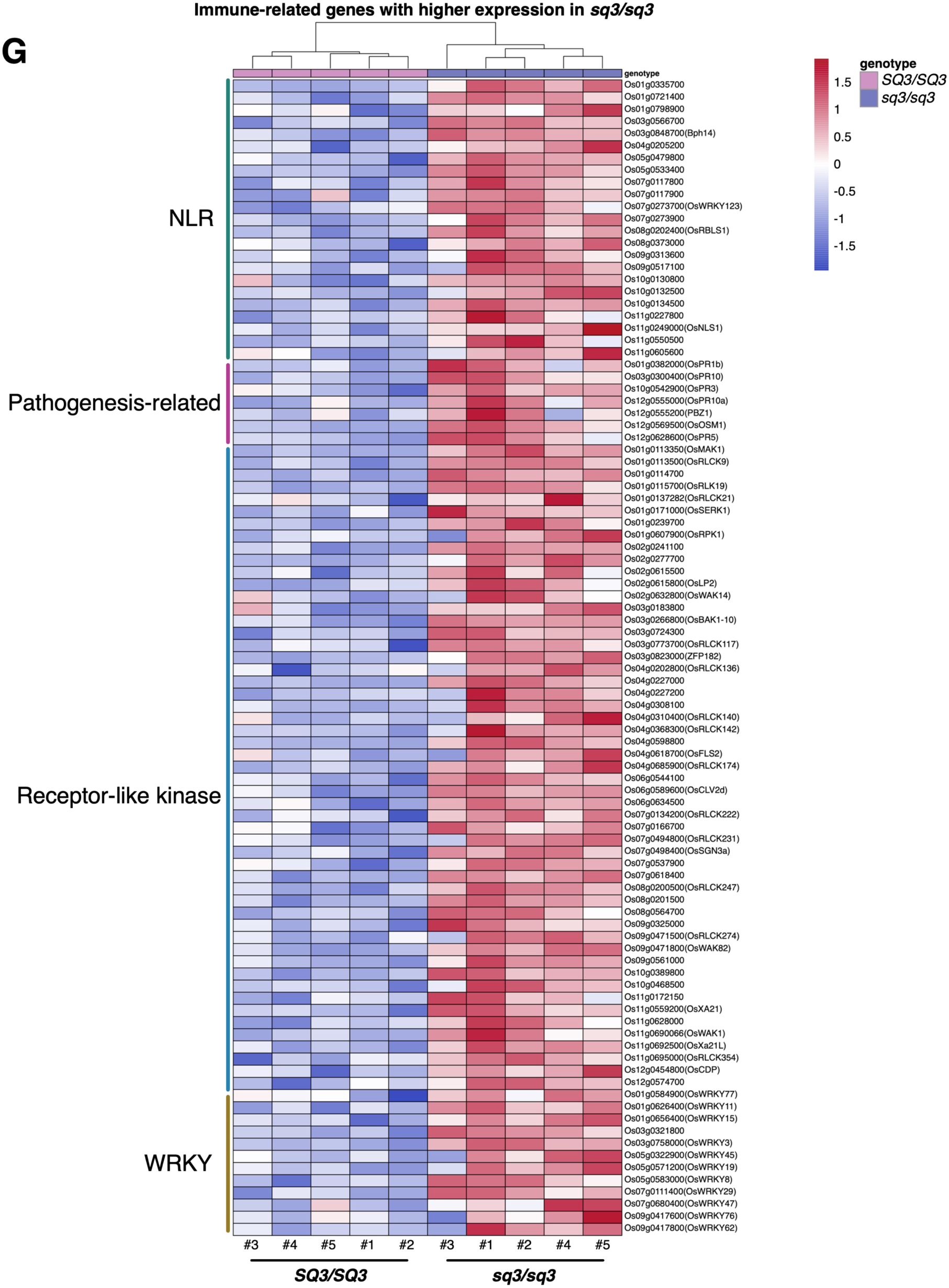
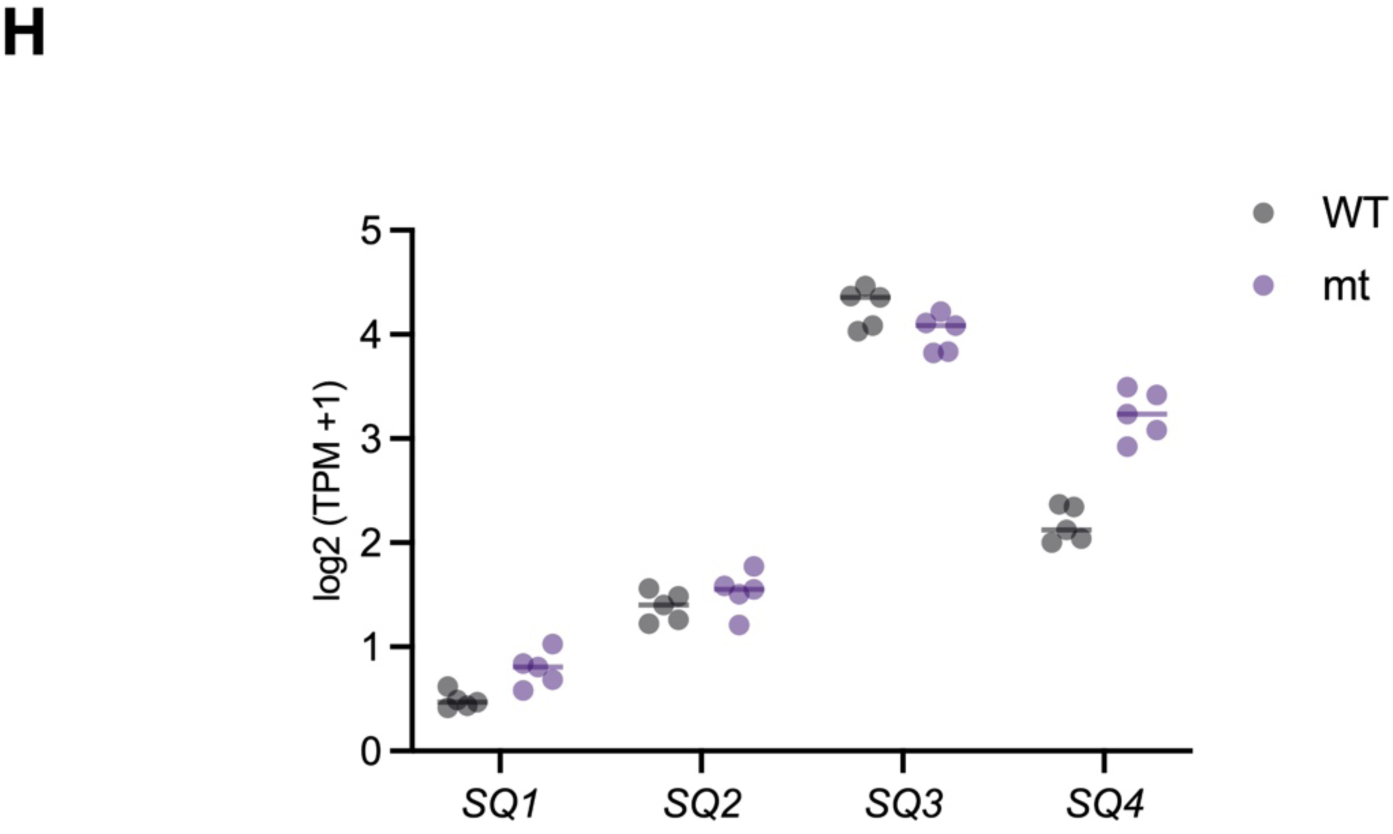
Loss of *SQ3* activates defense-related transcription before visible growth defects appear. **(A)** Experimental design for transcriptome analysis. Segregating T_2_ progeny derived from an *SQ3/sq3* heterozygous plant were grown on Murashige and Skoog (MS) medium under sterile conditions. After 1 week, seedlings were genotyped, and five *SQ3/SQ3* and five *sq3/sq3* seedlings were selected for RNA extraction and RNA sequencing before visible growth defects appeared. **(B)** Principal component analysis of RNA-seq samples based on normalized gene-expression profiles. *SQ3/SQ3* and *sq3/sq3* samples separated primarily along PC1, which explained 77.4% of the total variance. **(C)** Volcano plot of differential gene expression in *sq3/sq3* relative to *SQ3/SQ3* seedlings. Red and cyan dots indicate genes significantly upregulated in *SQ3/SQ3* and *sq3/sq3* seedlings, respectively; gray dots indicate genes that were not significantly differentially expressed. Differentially expressed genes were defined by |log2 fold change| ≥ 1 and FDR < 0.05 (dashed lines). **(D)** Numbers of genes significantly upregulated in *SQ3/SQ3* and *sq3/sq3* seedlings. **(E)** Gene Ontology (Biological Process) enrichment analysis of genes significantly upregulated in *sq3/sq3* (left) and *SQ3/SQ3* (right) seedlings. Dot size indicates the number of genes annotated to each term (Count); color indicates the BH-adjusted *P* value. Only GO BP terms with an adjusted *P* value < 0.05 are shown. **(F)** Heat map of differentially expressed genes belonging to significantly enriched defense-related GO BP terms: chitin catabolic process (Chitinase) and response to oxidative stress/hydrogen peroxide catabolic process (Peroxidase/oxidative stress). Gene membership reflects the enrichment result shown in **(E)**. Color represents row-scaled (z-score) VST-normalized expression; each column represents one biological replicate. **(G)** Heat map of immune-related genes showing higher expression in *sq3/sq3* than in *SQ3/SQ3*, irrespective of statistical significance: NLR receptors, pathogenesis-related proteins, receptor-like kinases, and WRKY transcription factors. Genes were identified by keyword search of RAP-DB InterPro and Description. Color represents row-scaled (z-score) VST-normalized expression; each column represents one biological replicate. **(H)** Expression levels of *SQ1*, *SQ2*, *SQ3*, and *SQ4* in *SQ3/SQ3* and *sq3/sq3* seedlings. Expression values are shown as log_2_(TPM + 1). Each dot represents an individual biological replicate, and horizontal lines indicate the mean.

**Fig. S15.**
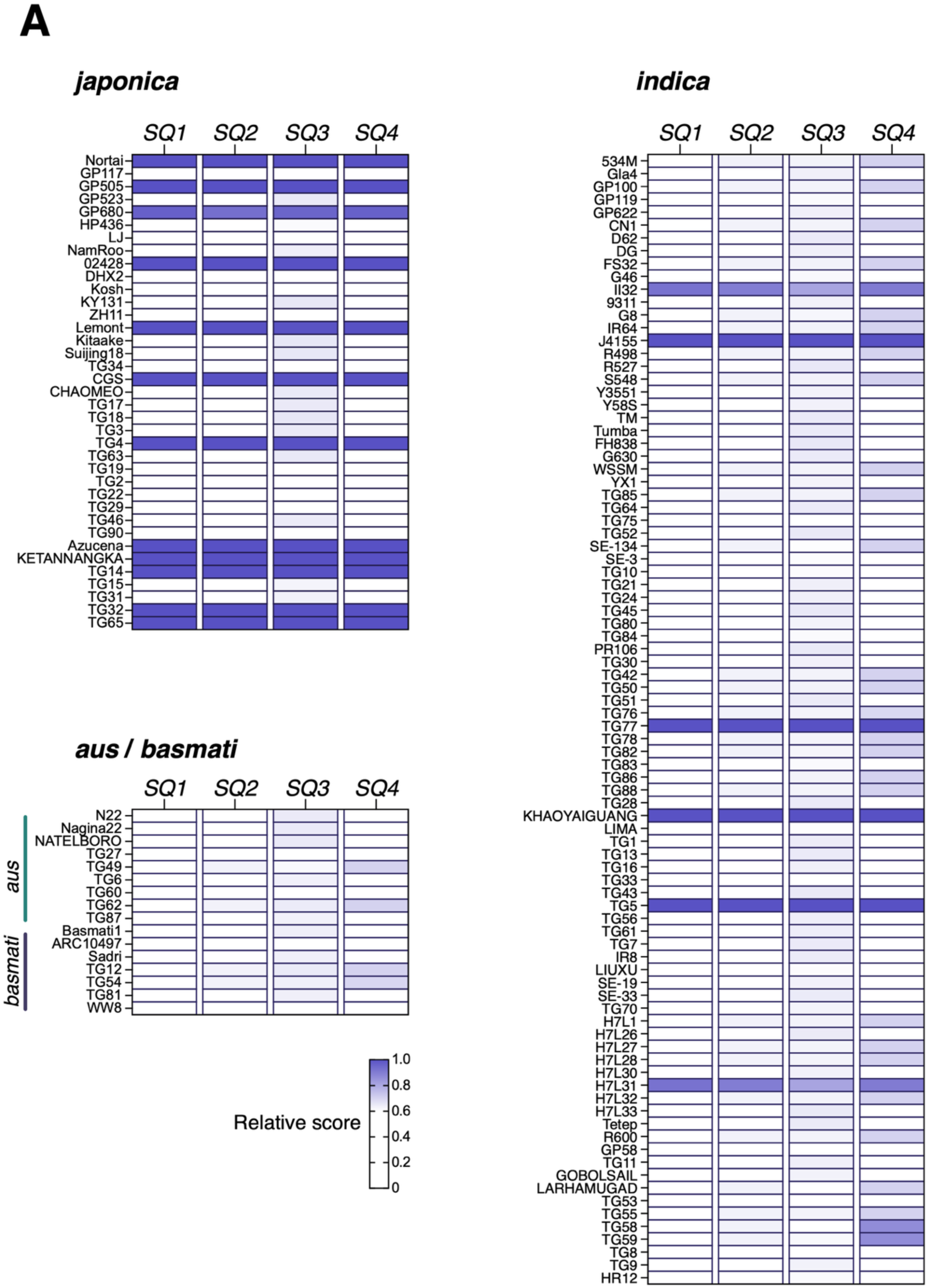
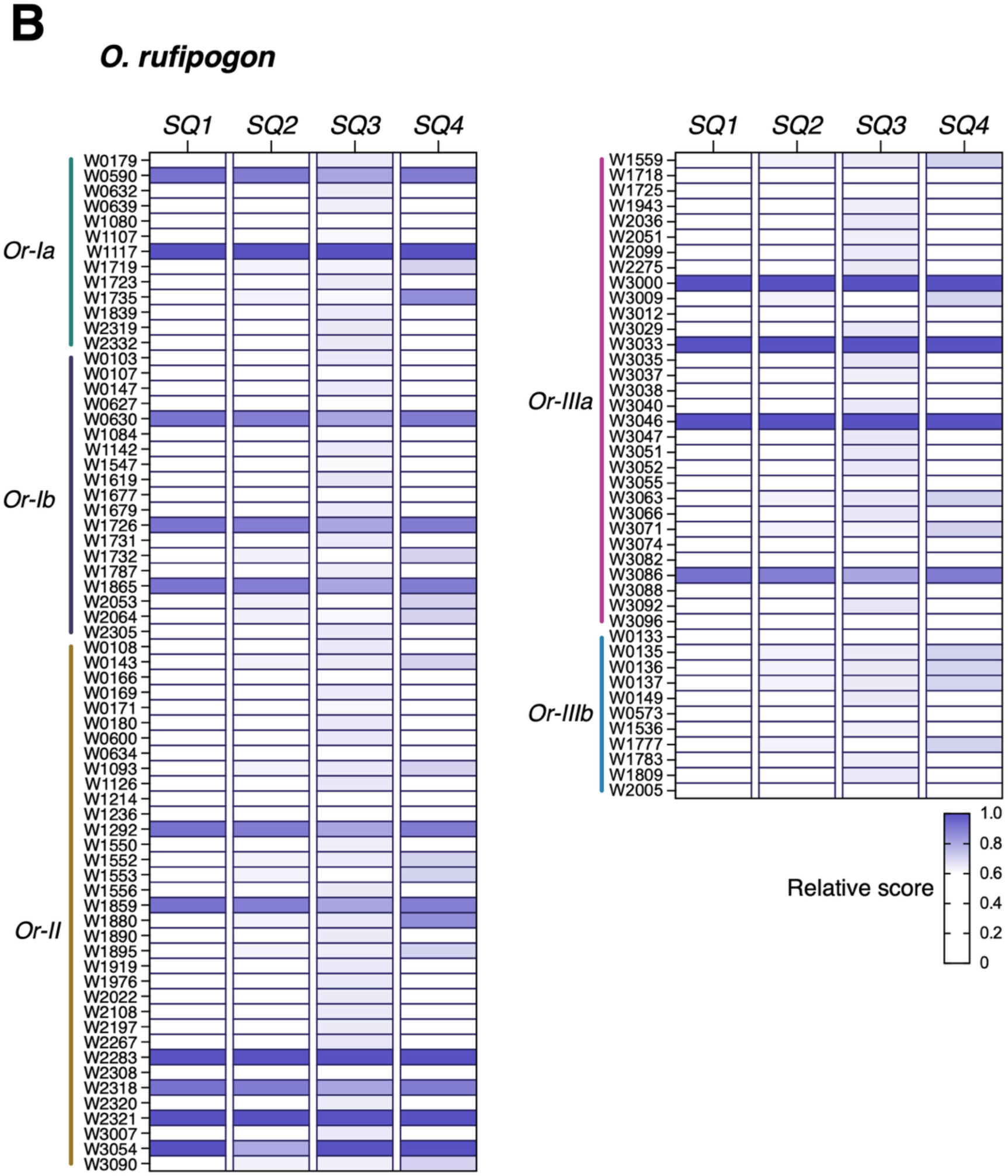
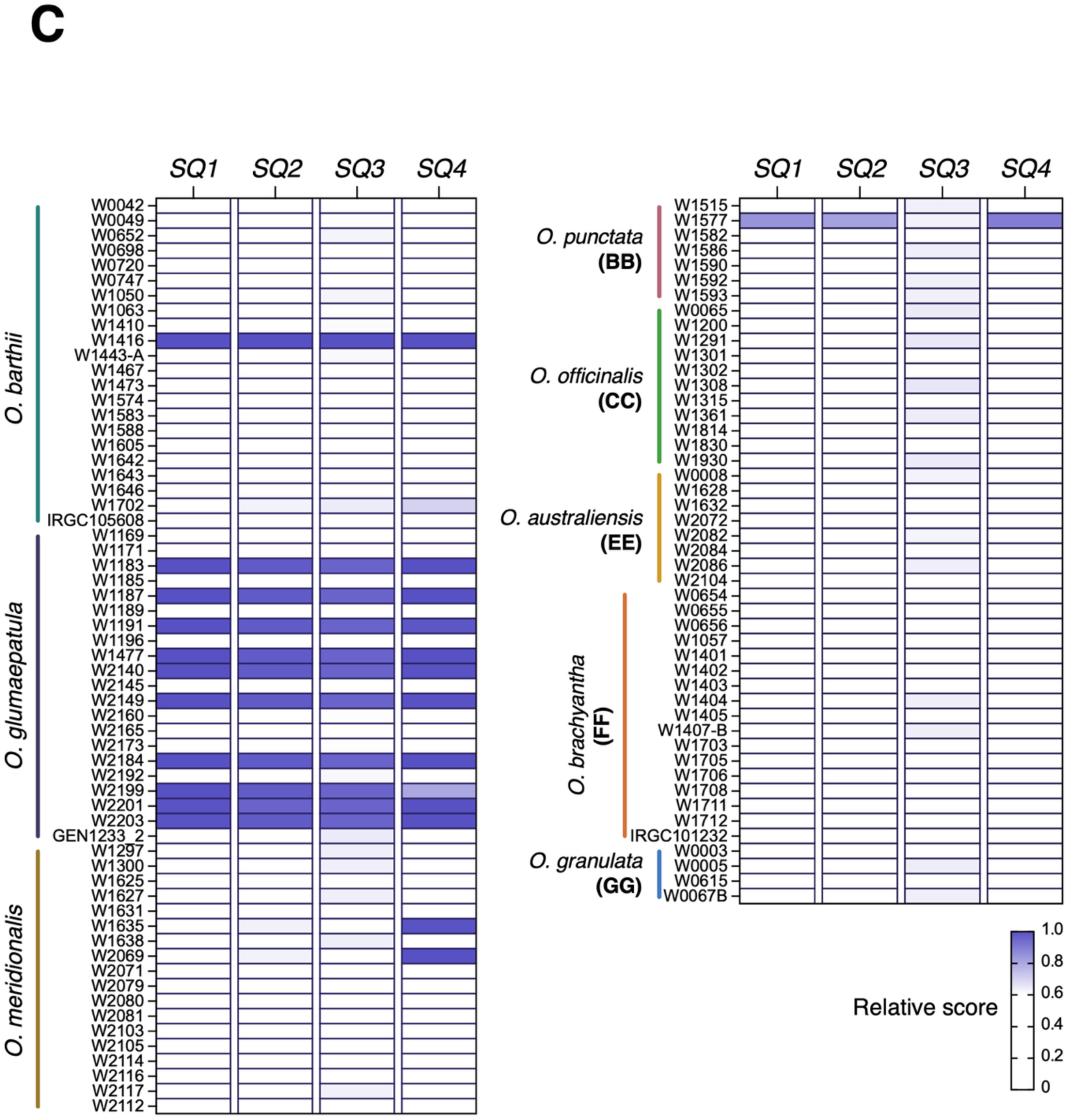
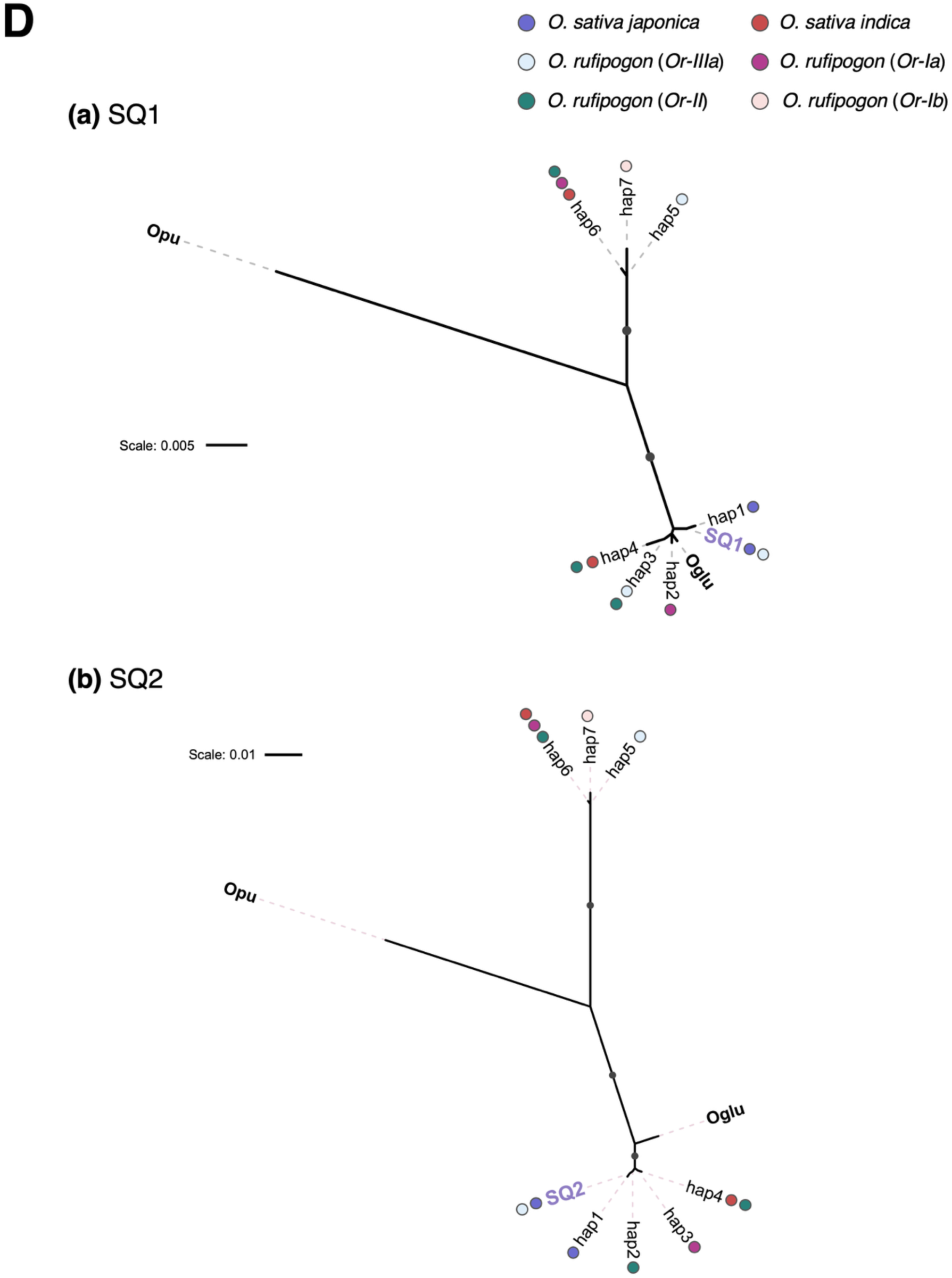
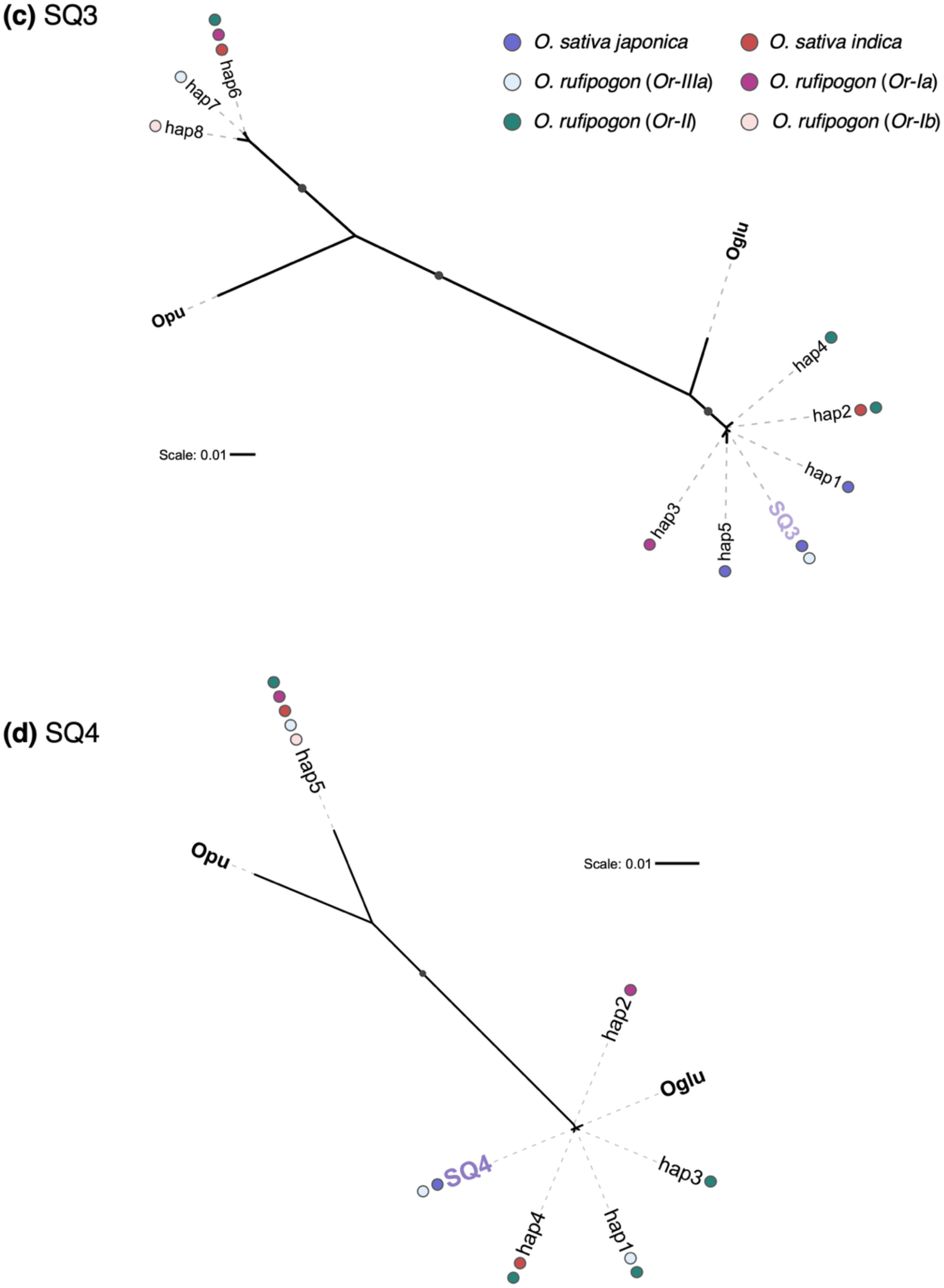
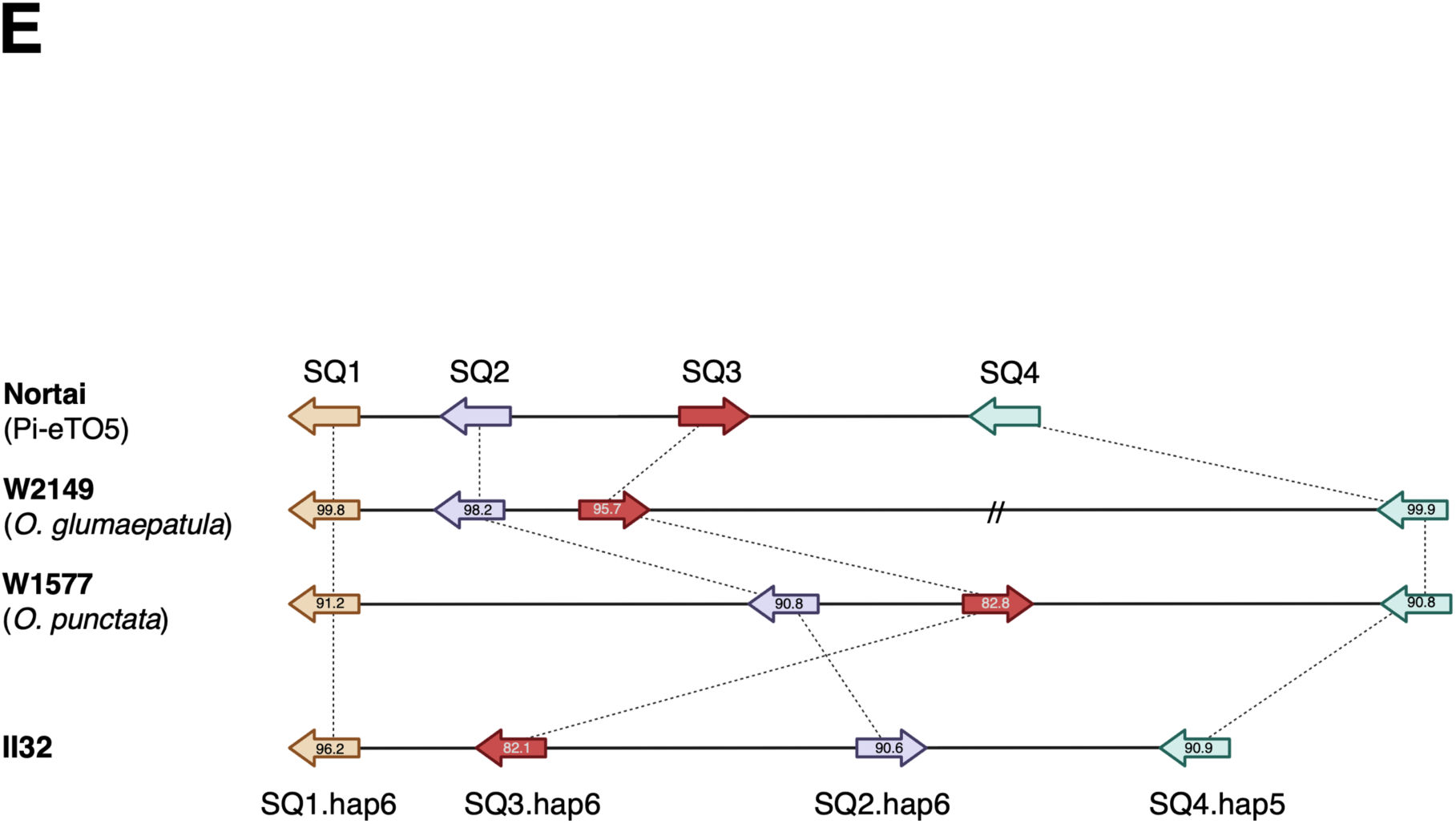
Conservation of *SQ1–SQ4* across cultivated and wild *Oryza* genomes. **(A)** Relative Exonerate score (normalized to Nortai) for SQ1–SQ4 in *O. sativa* accessions from the *japonica*, *aus*, *basmati*, and *indica* groups. **(B)** Relative Exonerate score for SQ1–SQ4 in *O. rufipogon* accessions from previously defined genetic groups *Or-Ia*, *Or-Ib*, *Or-II*, *Or-IIIa*, and *Or-IIIb* (44). **(C)** Relative Exonerate score for SQ1–SQ4 in other wild *Oryza* species (*O. barthii*, *O. glumaepatula*, *O. meridionalis*, *O. punctata* [BB], *O. officinalis* [CC], *O. australiensis* [EE], *O. brachyantha* [FF], *O. granulata* [GG]). In (A)–(C), darker colors indicate higher relative similarity to the corresponding Nortai protein sequence. Accessions with a relative score ≥ 0.6 for all four genes were conservatively classified as carrying a putative complete SQ1–SQ4 haplotype and are indicated in blue. Among the remaining accessions, haplotypes are numbered in order of decreasing frequency, with ties broken by increasing amino acid divergence from Nortai. **(D)** Maximum-likelihood phylogenetic trees of (a) SQ1, (b) SQ2, (c) SQ3, and (d) SQ4 amino acid sequences across the accessions in (A)–(C). Sequences were aligned with MAFFT v7 (--auto) and the tree was inferred with IQ-TREE v2 using ModelFinder (-m MFP), with branch support assessed by 1,000 ultrafast bootstrap replicates and 1,000 SH-aLRT replicates. (**E**) Gene organization of the SQ1–SQ4 region and amino acid identity (%) of each SQ protein relative to Nortai in II32 (*O. sativa indica*), W2149 (*O. glumaepatula*), and W1577 (*O. punctata*). Arrows indicate gene orientation. Gene positions in W2149 and W1577 were confirmed by Nanopore sequencing and *de novo* assembly (see Methods).

**Fig. S16.**
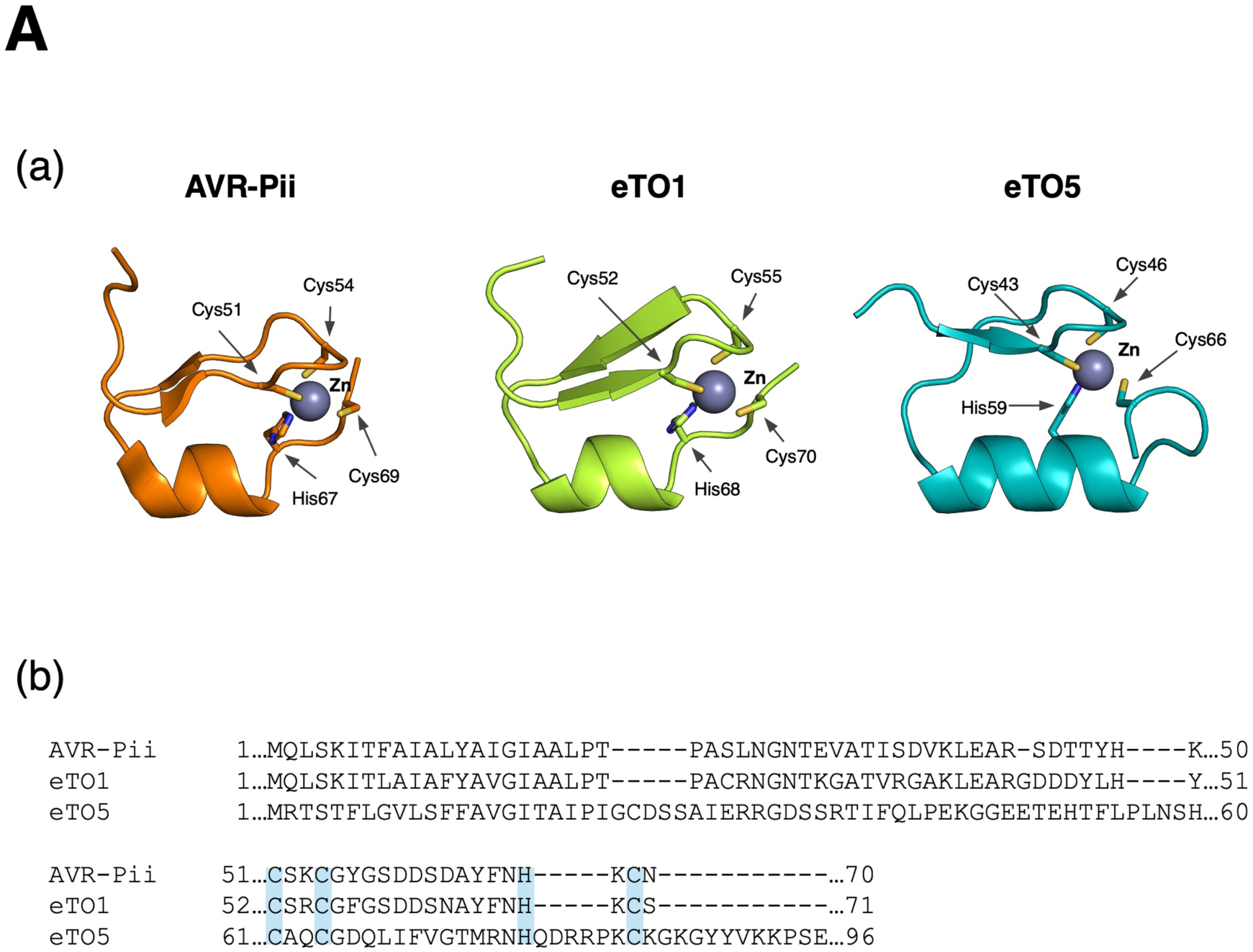
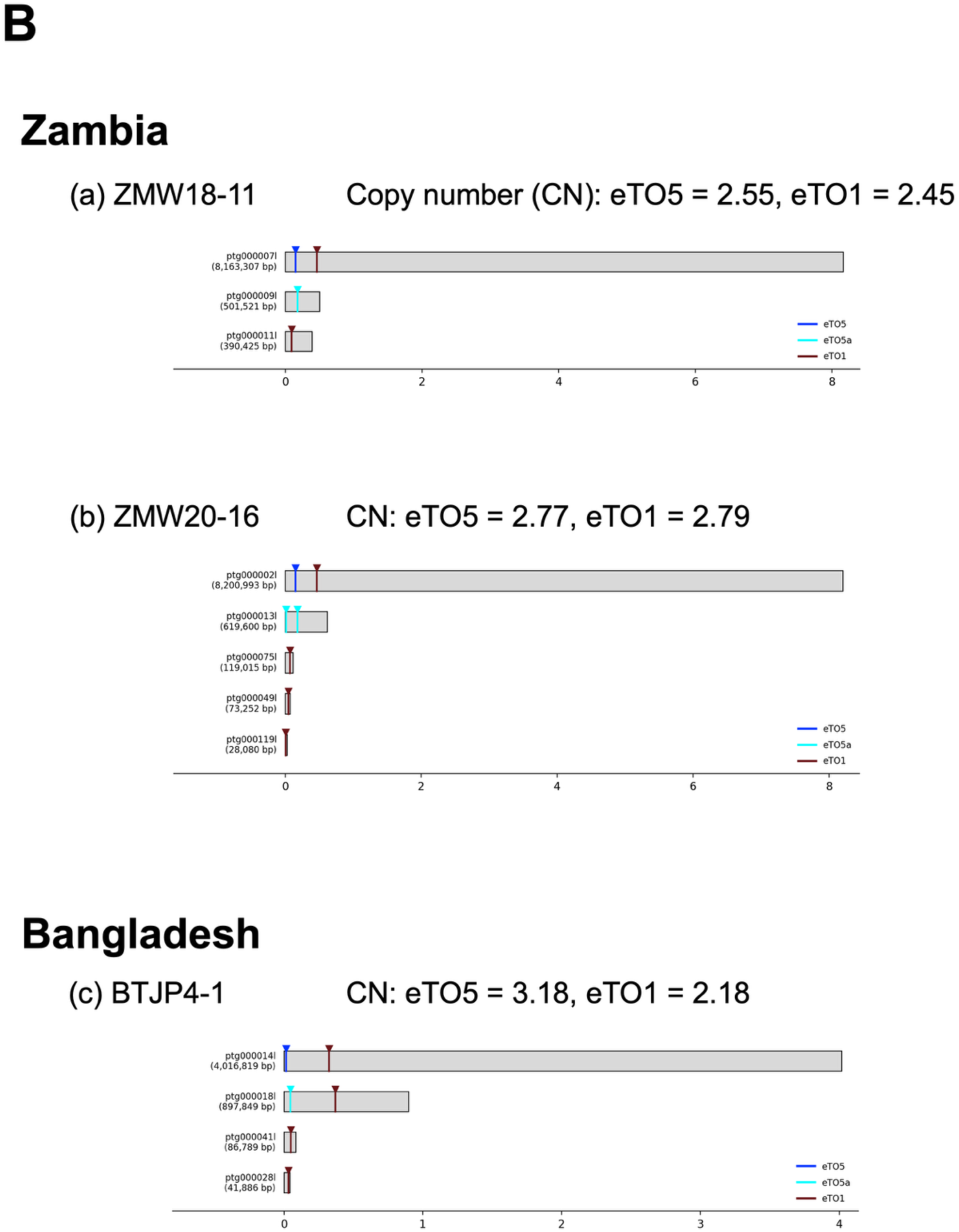
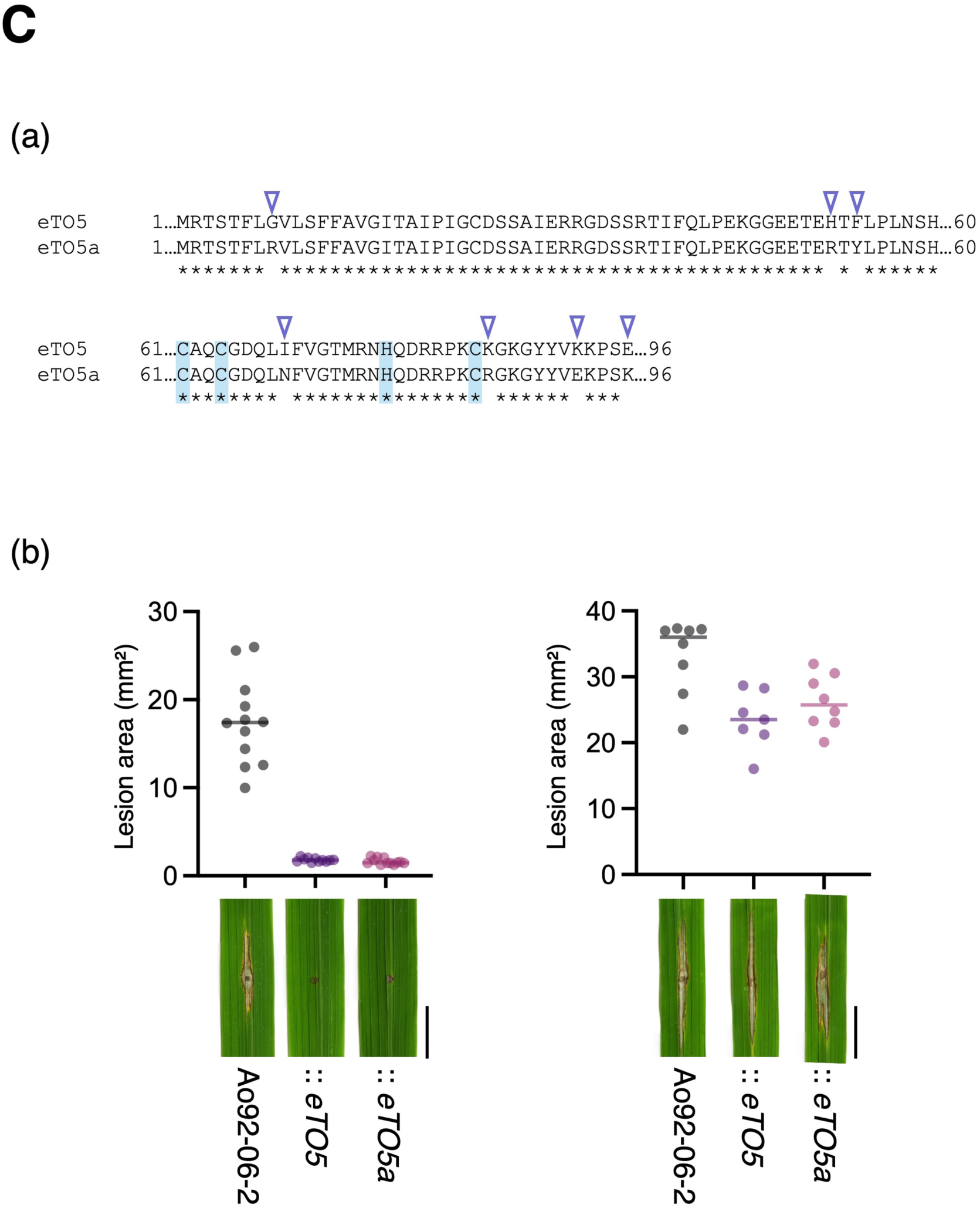
Structural conservation and genomic copy-number variation of ZiF-family effectors. **(A)** Structural and sequence conservation among the ZiF-family effectors AVR-Pii, eTO1, and eTO5. Ribbon representations of AVR-Pii (orange; PDB ID: 7PP2) and AlphaFold 3-predicted structures of eTO1 (green) and eTO5 (cyan) are shown. Zinc ions are depicted as gray spheres, and the side chains of cysteine and histidine residues coordinating the zinc ion are shown as sticks. Amino acid sequence alignment of AVR-Pii, eTO1, and eTO5 highlights conservation of residues forming the zinc-binding motif. **(B)** Assembly-based evidence for multiple eTO5/eTO1-family copies in representative isolates from Zambia and Bangladesh. Positions of *eTO5*, *eTO5a*, and *eTO1* sequence hits are shown on assembled contigs of the Zambian isolates ZMW20-16 and ZMW18-11 and the Bangladeshi isolate BTJP4-1. Contig lengths are indicated in parentheses. Corresponding read-depth-based copy-number (CN) estimates for *eTO5* and *eTO1* are shown for each isolate. **(C)** Sequence and functional comparison of eTO5 and eTO5a. Amino acid sequence alignment of eTO5 and eTO5a shows the sequence differences between the two proteins, with identical residues indicated by asterisks. Representative leaves and lesion-area measurements are shown following inoculation with Ao92-06-2, Ao92-06-2::*eTO5*, and Ao92-06-2::*eTO5a*.

**Fig. S17.**
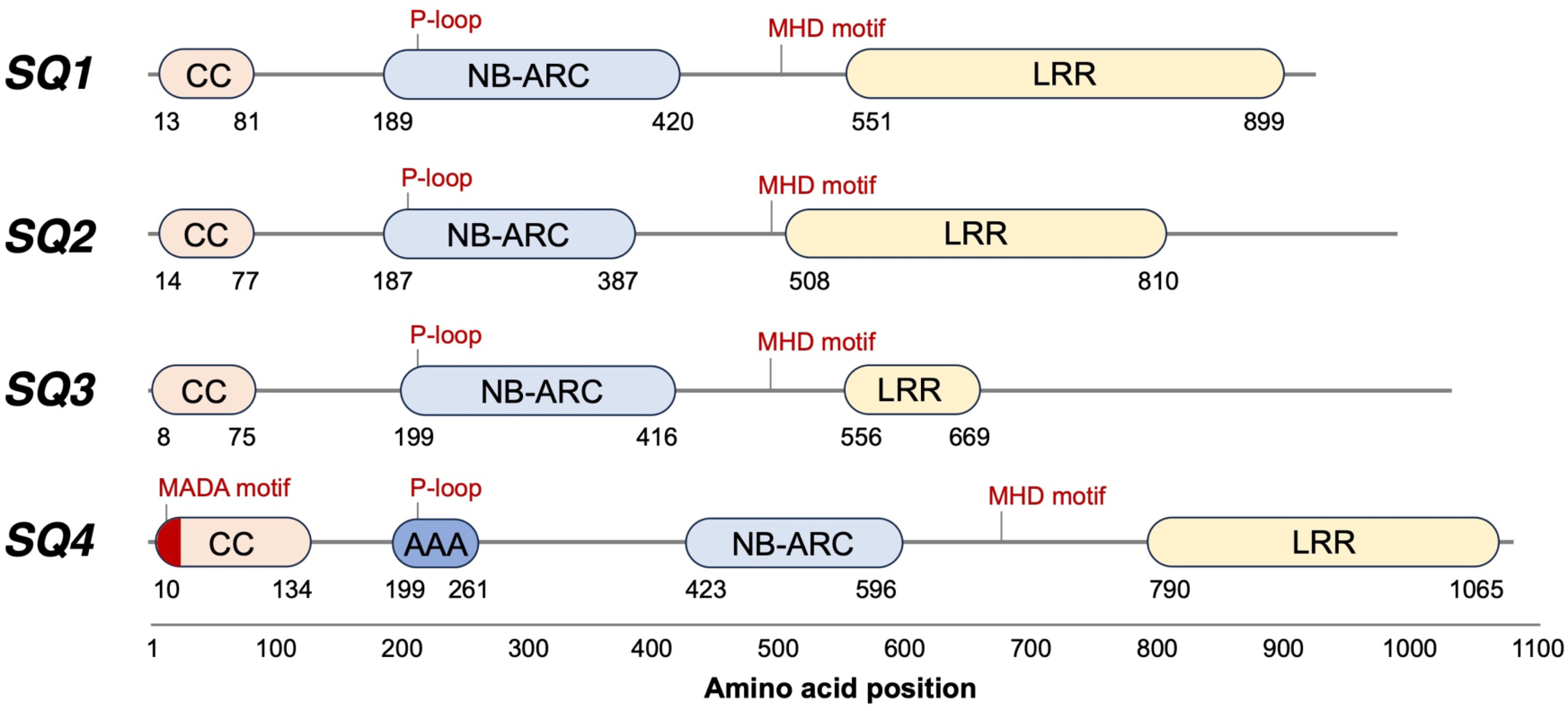
Domain architecture of Pi-eTO5 NLR proteins. Schematic representation of the predicted domain structures of SQ1, SQ2, SQ3, and SQ4, indicating conserved motifs associated with NLR function. All four proteins contain the P-loop and MHD motifs characteristic of NLR proteins. SQ4 uniquely contains a MADA motif, a conserved N-terminal motif associated with NLR-mediated cell-death activity. The distinct distribution of NLR-associated motifs among SQ1–SQ4 is consistent with functional specialization within the Pi-eTO5 NLR quartet.

**Fig. S18.**
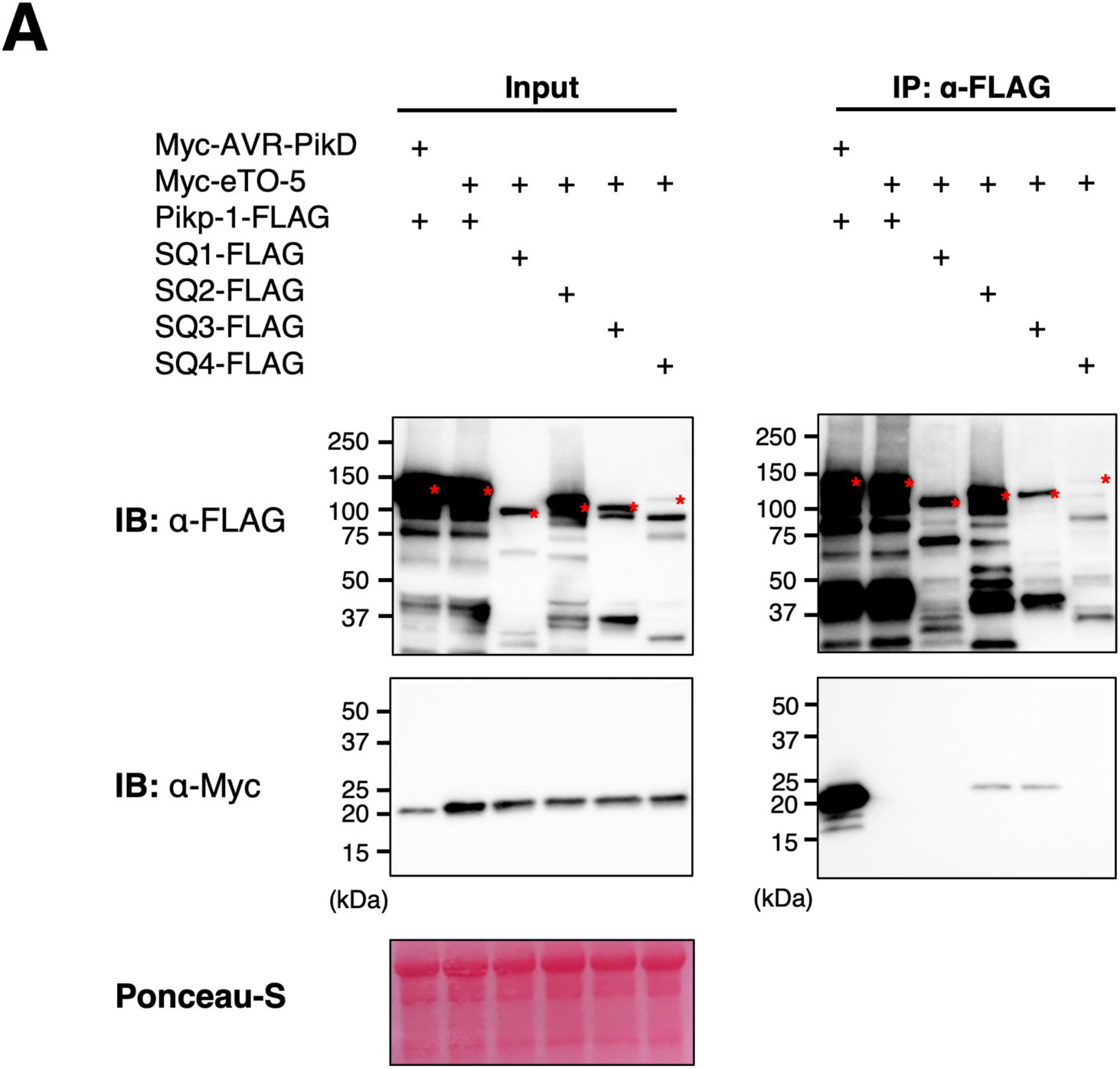
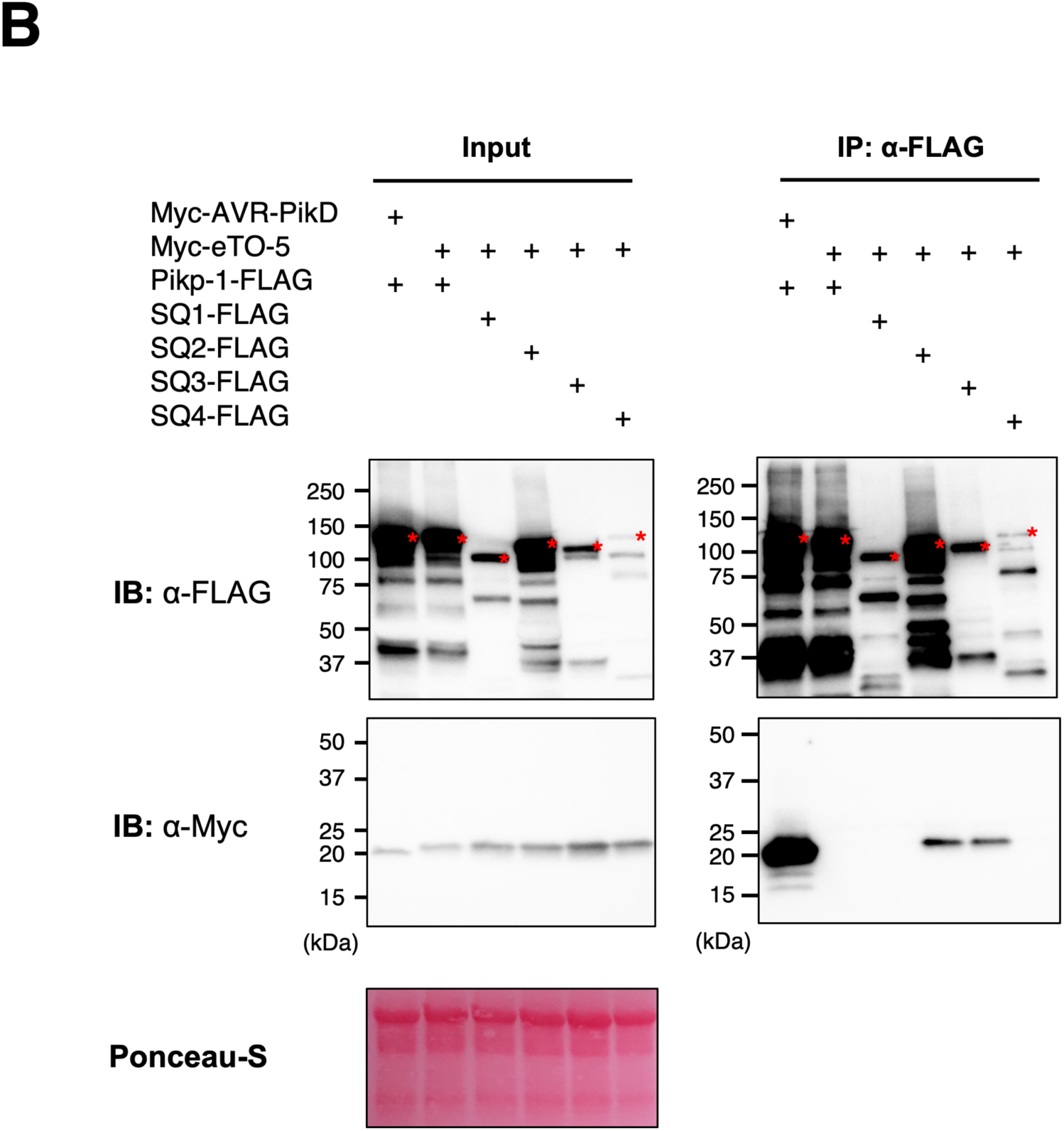
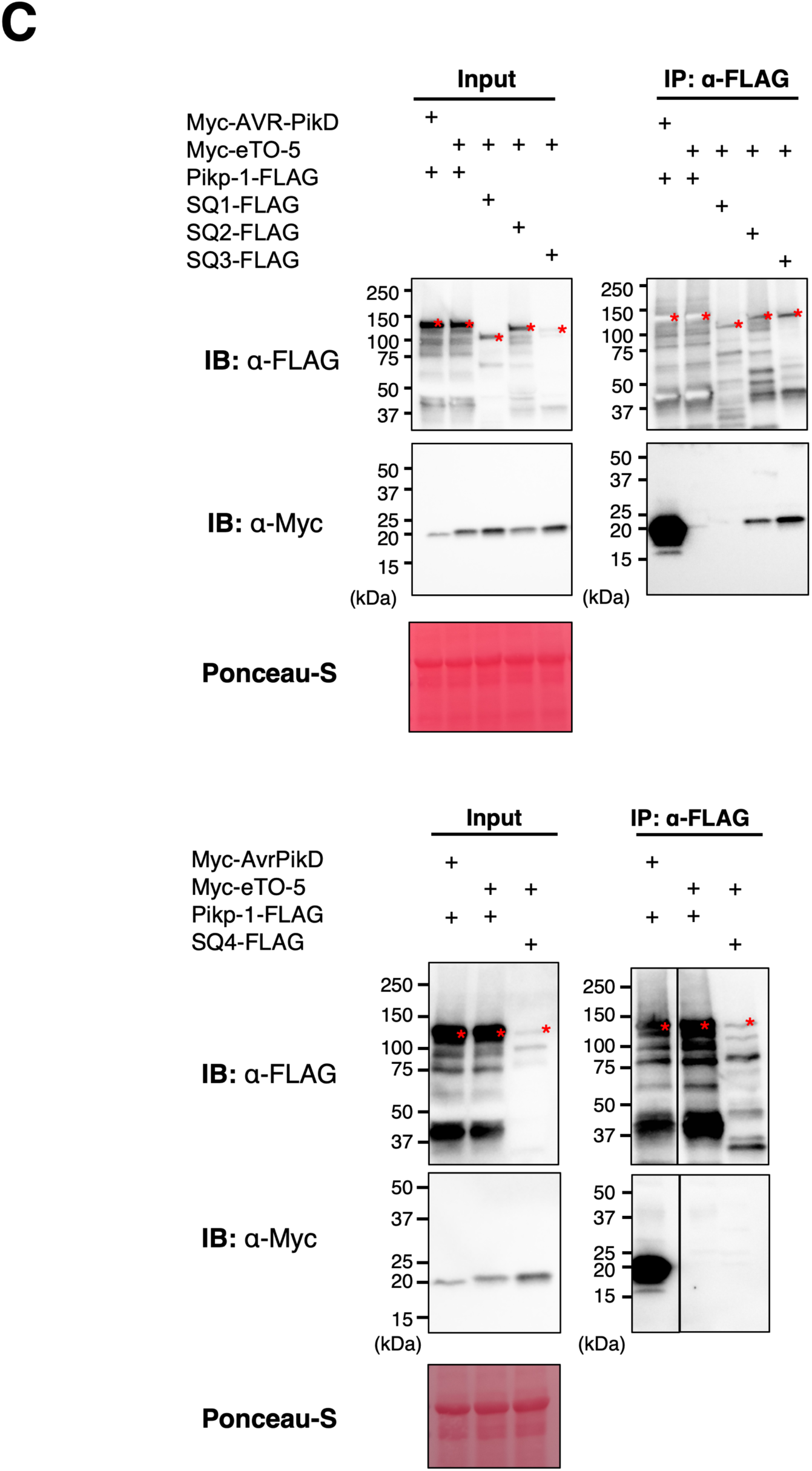
Reproducibility of co-immunoprecipitation assays between eTO5 and Pi-eTO5-linked NLR proteins. Co-immunoprecipitation assays were performed in *Nicotiana benthamiana* transiently expressing 3×FLAG-tagged SQ1, SQ2, SQ3, or SQ4 together with 4×Myc-tagged eTO5. Pikp-1–3×FLAG and AVR-PikD–4×Myc were included as controls. FLAG-tagged NLR proteins were immunoprecipitated with anti-FLAG agarose, and input and immunoprecipitated fractions were analyzed by immunoblotting with anti-FLAG and anti-Myc antibodies. Ponceau S staining is shown as a protein-loading and transfer control. (A–C) Three independent experiments performed using the same experimental conditions. The experiments reproducibly detected association of eTO5 with SQ2 and SQ3, whereas association with SQ1 or SQ4 was not detected under these conditions.

**Data S1.**

Summary statistics for the genome assembly of *M. oryzae* pathotype *Triticum* isolate Br48.

**Data S2.**

One MT isolate and nine MO isolates were used for comparative genomic analyses.

**Data S3.**

A list of 35 MT-specific candidate effector genes identified in *M. oryzae* pathotype *Triticum* isolate Br48.

**Data S4.**

Rice cultivars used for functional screening of MT-specific effector genes and genome sequence analysis.

**Data S5.**

Summary statistics for the Nanopore-based *de novo* genome assemblies of rice cultivars Nortai, WRC43, and Sesia.

**Data S6.**

Genes within the *Pi-eTO5* candidate interval identified by transcriptome-guided annotation of the Nortai genome.

**Data S7.**

Genes differentially expressed between *sq3/sq3* and *SQ3/SQ3* seedlings.

**Data S8.**

Gene Ontology Biological Process (GO BP) over-representation analysis of genes differentially expressed between *sq3/sq3* and *SQ3/SQ3* seedlings.

**Data S9.**

NB-ARC domain sequences used for phylogenetic analysis of rice NLR proteins (Fig. 3).

**Data S10.**

Conservation of SQ1-SQ4 across cultivated and wild *Oryza* genomes, assessed by Exonerate protein2genome alignment.

**Data S11.**

Exonerate alignment scores and haplotype classification of SQ1–SQ4 across *O. sativa* and *O. rufipogon* accessions.

**Data S12.**

Representative amino acid sequences for each SQ1–SQ4 haplotype defined in Data S11.

**Data S13.**

Summary statistics for the Nanopore-based *de novo* genome assemblies of the wild *Oryza* accessions W2149 (*O. glumaepatula*) and W1577 (*O. punctata*).

**Data S14.**

Metadata for 65 *M. oryzae* pathotype *Triticum* (MT) isolates used for SNP-based phylogenetic analysis and copy-number estimation (Fig. 4A).

**Data S15.**

Copy-number estimates of *eTO1, eTO2, eTO5, eTO16, eTO24,* and *eTO31* across 65 *M. oryzae* pathotype *Triticum* (MT) isolates.

**Data S16.**

Per-isolate BUSCO depth statistics used for copy-number normalization and single-copy classification thresholds.

**Data S17.**

Primer sequences in this study used.

## REFERENCES

1. J. Kourelis, R. A. L. van der Hoorn, Defended to the nines: 25 years of resistance gene cloning identifies nine mechanisms for *R* protein function. Plant Cell 30, 285–299 (2018).

2. S. Lee, V. M. Whitaker, S. F. Hutton, Potential applications of non-host resistance for crop improvement. Front. Plant Sci. 7, 997 (2016).

3. B. C. Couch, I. Fudal, M.-H. Lebrun, D. Tharreau, B. Valent, P. Van Kim, J.-L. Nottéghem, L. M. Kohn, Origins of host-specific populations of the blast pathogen *Magnaporthe oryzae* in crop domestication with subsequent expansion of pandemic clones on rice and weeds of rice. Genetics 170, 613–630 (2005).

4. P. Gladieux, B. Condon, S. Ravel, D. Soanes, J. L. N. Maciel, A. Nhani Jr., L. Chen, R. Terauchi, M.-H. Lebrun, D. Tharreau, T. Mitchell, K. F. Pedley, B. Valent, N. J. Talbot, M. Farman, E. Fournier, Gene flow between divergent cereal- and grass-specific lineages of the rice blast fungus *Magnaporthe oryzae*. mBio 9, e01219–17 (2018).

5. S. Igarashi, C. M. Utiamada, L. C. Igarashi, A. H. Kazuma, R. S. Lopes, Pyricularia em trigo. 1. Ocorrência de Pyricularia sp. no estado do Paraná. Fitopatol. Bras. 11, 351–352 (1986).

6. M. T. Islam, D. Croll, P. Gladieux, D. M. Soanes, A. Persoons, P. Bhattacharjee, M. S. Hossain, D. R. Gupta, M. M. Rahman, M. G. Mahboob, N. Cook, M. U. Salam, M. Z. Surovy, V. B. Sancho, J. L. N. Maciel, A. Nhani Júnior, V. L. Castroagudín, J. T. de Assis Reges, P. C. Ceresini, S. Ravel, R. Kellner, E. Fournier, D. Tharreau, M.-H. Lebrun, B. A. McDonald, T. Stitt, D. Swan, N. J. Talbot, D. G. O. Saunders, J. Win, S. Kamoun, Emergence of wheat blast in Bangladesh was caused by a South American lineage of *Magnaporthe oryzae*. BMC Biol. 14, 84 (2016).

7. P. K. Malaker, N. C. D. Barma, T. P. Tiwari, W. J. Collis, E. Duveiller, P. K. Singh, A. K. Joshi, R. P. Singh, H.-J. Braun, G. L. Peterson, K. F. Pedley, M. L. Farman, B. Valent, First report of wheat blast caused by *Magnaporthe oryzae* pathotype *triticum* in Bangladesh. Plant Dis. 100, 2330 (2016).

8. B. Tembo, R. M. Mulenga, S. Sichilima, K. K. M’siska, M. Mwale, P. C. Chikoti, P. K. Singh, X. He, K. F. Pedley, G. L. Peterson, R. P. Singh, H. J. Braun, Detection and characterization of fungus (*Magnaporthe oryzae* pathotype *Triticum*) causing wheat blast disease on rain-fed grown wheat (*Triticum aestivum* L.) in Zambia. PLOS ONE 15, e0238724 (2020).

9. S. M. Latorre, V. M. Were, A. J. Foster, T. Langner, A. Malmgren, A. Harant, S. Asuke, S. Reyes-Avila, D. R. Gupta, C. Jensen, W. Ma, N. U. Mahmud, M. S. Mehebub, R. M. Mulenga, A. N. M. Muzahid, S. K. Paul, S. M. F. Rabby, A. A. M. Rahat, L. Ryder, R.-K. Shrestha, S. Sichilima, D. M. Soanes, P. K. Singh, A. R. Bentley, D. G. O. Saunders, Y. Tosa, D. Croll, K. H. Lamour, T. Islam, B. Tembo, J. Win, N. J. Talbot, H. A. Burbano, S. Kamoun, Genomic surveillance uncovers a pandemic clonal lineage of the wheat blast fungus. PLOS Biol. 21, e3002052 (2023).

10. C. D. Cruz, G. L. Peterson, W. W. Bockus, P. Kankanala, J. Dubcovsky, K. W. Jordan, E. Akhunov, F. Chumley, F. D. Baldelomar, B. Valent, The 2NS translocation from *Aegilops ventricosa* confers resistance to the *Triticum* pathotype of *Magnaporthe oryzae*. Crop Sci. 56, 990–1000 (2016).

11. V. L. Anh, N. T. Anh, A. G. Tagle, T. T. P. Vy, Y. Inoue, S. Takumi, I. Chuma, Y. Tosa, *Rmg8*, a new gene for resistance to *Triticum* isolates of *Pyricularia oryzae* in hexaploid wheat. Phytopathology 105, 1568–1572 (2015).

12. Y. Inoue, T. T. P. Vy, K. Yoshida, H. Asano, C. Mitsuoka, S. Asuke, V. L. Anh, C. J. R. Cumagun, I. Chuma, R. Terauchi, K. Kato, T. Mitchell, B. Valent, M. Farman, Y. Tosa, Evolution of the wheat blast fungus through functional losses in a host specificity determinant. Science 357, 80–83 (2017).

13. J. Sperschneider, P. N. Dodds, EffectorP 3.0: Prediction of apoplastic and cytoplasmic effectors in fungi and oomycetes. Mol. Plant-Microbe Interact. 35, 146–156 (2022).

14. K. Yoshida, H. Saitoh, S. Fujisawa, H. Kanzaki, H. Matsumura, K. Yoshida, Y. Tosa, I. Chuma, Y. Takano, J. Win, S. Kamoun, R. Terauchi, Association genetics reveals three novel avirulence genes from the rice blast fungal pathogen *Magnaporthe oryzae*. Plant Cell 21, 1573–1591 (2009).

15. H. Takagi, A. Uemura, H. Yaegashi, M. Tamiru, A. Abe, C. Mitsuoka, H. Utsushi, S. Natsume, H. Kanzaki, H. Matsumura, H. Saitoh, K. Yoshida, L. M. Cano, S. Kamoun, R. Terauchi, MutMap-Gap: whole-genome resequencing of mutant F2 progeny bulk combined with de novo assembly of gap regions identifies the rice blast resistance gene *Pii*. New Phytol. 200, 276–283 (2013).

16. Y. Okuyama, H. Kanzaki, A. Abe, K. Yoshida, M. Tamiru, H. Saitoh, T. Fujibe, H. Matsumura, M. Shenton, D. C. Galam, J. Undan, A. Ito, T. Sone, R. Terauchi, A multifaceted genomics approach allows the isolation of the rice *Pia*-blast resistance gene consisting of two adjacent NBS-LRR protein genes. Plant J. 66, 467–479 (2011).

17. H. Takagi, A. Abe, K. Yoshida, S. Kosugi, S. Natsume, C. Mitsuoka, A. Uemura, H. Utsushi, M. Tamiru, S. Takuno, H. Innan, L. M. Cano, S. Kamoun, R. Terauchi, QTL-seq: rapid mapping of quantitative trait loci in rice by whole genome resequencing of DNA from two bulked populations. Plant J. 74, 174–183 (2013).

18. J. C. De la Concepcion, T. Langner, K. Fujisaki, X. Yan, V. Were, A. H. C. Lam, I. Saado, H. J. Brabham, J. Win, K. Yoshida, N. J. Talbot, R. Terauchi, S. Kamoun, M. J. Banfield, Zinc-finger (ZiF) fold secreted effectors form a functionally diverse family across lineages of the blast fungus Magnaporthe oryzae. PLoS Pathog. 20, e1012277 (2024).

19. H. Adachi, M. P. Contreras, A. Harant, C.-H. Wu, L. Derevnina, T. Sakai, C. Duggan, E. Moratto, T. O. Bozkurt, A. Maqbool, J. Win, S. Kamoun, An N-terminal motif in NLR immune receptors is functionally conserved across distantly related plant species. eLife 8, e49956 (2019).

## References

1. C. A. Schneider, W. S. Rasband, K. W. Eliceiri, NIH Image to ImageJ: 25 years of image analysis. Nat. Methods 9, 671–675 (2012).

2. K. Yoshida et al., Host specialization of the blast fungus *Magnaporthe oryzae* is associated with dynamic gain and loss of genes linked to transposable elements. BMC Genomics 17, 370 (2016).

3. C. C. Lo, P. S. G. Chain, Rapid evaluation and quality control of next generation sequencing data with FaQCs. BMC Bioinformatics 15, 366 (2014).

4. A. V. Zimin et al., The MaSuRCA genome assembler. Bioinformatics 29, 2669–2677 (2013).

5. M. Manni et al., BUSCO update: Novel and streamlined workflows along with broader and deeper phylogenetic coverage for scoring of eukaryotic, prokaryotic, and viral genomes. Mol. Biol. Evol. 38, 4647–4654 (2021).

6. D. Kim, J. M. Paggi, C. Park, C. Bennett, S. L. Salzberg, Graph-based genome alignment and genotyping with HISAT2 and HISAT-genotype. Nat. Biotechnol. 37, 907–915 (2019).

7. M. Pertea et al., StringTie enables improved reconstruction of a transcriptome from RNA-seq reads. Nat. Biotechnol. 33, 290–295 (2015).

8. G. Pertea, M. Pertea, GFF utilities: GffRead and GffCompare. F1000Research 9, 304 (2020).

9. B. J. Haas et al., *De novo* transcript sequence reconstruction from RNA-seq using the Trinity platform for reference generation and analysis. Nat. Protoc. 8, 1494–1512 (2013).

10. L. Fu, B. Niu, Z. Zhu, S. Wu, W. Li, CD-HIT: Accelerated for clustering the next-generation sequencing data. Bioinformatics 28, 3150–3152 (2012).

11. T. N. Petersen, S. Brunak, G. von Heijne, H. Nielsen, SignalP 4.0: Discriminating signal peptides from transmembrane regions. Nat. Methods 8, 785–786 (2011).

12. J. Sperschneider, P. N. Dodds, EffectorP 3.0: Prediction of apoplastic and cytoplasmic effectors in fungi and oomycetes. Mol. Plant-Microbe Interact. 35, 146–156 (2022).

13. B. Langmead, S. L. Salzberg, Fast gapped-read alignment with Bowtie 2. Nat. Methods 9, 357–359 (2012).

14. P. Danecek et al., Twelve years of SAMtools and BCFtools. GigaScience 10, giab008 (2021).

15. B. S. Pedersen, A. R. Quinlan, Mosdepth: Quick coverage calculation for genomes and exomes. Bioinformatics 34, 867–868 (2018).

16. K. Yoshida et al., Association genetics reveals three novel avirulence genes from the rice blast fungal pathogen *Magnaporthe oryzae*. Plant Cell 21, 1573–1591 (2009).

17. J. A. Sweigard, F. Chumley, A. Carroll, L. Farrall, B. Valent, A series of vectors for fungal transformation. Fungal Genetics Reports 44, 52–53 (1997).

18. H. Takagi et al., QTL-seq: Rapid mapping of quantitative trait loci in rice by whole-genome resequencing of DNA from two bulked populations. Plant J. 74, 174–183 (2013).

19. H. Takagi et al., Rice blast resistance gene Pii is controlled by a pair of NBS-LRR genes Pii-1 and Pii-2. bioRxiv, 227132 (2017).

20. Y. Okuyama et al., A multifaceted genomics approach allows the isolation of the rice Pia-blast resistance gene consisting of two adjacent NBS-LRR protein genes. Plant J. 66, 467– 479 (2011).

21. H. Cheng et al., Efficient near-telomere-to-telomere assembly of nanopore simplex reads. Nature 655, 166–173 (2026).

22. M. Alonge et al., Automated assembly scaffolding using RagTag elevates a new tomato system for high-throughput genome editing. Genome Biol. 23, 258 (2022).

23. S. Chen, Y. Zhou, Y. Chen, J. Gu, fastp: An ultra-fast all-in-one FASTQ preprocessor. Bioinformatics 34, i884–i890 (2018).

24. Y. Sugihara et al., High-performance pipeline for MutMap and QTL-seq. PeerJ 10, e13170 (2022).

25. S. Natsume et al., V-primer: Software for the efficient design of genome-wide InDel and SNP markers from multi-sample variant call format (VCF) genotyping data. Breed. Sci. 73, 415– 420 (2023).

26. J. Kourelis, T. Sakai, H. Adachi, S. Kamoun, RefPlantNLR is a comprehensive collection of experimentally validated plant disease resistance proteins from the NLR family. PLoS Biol. 19, e3001124 (2021).

27. N. Tanaka et al., Investigation of the genetic diversity of a rice core collection of Japanese landraces using whole-genome sequencing. Plant Cell Physiol. 61, 2087–2096 (2020a).

28. N. Tanaka et al., Whole-genome sequencing of the NARO World Rice Core Collection (WRC) as the basis for diversity and association studies. Plant Cell Physiol. 61, 922–932 (2020b).

29. M. Mikami, S. Toki, M. Endo, Comparison of CRISPR/Cas9 expression constructs for efficient targeted mutagenesis in rice. Plant Mol. Biol. 88, 561–572 (2015).

30. Y. Naito, K. Hino, H. Bono, K. Ui-Tei, CRISPRdirect: Software for designing CRISPR/Cas guide RNA with reduced off-target sites. Bioinformatics 31, 1120–1123 (2015).

31. Y. Liao, G. K. Smyth, W. Shi, featureCounts: An efficient general purpose program for assigning sequence reads to genomic features. Bioinformatics 30, 923–930 (2014).

32. D. J. McCarthy, Y. Chen, G. K. Smyth, Differential expression analysis of multifactor RNA-Seq experiments with respect to biological variation. Nucleic Acids Res. 40, 4288–4297 (2012).

33. M. I. Love, W. Huber, S. Anders, Moderated estimation of fold change and dispersion for RNA-seq data with DESeq2. Genome Biol. 15, 550 (2014).

34. T. Wu et al., clusterProfiler 4.0: A universal enrichment tool for interpreting omics data. The Innovation 2, 100141 (2021).

35. H. Sakai et al., Rice Annotation Project Database (RAP-DB): An integrative and interactive database for rice genomics. Plant Cell Physiol. 54, e6 (2013).

36. I. Ashikawa et al., Two adjacent nucleotide-binding site–leucine-rich repeat class genes are required to confer Pikm-specific rice blast resistance. Genetics 180, 2267–2276 (2008).

37. K. Hayashi, H. Yoshida, Refunctionalization of the ancient rice blast disease resistance gene Pit by the recruitment of a retrotransposon as a promoter. Plant J. 57, 413–425 (2009).

38. R. C. Edgar, Muscle5: High-accuracy alignment ensembles enable unbiased assessments of sequence homology and phylogeny. Nature Commun. 13, 6968 (2022).

39. D. T. Jones, W. R. Taylor, J. M. Thornton, The rapid generation of mutation data matrices from protein sequences. Comput. Appl. Biosci. 8, 275–282 (1992).

40. A. M. Kozlov, D. Darriba, T. Flouri, B. Morel, A. Stamatakis, RAxML-NG: A fast, scalable and user-friendly tool for maximum likelihood phylogenetic inference. Bioinformatics 35, 4453–4455 (2019).

41. I. Letunic, P. Bork, Interactive Tree of Life (iTOL) v6: Recent updates to the phylogenetic tree display and annotation tool. Nucleic Acids Res. 52, W78–W82 (2024).

42. P. Qin et al., Pan-genome analysis of 33 genetically diverse rice accessions reveals hidden genomic variations. Cell 184, 3542–3558.e16 (2021).

43. F. Zhang et al., Long-read sequencing of 111 rice genomes reveals significantly larger pan-genomes. Genome Res. 32, 853–863 (2022).

44. D. Guo et al., A pangenome reference of wild and cultivated rice. Nature 642, 662–671 (2025).

45. G. S. C. Slater, E. Birney, Automated generation of heuristics for biological sequence comparison. BMC Bioinformatics 6, 31 (2005).

65. M. Shimizu, et al., A genetically linked pair of NLR immune receptors shows contrasting patterns of evolution. Proc. Natl. Acad. Sci. U.S.A. 119, e2116896119 (2022).

47. G. Holley et al., Ratatosk: Hybrid error correction of long reads enables accurate variant calling and assembly. Genome Biol. 22, 28 (2021).

48. K. Katoh, D. M. Standley, MAFFT multiple sequence alignment software version 7: Improvements in performance and usability. Mol. Biol. Evol. 30, 772–780 (2013).

49. L.-T. Nguyen, H. A. Schmidt, A. von Haeseler, B. Q. Minh, IQ-TREE: A fast and effective stochastic algorithm for estimating maximum-likelihood phylogenies. Mol. Biol. Evol. 32, 268–274 (2015).

50. S. Kalyaanamoorthy, B. Q. Minh, T. K. F. Wong, A. von Haeseler, L. S. Jermiin, ModelFinder: Fast model selection for accurate phylogenetic estimates. Nat. Methods 14, 587–589 (2017).

51. M. T. Islam et al., Emergence of wheat blast in Bangladesh was caused by a South American lineage of *Magnaporthe oryzae*. BMC Biol. 14, 84 (2016).

52. Z. Peng et al., Effector gene reshuffling involves dispensable mini-chromosomes in the wheat blast fungus. PLoS Genet. 15, e1008272 (2019).

53. S. Asuke, A. Horie, K. Komatsu, R. Mori, T. T. P. Vy, Y. Inoue, Y. Jiang, Y. Tatematsu, M. Shimizu, Y. Tosa, Loss of PWT7, Located on a Supernumerary Chromosome, Is Associated with Parasitic Specialization of *Pyricularia oryzae* on Wheat. Mol. Plant-Microbe Interact. 36, 716–725 (2023).

54. Y. Inoue et al., Evolution of the wheat blast fungus through functional losses in a host specificity determinant. Science 357, 80–83 (2017).

55. V. Were et al., Genome sequences of sixty *Magnaporthe oryzae* isolates from multiple host plant species. Zenodo (2021); 10.5281/zenodo.4627042.

56. B. Tembo et al., Whole genome shotgun sequences of *Magnaporthe oryzae* wheat blast isolates from Zambia. Zenodo (2021); 10.5281/zenodo.4637174.

57. H. Li, R. Durbin, Fast and accurate short read alignment with Burrows–Wheeler transform. Bioinformatics 25, 1754–1760 (2009).

58. E. M. Ortiz, vcf2phylip v2.0: Convert a VCF matrix into several matrix formats for phylogenetic analysis. Zenodo (2019).

59. B. Q. Minh et al., IQ-TREE 2: New models and efficient methods for phylogenetic inference in the genomic era. Mol. Biol. Evol. 37, 1530–1534 (2020).

60. P. O. Lewis, A likelihood approach to estimating phylogeny from discrete morphological character data. Syst. Biol. 50, 913–925 (2001).

61. D. T. Hoang, O. Chernomor, A. von Haeseler, B. Q. Minh, L. S. Vinh, UFBoot2: Improving the ultrafast bootstrap approximation. Mol. Biol. Evol. 35, 518–522 (2018).

62. S. Guindon, J.-F. Dufayard, V. Lefort, M. Anisimova, W. Hordijk, O. Gascuel, New algorithms and methods to estimate maximum-likelihood phylogenies: Assessing the performance of PhyML 3.0. Syst. Biol. 59, 307–321 (2010).

63. J. Win, E. Chanclud, C. S. Reyes-Avila, T. Langner, T. Islam, S. Kamoun, Nanopore sequencing of genomic DNA from *Magnaporthe oryzae* isolates from different hosts. Zenodo (2019); 10.5281/zenodo.2564950.

64. C. Camacho, et al., BLAST+: Architecture and applications. BMC Bioinformatics 10, 421 (2009).

65. E. Weber, C. Engler, R. Gruetzner, S. Werner, S. Marillonnet, A modular cloning system for standardized assembly of multigene constructs. PLoS ONE 6, e16765 (2011).

66. C. Engler et al., A Golden Gate modular cloning toolbox for plants. ACS Synth. Biol. 3, 839– 843 (2014).

67. K. Honda, T. Fujiwara, H. Adachi, Blue Native-PAGE Assay of NLR Protein Complexes Combined with Genetic Complementation in Nicotiana benthamiana. In: Plant Immunity. Methods in Molecular Biology, vol. 3012, pp. 129–141 (2026).

